# AGPAT2 Inflammatory Metabolic Reprogramming Mediates Immune Checkpoint Inhibitor-associated Myocarditis via LPAR5-pyroptosis Axis

**DOI:** 10.64898/2026.11.11.766056

**Authors:** Yifan Chen, Yihui Shen, Hui Zhang, Xuejun Wang, Yuchen Xu, Jian Zhang, Xiaozhen He, Mengxiao Han, Leilei Cheng, Junbo Ge

## Abstract

**Objectives:** Immune checkpoint inhibitors (ICIs), especially programmed cell death protein-1 (PD1) inhibitors, are effective onco-therapeutic agents. However, ICI-associated myocarditis (ICIAM) greatly limits their clinical utility. Imbalance between 1-acylglycerol-3-phosphate O-acyltransferases (AGPAT2) and lysophosphatidic acid (LPA) induced by inflammatory metabolic reprogramming (IMRP) might cause cardiovascular injury. This study aims to explore the link between AGPAT2 pathway and ICIAM as well as identifies diagnosis and treatment targets of ICIAM.

**Methods:** ICIAM model was established by injecting anti-PD1 (αPD1) in wild-type (WT) and AGPAT2 heterozygous knockout (AGPAT2^+/-^) mice. The pathogenic and regulating mechanisms of AGPAT2-LPA and its downstream signaling were examined in vitro and in vivo. A tumor-patient cohort was enrolled to observe the association between serum LPA and risk of ICIAM.

**Results:** In the αPD1-induced ICIAM model, myocardial IMRP occurred due to upstream immunity disorder, leading to downstream AGPAT2 degradation via lysosomal pathway and subsequent LPA accumulation. Knockout of AGPAT2 gene also promoted LPA accumulation. The accumulated LPA, by binding to its dominant receptor subtype 5 (LPAR5), triggered pyroptotic myocardial damage and resultant ICIAM. LPAR5 knockdown with siRNA or inhibition with TCLPA5-4 played the most significant rescue effect on myocardial pyroptosis, while stimulating LPAR5 with overdose LPA significantly increased myocardial pyroptosis. In ICI-treated patients, elevated serum LPA was associated with increased risk of ICIAM.

**Conclusions:** IMRP-triggered dysfunction of AGPAT2-LPA-LPAR5-pyroptosis axis is involved in pathogenesis of ICIAM. Several elements in this axis may be the promising targets in diagnosis (LPA as a biomarker) and treatment (LPA or LPAR5 as therapeutics) of ICIAM.

## Introduction

Immune checkpoint inhibitors (ICIs), such as programmed cell death protein-1 (PD1) inhibitors, have become an effective anti-tumor therapy. Unfortunately, over 70% of patients treated with ICIs experienced immune-related adverse events (irAEs) in the heart, lung, liver, intestine and skin(**1**), among which immune checkpoint inhibitor-associated myocarditis (ICIAM) is infrequent but lethal with mortality of 36-67%(**2, 3**), greatly restricting clinical application of ICIs. Hence, early diagnose or effective treatment of ICIAM has been a clinical priority in cardio-oncology.

The algorithm for early diagnosis of ICIAM has been recommended in current guidelines but the efficacies of treatment regimens, mostly using immunosuppressors, remain uncertain(**4–7**). For instance, glucocorticoid, though commonly used clinically, may offset the anti-tumor potency of ICIs with side-effects(**8**), highlighting the need of further exploration to find new diagnostic markers or therapeutic targets.

There are several proposed pathogeneses of ICIAM, including T-cell activation and misidentification(**9–11**), autoantigen-antibody formation(**12**) and cardiac immune protection failure(**12–14**) etc., most of them mainly address the very upstream of immunity disorders, thus leading to paradoxical use of immunosuppressants to treat ICIAM and resultant controversial outcomes(**11, 15, 16**).

Recently, the concept of inflammatory metabolic reprogramming (IMRP) has been adopted in oncologic studies. Early theories in tumor immunology defined IMRP as the dynamic alterations in the chain of inflammatory metabolism that was regulated by the adaption of immune cell metabolism after inflammation stimulations(**17, 18**).

This hypothesis can also be applied to explain the underlying mechanism through which anti-tumor medications disturb the immune homeostasis of myocardium, ultimately resulting in cardiomyocyte injury(**19–23**). Obviously, IMRP hypothesis, focusing on more downstream pathophysiology of ICIAM, may enable us to search new diagnostic markers or therapeutic targets.

Based on IMRP theory, we conducted a pre-study for screening potential targets with Gene Ontology (GO) and Kyoto Encyclopedia of Genes and Genomes (KEGG) analyses, identifying some clues of 1-acylglycerol-3-phosphate o-acyltransferase 2 (AGPAT2) downregulation, lysophosphatidic acid (LPA) elevation and cardiomyocyte pyroptosis both in patients and mice with ICIAM. These promising hints merit further studies to determine whether AGPAT2-LPA-pyroptosis axis involves in pathogenesis of ICIAM.

## Results

### Establishment of a Mouse ICIAM Model Mimicking Clinical Hallmarks

By intraperitoneal injection of anti-PD1 (αPD1) in the context of immunization with troponin (Figure 1A), we created a wild type (WT) mouse ICIAM model characterized by major CD44^+^CD8^+^T and minor CD4^+^T infiltrating sporadically in the left ventricular subepicardium and sheet-like in the pericardium accompanied by patchy myocardial damage without noticeable fibrosis (Figure 1B-I), elevation of myocardial injury biomarkers (cTnT and CK-MB) and cell-released substances (IL-1β/18, HMGB1, LDH) (Figure 1J-O), decrease of ATP level (Figure 1P) and reduction of the left ventricular ejection fraction (LVEF) and left ventricular fraction shortening (LVFS) (Figure 1Q-R), along with poor general condition such as loss of body weight (Supplemental Figure 1A), changes in heart weight and ventricular wall thickness (Supplemental Figure 1B-H), all of which recapitulates the hallmarks of ICIAM patients, thus laying the fundamental for further study. Based on the diagnostic criteria for ICIAM(**24**), the modelling success rate in our study was 83% with animal survival rate of 95%.

**Figure 1.**
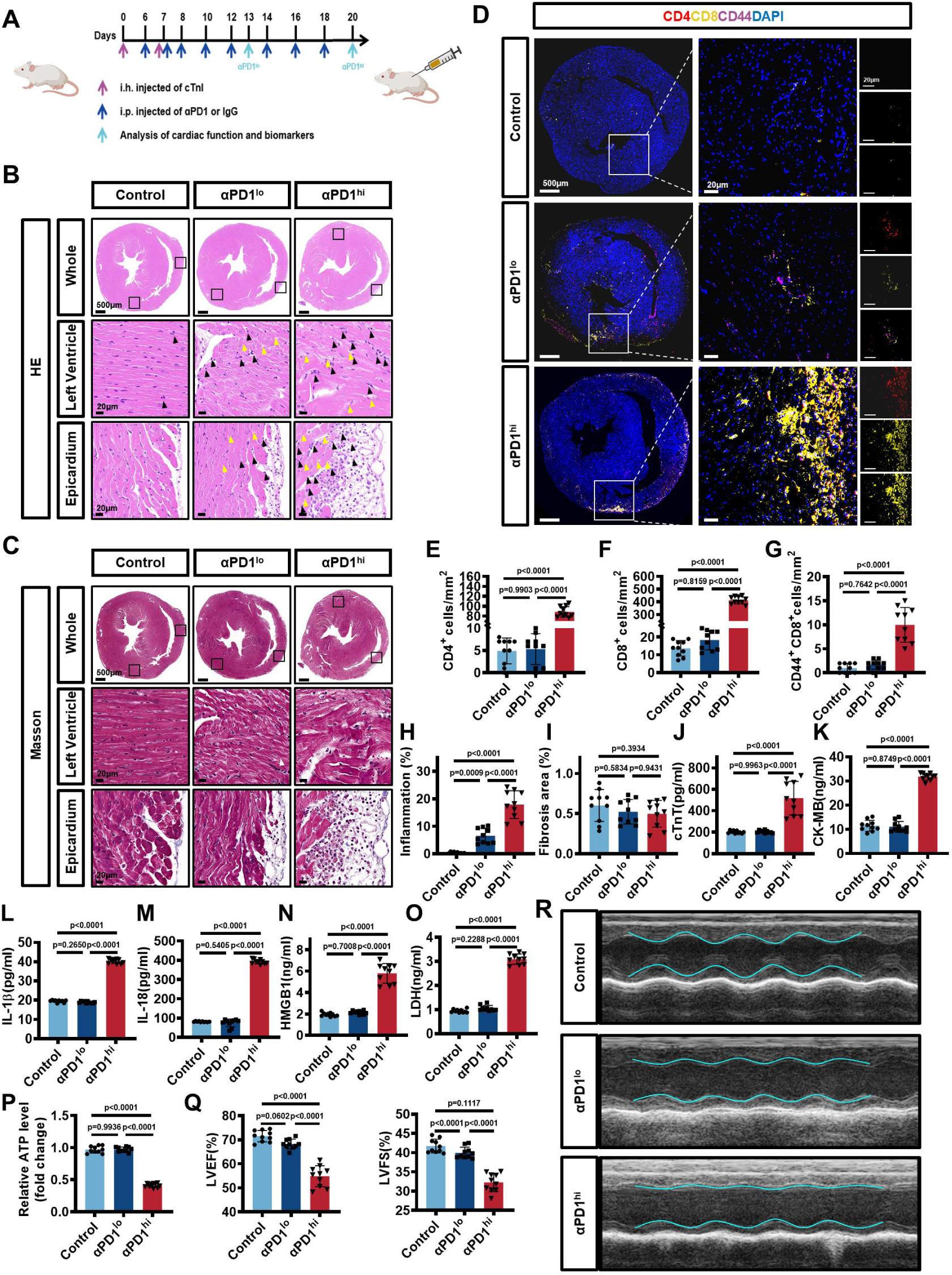
A Novel ICIAM Mouse Model Recapitulating Clinical Characteristics of Patients. A. Schematizing a new approach to ICIAM mouse modelling: in the context of immunization with cTnI, 5 doses (low dosage) or 8 doses (high dosage) of αPD1 (10mg/kg/dose) were sequentially injected intraperitoneally to construct the mouse models of αPD1lo group or αPD1hi group, respectively. The mice with only injecting IgG served as the Controls (control group). B-C, H-I. Representative HE (B) and Masson’s trichrome staining (C) and quantification of inflammation infiltration (H) and fibrosis area (I) in each group (n = 10/group, one-way ANOVA with post-hoc Tukey HSD test). The left ventricle myocardium and right ventricle epicardium in each group are shown respectively. The immune cells in each group are marked with black arrows, and the disordered cardiomyocytes are marked with yellow arrows. D-G. Representative immunofluorescence staining (D) and quantification of CD4+ cells (E), CD8+ cells (F) and CD44+CD8+ double-positive cells (G), with staining CD4 in red, CD8 in yellow, CD44 in magenta and nuclei in blue in each group (n = 10/group, one-way ANOVA with post-hoc Tukey HSD test). Scale bar, 500μm or 20μm. J-P. The serum cTnT, CK-MB, IL-1β, IL-18, HMGB1 and LDH levels were measured by ELISA assays, and the ATP level in the heart was measured by ATP assay in each group (n = 10/group, one-way ANOVA with post-hoc Tukey HSD test). Q-R. Representative M-mode echocardiographic images (R) and quantification of LVEF and LVFS (Q) in each group (n = 10/group, one-way ANOVA with post-hoc Tukey HSD test). Data were presented as mean ± standard deviation (SD) (≥3 experiments). (The same requirement for data expression unless specifically indicated in the following figures). αPD1, anti-PD1; ATP, adenosine triphosphate; CK-MB, creatine kinase-myocardial band; cTnI, cardiac troponin I; cTnT, cardiac troponin T; HMGB1, High-mobility group box 1; i.h., hypodermic injection; i.p., intraperitoneal injection; IgG, immunoglobulin G; IL-1β, interleukin-1 beta; IL-18, Interleukin-18; LDH, lactate dehydrogenase; LVEF, left ventricular ejection fraction; LVFS, left ventricular fractional shortening.

### Multi-omic Identification of Potential Pathogenic Clues of ICIAM

Multi-omics enrichments and signaling pathways investigation were conducted with metabolomics and proteomics analyses of patients and mice with ICIAM (Figure 2A-C) for explore potential pathogenic clues of ICIAM with referencing the human single-cell database.

**Figure 2.**
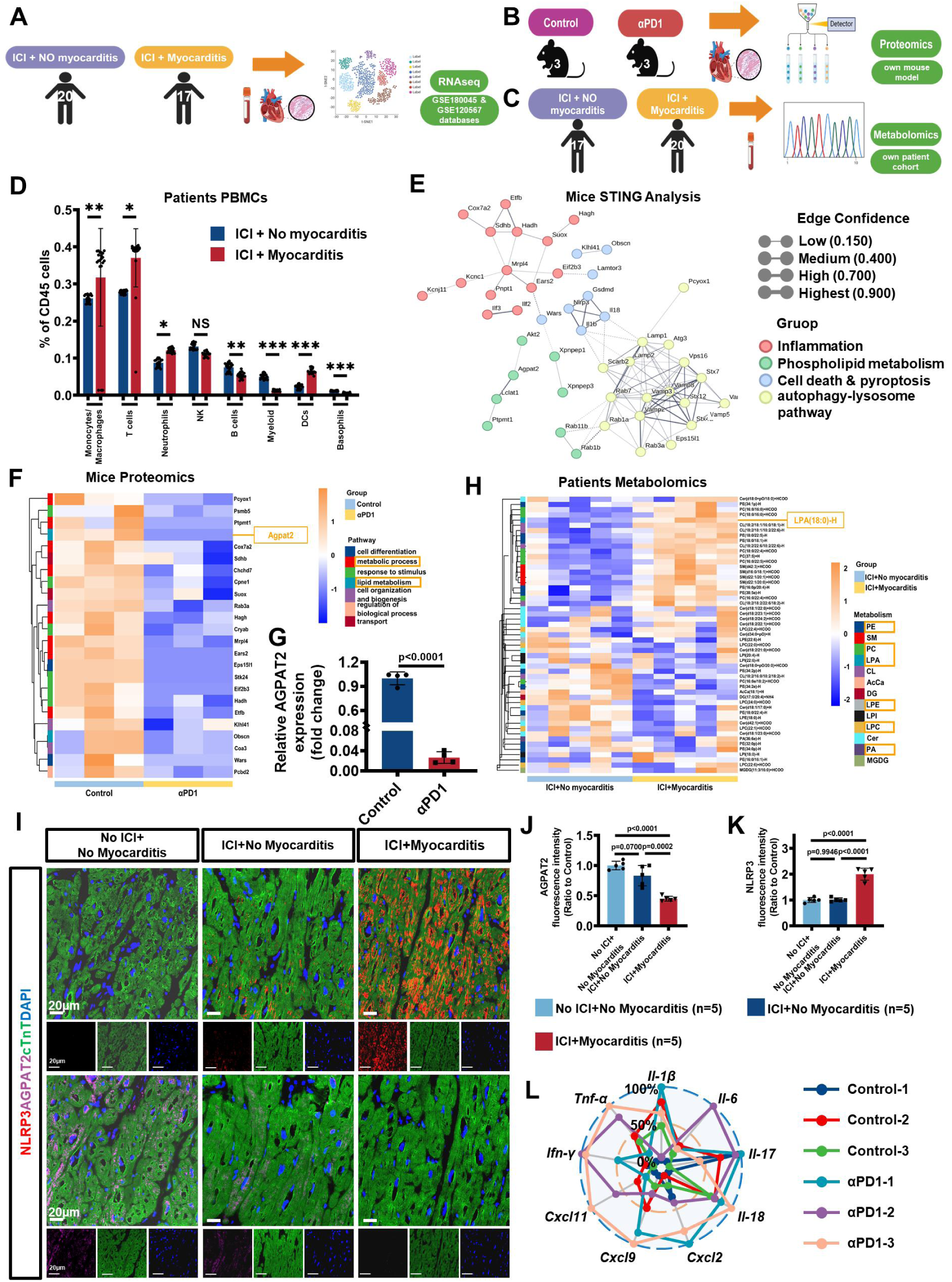
Omics Reveals AGPAT2-IMRP Link in ICIAM Patients and Mice. A. RNAseq analysis of tumor patients with ICI+No myocarditis or ICI+Myocarditis from online databases (GSE180045 and GSE120567). B-C. Proteomics analysis of our own mice in Control and αPD1groups (B), and metabolomics analysis of our own tumor patient cohort with ICI+No myocarditis or ICI+Myocarditis (C). D. Single cell analysis and quantification of different immune cells subgroups (n = 20 for ICI+No myocarditis, n = 17 for ICI+Myocarditis, unpaired two-tailed Student’s t-test). E. STING analysis revealed major differential proteins, pathways and their relationships in Control and ICIAM mice (αPD1 groups). F-G. The clustering heatmap and quantification showing the most significant enrichment differential proteins in Control and ICIAM mice. H. Clustering heatmaps of most significant enrichment differential lipid metabolites in tumor patients with ICI+No myocarditis or ICI+Myocarditis (n = 5/group). I-K. Representative immunofluorescence staining (I) and quantification of AGPAT2 (J), NLRP3 (K) in myocardial biopsy sections from tumor patients with No ICI+No myocarditis, ICI+No myocarditis and ICI+Myocarditis, with staining NLRP3 in red, AGPAT2 in magenta, cTnT in green and nuclei in blue (n = 5/group, one-way ANOVA with post-hoc Tukey HSD test). Scale bar, 20μm. L. Transcription levels of *Il-1β、Il-6、Il-17、Il-18、Tnf-α、Ifn-γ、Cxcl2、Cxcl9、Cxcl11* in Control and ICIAM mice (n = 3/group). Data were presented as mean ± SD. αPD1, anti-PD1; AGPAT2, 1-acylglycerol-3-phosphate O-acyltransferase 2; Cxcl2, chemokine (C-X-C motif) ligand 2; Cxcl9, chemokine (C-X-C motif) ligand 9; Cxcl11, chemokine (C-X-C motif) ligand 11; cTnT, cardiac troponin T; ICI, immune checkpoint inhibitors; ICIAM, immune checkpoint inhibitor-associated myocarditis; Ifn-γ, interferon gamma; IMRP, inflammatory metabolic reprograming; Il-1β, interleukin-1 beta; Il-6, interleukin-6; Il-17, Interleukin-17; Il-18, Interleukin-18; NLRP3, NOD-like receptor thermal protein domain-associated protein 3; PBMC, peripheral blood monocyte; Tnf-α, tumor necrosis factor alpha.

Single-cell RNA sequencing (scRNAseq) data (GSE180045) of peripheral blood mononuclear cell was combined with the myocardial biopsy RNA sequencing data (GSE120567) of ICIAM patients to construct an immune cell atlas (Supplemental Figure 2A). Using the singleR algorithm, eight subtypes of immunocytes were annotated (Figure 2D, Supplemental Figure 2B), seven of them exhibited significant proportion differences between the ICI+No myocarditis and ICI+Myocarditis groups (Figure 2D), implying significant immunity disorder in the setting of ICIAM. Moreover, gene network also revealed ICIAM-related dysregulation in the phospholipid metabolism (Supplemental Figure 2C).

When KEGG and GO analyses were performed on mice model, the αPD1 group exhibited activation of signaling pathways related to inflammation, metabolism, cell death and autophagy-lysosome (Supplemental Figure 2D). Protein-protein interaction network(**25**) categorized the altered proteins into myocardial inflammation, phospholipid metabolism, lysosome, and pyroptosis pathways (Figure 2E).

Above findings imply that dysimmunity, inflammation, phospholipid metabolic disorder and pyroptosis may involve in ICIAM pathogenesis, of which phospholipid metabolic disorder may be crucial. Actually, in ICIAM patients, both clustering heatmaps (Figure 2H) and metabolite volcano plots (Supplemental Figure 2E) showed substantially increased serum levels of phosphatidic acid metabolites, particularly LPA. In ICIAM mice, αPD1 markedly reduced the level of AGPAT2 (Figure 2F-G), a key metabolic enzyme for LPA degradation.

Altogether, these results hint that AGPAT2-IMRP, especially AGPAT2-LPA imbalance may be a potential pathway in ICIAM pathogenesis, worthy of further exploration.

### Involvement of Myocardial AGPAT2-IMRP in Patients or Mice with ICIAM

In ICIAM patients (biopsy) or mice, myocardial tissues with immunofluorescent co-staining (IFCS) revealed that the levels of NLRP3, cleaved-Caspase 1, Granzyme B and GSDMD were significantly increased and AGPAT2 expression was decreased (Figure 2I-K, Supplemental Figure 2F, H); CD4, CD8, CD44CD8 positive signals demonstrated increased inflammatory cells infiltration in the myocardium (Supplemental Figure 2G, I) ; serum enzyme-linked immunoassay (ELISA) showed that myocardial injury markers of cTnT and CK-MB (Supplemental Figure 2J-K), pyroptosis-related cell-released substances of IL-1β, IL-18, HMGB1, LDH (Supplemental Figure 2L-O) were elevated while ATP level was decreased (Supplemental Figure 2P) with LPA accumulation in ICIAM patient (Supplemental Figure 2Q). Additionally, in ICIAM mice, transcription of pro-inflammatory and chemotactic cytokines was markedly increased (Figure 2L); and expression of AGPAT2 protein was decreased significantly with increasing doses of αPD1 (Supplemental Figure 2R-S), accompanied by serum LPA elevation (Supplemental Figure 2T).

These results support AGPAT2-IMRP (immunity-initiated AGPAT2 degradation with LPA accumulation) involves in the pathogenesis of ICIAM in patients and mice. Yet, the intracellular mechanisms of AGPAT2-LPA imbalance, subsequent myocardial injury and resultant ICIAM remain elusive.

### Degradation of AGPAT2 Mainly via Lysosome Pathway

To explore the mechanisms underlying AGPAT2-LPA dysregulation, the KEGG clustering bubble chart was performed. The results pointed to the involvement of vesicle metabolism, endocytosis, and various lysosome-related degradation pathways, suggesting that the AGPAT2 reduction may be mediated by lysosome pathway (Supplemental Figure 3A).

In the vitro experiment with HL-1 cells stimulated by LPS+Nig in presence or absence of lysosome inhibitor (BafA1) or proteasome inhibitor (MG132) showed that BafA1 significantly reduced AGPAT2 degradation in pyroptotic cells (Figure 3A, B). Immunoprecipitation-mass spectrometry (IP-MS) examined potential main proteins interacted with AGPAT2, finding lysosomal-associated membrane protein 1 (LAMP1) was the most relevant protein (Supplemental Figure 3B, C). Endogenous co-immunoprecipitation (Co-IP) analysis confirmed their interaction (Figure 3C, D) and this finding was further confirmed by 2D and 3D immunofluorescence co-staining (Figure 3E, H, I, Supplemental Figure 3D).

**Figure 3.**
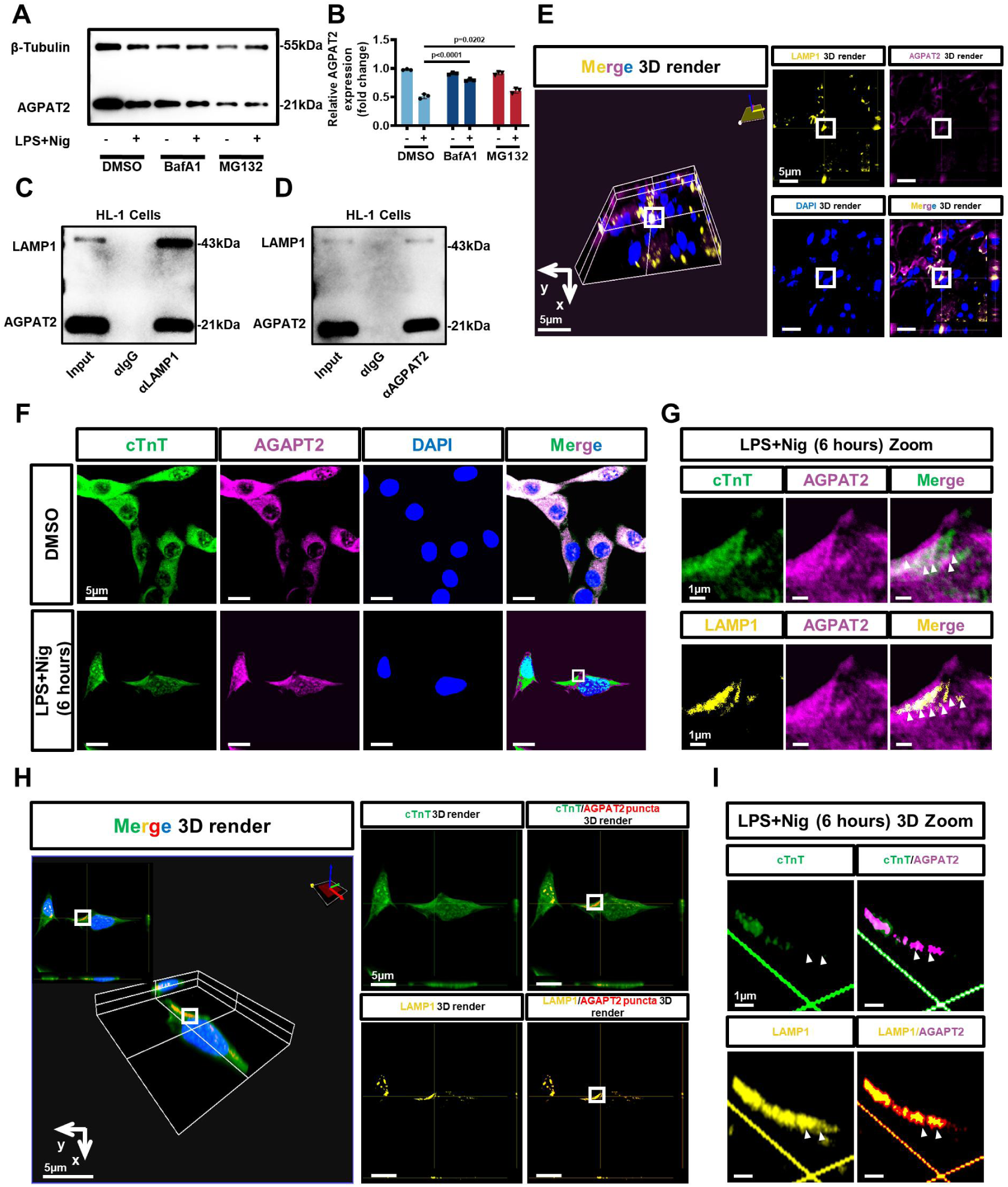
AGPAT2 Degradation upon Inflammation Stimulation via Lysosomal Metabolic Reprogramming. A-B. Representative bands (A) and quantification (B) of AGPAT2 protein from HL-1 mouse cardiomyocytes subjected to LPS+Nig stimulation for 6 hours and intervened by different inhibitors. (n = 3/group, two-way ANOVA with post-hoc Tukey HSD test). C-D. Endogenous Co-IP validated the interaction between AGPAT2 and LAMP1 proteins in HL-1 cardiomyocytes. E. Representative 3D immunofluorescence colocalization images (marked with square) of mice hearts subjected to αPD1 treatment, with staining LAMP1 in yellow, AGPAT2 in magenta and nuclei in blue (n = 4/group). Scale bar, 5μm. F. Representative immunofluorescence images of HL-1 cardiomyocytes stimulated with or without LPS+Nig for 6 hours with staining cTnT in green, AGPAT2 in magenta and nuclei in blue (colocalization marked with square). Scale bar, 5μm. G. Magnified region from (F) highlighted sequestration of AGPAT2 signal into puncta, distinct from cTnT signal and enclosed in a LAMP1-positive compartment (marked with arrow), with staining cTnT in green, AGPAT2 in magenta, LAMP1 in yellow and nuclei in blue. Scale bar, 1μm. H. Representative three-dimensional immunofluorescence colocalization images of HL-1 cardiomyocytes stimulated with LPS+Nig for 6 hours, highlighted AGPAT2 puncta localization within LAMP1 compartment (marked with square), with staining cTnT in green, AGPAT2 puncta in red, LAMP1 in yellow and nuclei in blue. Scale bar, 5 μm. I. Magnified 3D regions from (H) highlighted AGPAT2 puncta localization outside the cTnT signal and colocalized within LAMP1-positive spherical structures (marked with arrow), with staining cTnT in green, AGPAT2 in magenta, LAMP1 in yellow. Scale bar, 1μm. Data were presented as mean ± SD. AGPAT2, 1-acylglycerol-3-phosphate O-acyltransferase 2; αIgG, immunoglobulin G antibody; BafA1, Bafilomycin A1, lysosomal inhibitors; cTnT, cardiac troponin T; Co-IP, co-immunoprecipitation; DMSO, Dimethyl sulfoxide; LAMP1, lysosomal-associated membrane protein 1; LPS, lipopolysaccharide; MG132, Carbobenzoxy-L-Leucyl-L-Leucyl-L-Leucinal, proteasome inhibitor; Nig, nigericin.

Co-localization of AGPAT2 and LAMP1 in myocardia showed selective AGPAT2 delivery to endolysosomal compartments (Figure 3E, Supplemental Figure 3D). In HL-1 cells stimulated with LPS+<u>Nig</u>, there were two subsets’ expressions of AGPAT2 puncta (Figure 3F-I, Supplemental Figure 3E, F): one sequestrated into spherical LAMP1 structures was upregulated, the other retained within the myocardial cells was downregulated.

Above results suggest that lysosome is the major pathway mediating metabolic reprogramming and AGPAT2 degradation in cardiomyocytes post inflammatory reprogramming stimulation.

### Impact of AGPAT2 Deficiency on Myocardial Pyroptosis and Injury

Above bioinformatic analyses found that pyroptosis was involved in ICIAM pathogenesis (Figure 2E). Subsequently, the impact of AGPAT2 downregulation on myocardial injury was examined. WT mice received low or high dose of αPD1 (αPD1^lo^ or αPD1^hi^) (Figure 1A), adult mouse left ventricular primary cardiomyocytes were isolated after modeling, AGPAT2 protein expression began to decrease at αPD1^lo^ and decreased more prominently at αPD1^hi^ (Supplemental Figure 4A-B), dose-dependent upregulation was observed in pyroptosis-related proteins (Supplemental Figure 4A, C-G). Myocardial injury markers (Figure 1J-K), pyroptosis-related cell-released substances and LPA levels (Figure 1L-P, Supplemental Figure 2T) were significantly elevated upon high dose of αPD1 stimulation. Additionally, myocardial immunofluorescence staining of AGPAT2, and pyroptosis markers like NLRP3, cleaved Caspases-1, Granzyme B and GSDMD also revealed corresponding alterations (Supplemental Figure 4H-M).

To verify the mechanistic link, heterozygous AGPAT2 gene knockout (AGPAT2^+/-^) mice (Supplemental Figure 9A, D) were generated and ICIAM model was established in the AGPAT2^+/-^ mice (Figure 4A, B). The AGPAT2^+/-^ mice exhibited inherent low-level transcription and protein expression of AGPAT2 (Figure 4C,O, Supplemental Figure 9C), allowing more insight into the impact of αPD1 on myocardial pyroptosis in the context of AGPAT2 deficiency.

**Figure 4.**
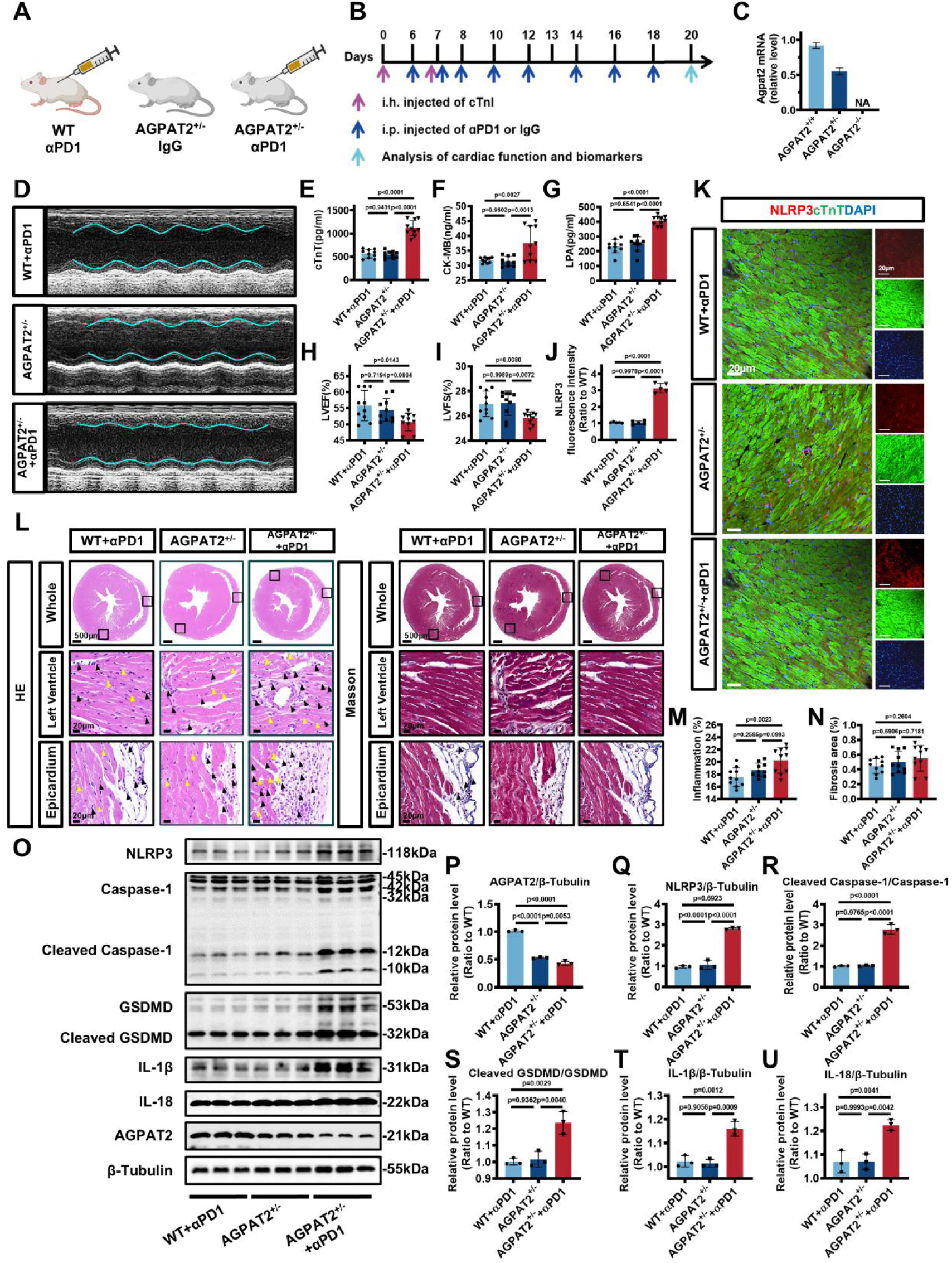
AGPAT2 Downregulation Exacerbates Myocardial Pyroptosis in ICIAM Mice. A-B. Experimental scheme depicted wild-type (WT) and heterozygous AGPAT2 knockout (AGPAT2^+/-^) mice with or without ICIAM mouse modelling method. C. The mRNA expression of mice with different genotypes of AGPAT2. D, H-I. Representative M-mode echocardiographic images (D) and quantification of LVEF (H) and LVFS (I) in each group (n = 10/group, one-way ANOVA with post-hoc Tukey HSD test). E-G. The serum cTnT (E), CK-MB (F) and LPA (G) levels measured by ELISA assay (n = 10/group, one-way ANOVA with post-hoc Tukey HSD test). L-N. Representative HE and Masson’s trichrome staining (L) and quantification of inflammation infiltration (M) and fibrosis area (N) in each group (n = 10/group, one-way ANOVA with post-hoc Tukey HSD test). The left ventricle myocardium and right ventricle epicardium in each group are shown respectively. The immune cells in each group are marked with black arrows, and the disordered cardiomyocytes are marked with yellow arrows. J-K. Representative immunofluorescence images (K) and quantification (J) in each group, with staining NLRP3 in red, cTnT in green and nuclei in blue (n = 10/group, one-way ANOVA with post-hoc Tukey HSD test). Scale bar, 20μm. O-U. Representative bands (O) and quantification (P-U) of AGPAT2 and pyroptosis pathway-associated proteins in each group (n = 10/group, one-way ANOVA with post-hoc Tukey HSD test). Data were presented as mean ± SD. AGPAT2, 1-acylglycerol-3-phosphate O-acyltransferase 2; AGPAT2^+/-^, heterozygous AGPAT2 knockout; AGPAT2^-/-^, homozygote AGPAT2 knockout; αPD1, anti-PD1 (PD1 inhibitor); Caspase-1, Cysteine-aspartic proteases 1; CK-MB, creatine kinase-myocardial band; cleaved Gasdermin D, N-terminal gasdermin domain; cTnT, cardiac troponin T; GSDMD, Gasdermin D; HE staining, Hematoxylin and Eosin staining; ICIAM, immune checkpoint inhibitor-associated myocarditis; IL-1β, Interleukin-1β; IL-18, Interleukin-18; LPA, lysophosphatidic acid; LVEF, left ventricular ejection fraction; LVFS, left ventricular fractional shortening; NLRP3, NOD-like receptor pyrin domain-containing protein 3; WT, wild-type.

Compared to the WT ICIAM models and AGPAT2^+/-^ mice in the absence of αPD1, the AGPAT2^+/-^ ICIAM model mice showed more reduction of body weight, higher ratio of heart weight to body weight without differences in heart weight or the ratio of heart weight to tibia length (Supplemental Figure 4N); had more increase of cTnT and CK-MB, more elevated cell-released substances and more accumulation of LPA (Figure 4E-G, Supplemental Figure 4O-S) and more decrease of LVEF and LVFS (Figure 4D, H-I), and had more infiltration of immunocyte (Figure 4L, M), more upregulation of NLRP3 and other pyroptosis-related proteins like cleaved Caspases-1, Granzyme B, and the GSDMD (Figure 4J, K, Supplemental Figure 4T-Y) without fibrosis progression in myocardium (Figure 4L, N). Meanwhile, AGPAT2 protein levels were reduced (Figure 4O, P) in adult mouse left ventricular primary cardiomyocytes, and expressions of pyroptosis-related proteins were significantly upregulated (Figure 4O, Q-U).

Altogether, myocardial pyroptosis and injury can be modulated by AGPAT2 downregulation, exacerbated by AGPAT2 deficiency.

### Myocardial Injury by LPA is Mainly Mediated via Pyroptotic Pathway

The above experiments have revealed that myocardial injury in ICIAM can be caused by myocardial pyroptosis. To test if other cardiomyocyte death pathways (i.e., necroptosis, apoptosis) are involved in pathogenesis of ICIAM or not, we first determined an optimal LPA concentration of 20μM for stimulating cardiomyocytes based on in vivo studies and CCK-8 assays (Figure 5A). Subsequently, inhibitors targeting necroptosis (SPD304), apoptosis (Z-DEVD-FMK) and pyroptosis (INF39) were applied to LPA-stimulated HL-1 cells to examine the rescue effect on cell death. Notably, INF39 exhibited the most significant rescue effect on the cell viability (Figure 5B), indicating that LPA-caused cardiomyocyte death is mainly mediated by pyroptosis pathway.

**Figure 5.**
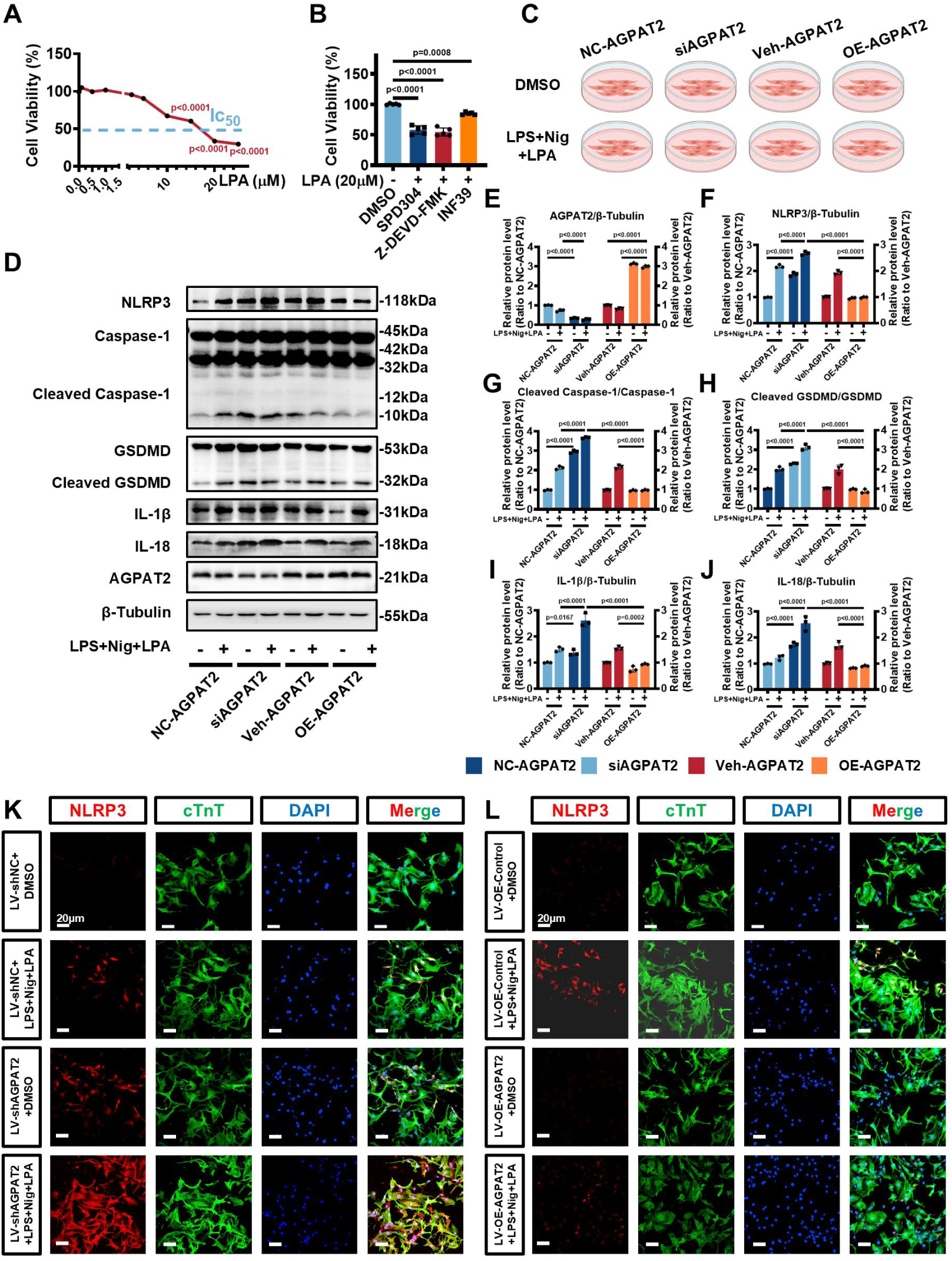
AGPAT2-LPA Imbalance Drives LPA-induced Pyroptosis in Cardiomyocytes. A. The Impact of different concentrations of LPA on HL-1 cell viability (n = 5/group, one-way ANOVA with post-hoc Tukey HSD test). B. The abilities of different inhibitors targeting necroptosis (SPD304), apoptosis (Z-DEVD-FMK), and pyroptosis (INF39) were evaluated for rescuing cell death (n = 5/group, one-way ANOVA with post-hoc Tukey HSD test). The HL-1 cells were pre-treated with 20μM LPA. C. Experimental scheme of silencing or overexpressing AGPAT2 in HL-1 cells subjected to LPS+Nig+LPA treatment. D-J. Representative bands (D) and quantification (E-J) of AGPAT2 and pyroptosis pathway-associated proteins from AGPAT2-silencing or overexpressing HL-1 cells subjected to LPS+Nig+LPA treatment (n = 3/group, two-way ANOVA with post-hoc Tukey HSD test). K-L. Representative immunofluorescence images of AGPAT2-silencing (LV-shAGPAT2) (K) or AGPAT2-overexpressing (LV-OE-AGPAT2) (L) in NRVCs subjected to LPS+Nig+LPA treatment, with staining NLRP3 in red, cTnT in green and nuclei in blue (n = 3/group). Scale bar, 20μm. Data were presented as mean ± SD. Caspase-1, Cysteine-aspartic proteases 1; cleaved Gasdermin D, N-terminal gasdermin domain; DMSO, Dimethyl sulfoxide; GSDMD, Gasdermin D; IL-1β, Interleukin-1β; IL-18, Interleukin-18; LPS, lipopolysaccharide; LPA, lysophosphatidic acid; LV-OE-AGPAT2, short hairpin-overexpression AGPAT2 virus; LV-OE-Control, short hairpin-overexpression control virus; LV-shAGPAT2, short hairpin-AGPAT2 virus; LV-shNC, short hairpin-negative control virus; NC-AGPAT2, negative control-AGPAT2 RNA; Nig, nigericin; NLRP3, NOD-like receptor pyrin domain-containing protein 3; NRVC, neonatal rat ventricular cardiomyocyte; OE-AGPAT2, overexpression-AGPAT2 plasmid; siAGPAT2, small interfering-AGPAT2 RNA; Veh-AGPAT2, vehicle-AGPAT2 plasmid.

### Effect of AGPAT2 Konckdown/Overexpression on Cardiomyocyte Pyroptosis

To examine the relationship between AGPAT2 and cardiomyocyte pyroptosis, and to further determine the role of pyroptosis pathway on the myocardial injury in ICIAM, rescue experiment was performed on HL-1 cells and primary neonatal rat ventricular cardiomyocytes (NRVCs) (Figure 5C).

In HL-1 cells, silencing AGPAT2 (Figure 5D, E) led to upregulation or activation in the pyroptosis-related proteins. The overall enhancement of pyroptosis pathway was further pronounced following stimulation with LPS+Nig+LPA (Figure 5D, F-J). Conversely, overexpressing AGPAT2 (Figure 5D, E) markedly attenuated cardiomyocyte pyroptosis triggered by the LPS+Nig+LPA stimulation (Figure 5D, F-J).

In NRVCs, pyroptosis level was detected by NLRP3 fluorescence intensity. Aligning with the outcomes in HL-1 cells, knockdown of AGPAT2 led to elevated NLRP3 fluorescence intensity, which was further enhanced by LPS+Nig+LPA stimulation (Figure 5K). In contrast, overexpressing AGPAT2 offered a noteworthy protective effect against pyroptosis (Figure 5L).

The above cellular studies demonstrate that regulating of AGPAT2 expression significantly affects the AGPAT2-LPA balance and AGPAT2-LPA dysregulation can modulate cardiomyocyte pyroptosis.

### Myocardial Pyroptotic Injury with LPAAccumulation

The role of LPA accumulation in ICIAM was further validated through animal experiments (Supplemental Figure 5A). After additionally intraperitoneal administration of LPA (1 mg/kg/day) to mimic LPA accumulation, the αPD1+LPA group exhibited the most pronounced reduction in LVEF and LVFS (Supplemental Figure 5B, C), accompanied by elevated levels of cTnT and CK-MB, pyroptosis-related cell-released substances (IL-1β, IL-18, HMGB1, LDH) and decreased ATP compared to the Control and αPD1 groups (Supplemental Figure 5D-J). WB analysis of left ventricular primary cardiomyocytes also revealed a marked downregulation of AGPAT2 in this group (Supplemental Figure 5K, L), concomitant with a broad upregulation of myocardial pyroptosis pathway (Supplemental Figure 5K, M-Q). Histopathological examination showed a more prominent infiltration of inflammation cells in myocardia in the αPD1+LPA group compared to the other groups (Supplemental Figure 5R, T). Meanwhile, immunofluorescence analysis indicated upregulated expression of NLRP3, cleaved Caspase-1, Granzyme B and GSDMD in αPD1+LPA group (Supplemental Figure 5S, U-X).

The above in vivo studies provide provides solid evidence that LPA accumulation exacerbates myocardial pyroptotic injury and ICIAM.

### Mediation of Cardiomyocyte Pyroptosis Dominantly via LPAR5

The subtypes of LPA receptors (LPARs), including LPAR1, LPAR2, LPAR3 and LPAR5, can involve in regulating immune homeostasis. To determine the dominant LPAR in mediation of LPA-induced myocardial pyroptosis in ICIAM, inhibitors and siRNA experiments were carried out (Figure 6A, B).

**Figure 6.**
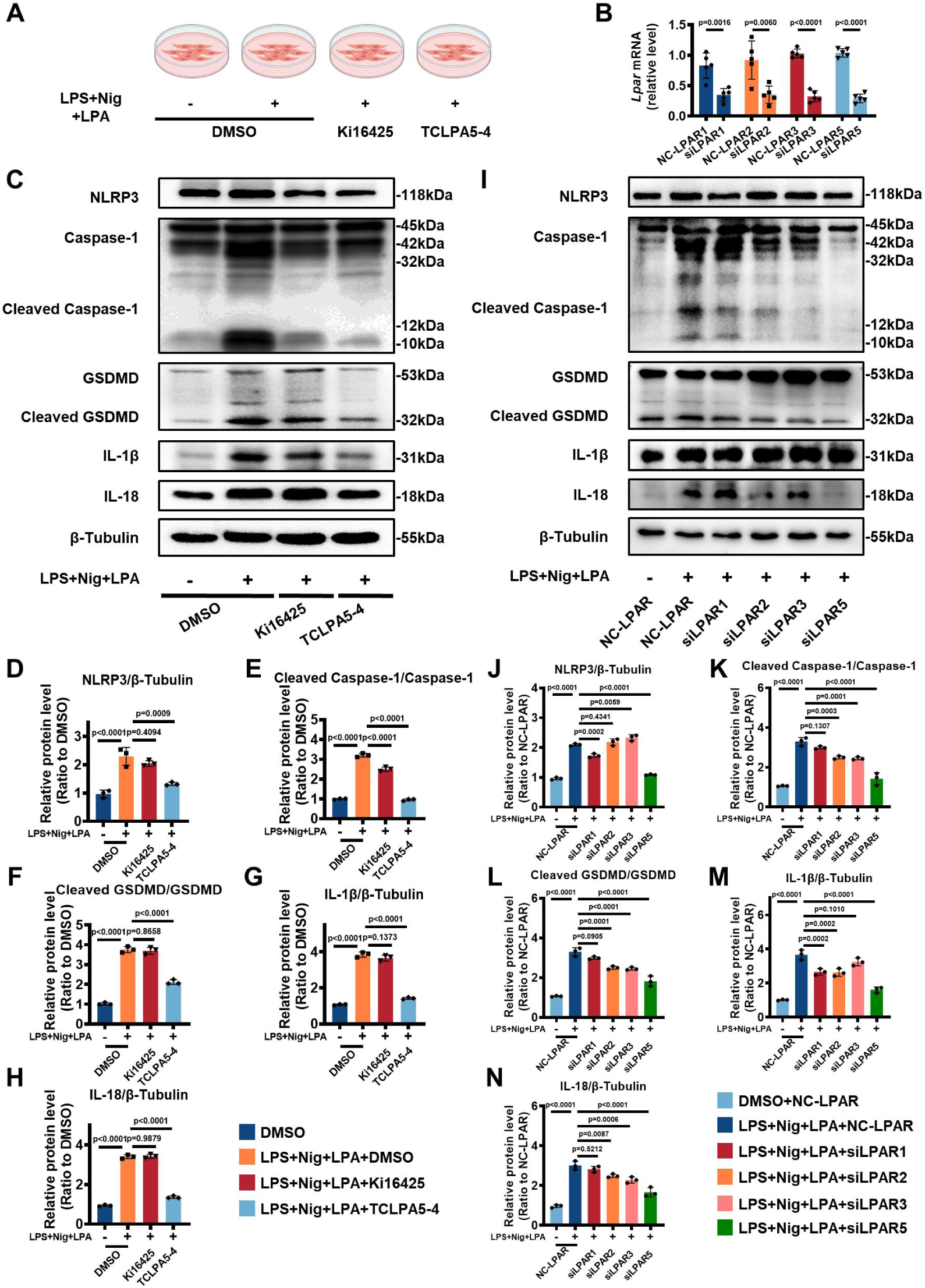
Roles of LPA Receptor 5 on Regulating of Cardiomyocyte Pyroptosis. Experimental scheme of pharmacological inhibition of different lysophosphatidic acid receptors (LPARs) in HL-1 cells treated with LPS+Nig+LPA. B. Knockdown efficiency of each siRNA on the transcription level of their corresponding LPARs (n = 5/group, unpaired two-tailed Student’s t-test). C-H. Representative bands (C) and quantification (D-H) of pyroptosis pathway-associated proteins from HL-1 cells, indicating the rescue effects of different LPAR pharmacological inhibitors on LPA-induced cardiomyocyte pyroptosis (n = 3/group, one-way ANOVA with post-hoc Tukey HSD test). I-N. Representative bands (I) and quantification (J-N) of pyroptosis pathway-associated proteins from HL-1 cells, indicating the rescue effects of siRNAs for different LPARs on LPA-induced cardiomyocytes pyroptosis (n = 3/group, one-way ANOVA with post-hoc Tukey HSD test). Data were presented as mean ± SD. Caspase-1, Cysteine-aspartic proteases 1; cleaved Gasdermin D, N-terminal gasdermin domain; DMSO, Dimethyl sulfoxide; GSDMD, Gasdermin D; IL-1β, Interleukin-1β; IL-18, Interleukin-18; Ki16425, inhibitor of LPAR1, LPAR2, and LPAR3; LPS, lipopolysaccharide; LPA, lysophosphatidic acid; LPAR, lysophosphatidic acid receptor; NC-LPAR1, LPA receptor 1 negative control siRNA; NC-LPAR2, LPA receptor 2 negative control siRNA; NC-LPAR3, LPA receptor 3 negative control siRNA; NC-LPAR5, LPA receptor 5 negative control siRNA; Nig, nigericin; NLRP3, NOD-like receptor pyrin domain-containing protein 3; siLPAR1, LPA receptor 1 siRNA; siLPAR2, LPA receptor 2 siRNA; siLPAR3, LPA receptor 3 siRNA; siLPAR5, LPA receptor 5 siRNA; TCLPA5-4, inhibitor of LPAR5.

Following intervention of LPS+Nig+LPA, WB indicated that Ki16425 could only reduce the cleaved Caspase-1/Caspase-1 ratio (Figure 6E) with a limited effectiveness on rescuing cell pyroptosis (Figure 6C-H). On the other hand, TCLPA5-4, even when exposed to LPS+Nig+LPA, continued to significantly counteract the elevating trend of cardiomyocyte pyroptosis (Figure 6C-H), providing initial evidence that LPAR5 may be a crucial regulatory receptor in LPA-induced cardiomyocyte pyroptosis.

The siRNA knockdown efficiency consistently ranged between 65% and 70% for each subtype of LPARs (Figure 6B). In this context, subsequent intervention with LPS+Nig+LPA revealed that siLPAR5 exhibited the most pronounced reversal effect on cardiomyocyte pyroptosis, with a significant downregulation of key pyroptotic proteins (Figure 6I-N). In contrast, the rescue effects of alternative siLPARs were found to be inferior to that of siLPAR5 (Figure 6I-N), reinforcing the pivotal role of LPAR5 as a critical regulatory node in LPA-induced cardiomyocyte pyroptosis.

### Mitigation of ICIAM by Blocking LPA-LPAR5-Apoptosis Axis

Following identifying LPAR5 as a dominant receptor involved in cardiomyocyte pyroptosis, the regulatory effect was further confirmed in the background of the AGPAT2^+/-^ mice (Figure 7A), WT mice (Supplemental Figure 7A) and melanoma-bearing mice (Supplemental Figure 8A).

**Figure 7.**
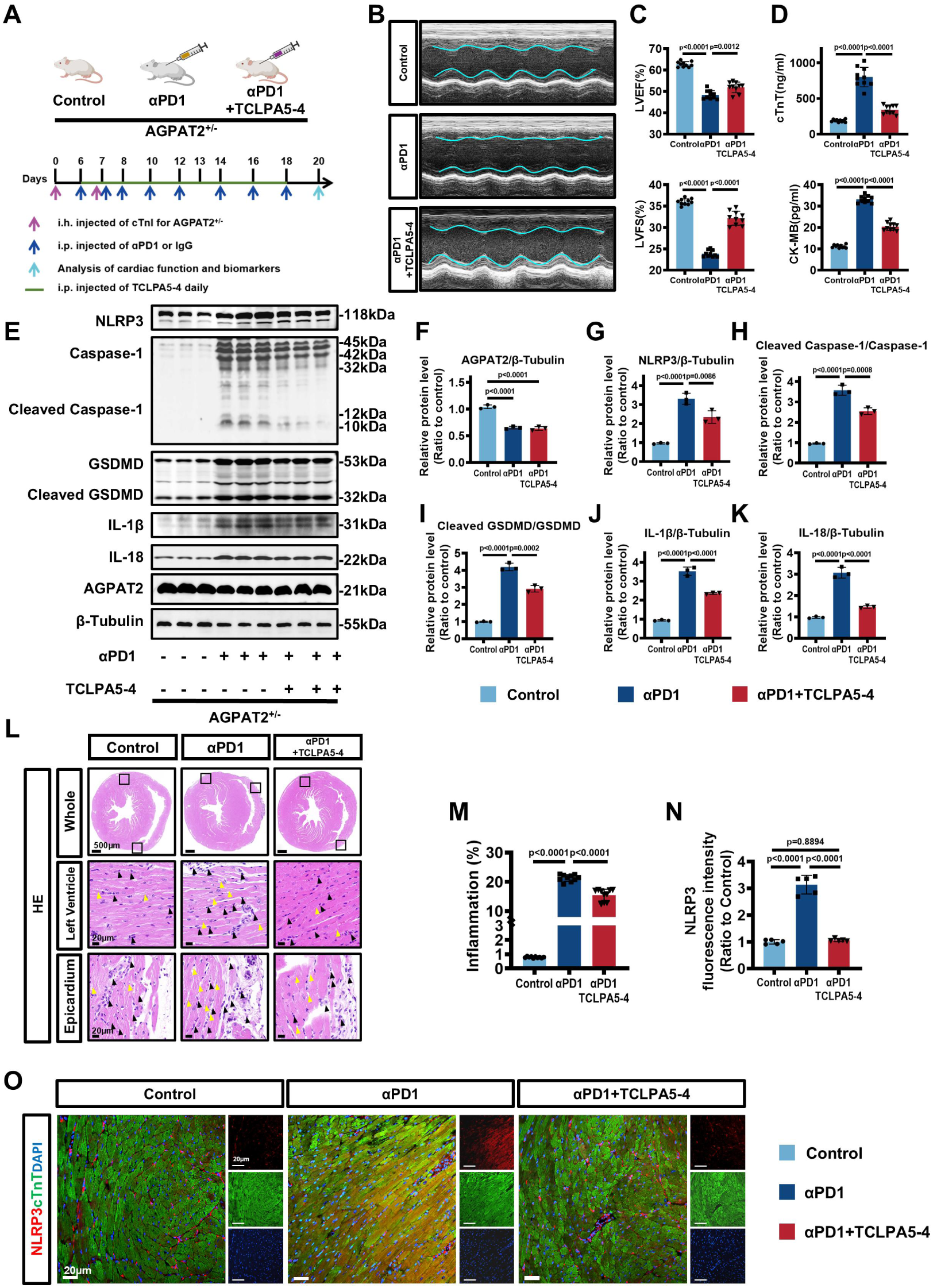
LPAR5 Blockade by TCLPA5-4 Reduces Pyroptosis in AGPAT2-deficient Mice. A. Experimental scheme depicted the effect of TCLPA5-4 in AGPAT2-deficiency (AGPAT2^+/-^) ICIAM mice. B-C. Representative M-mode echocardiographic images (B) and quantification of LVEF and LVFS (C) of AGPAT2^+/-^ ICIAM mice treated with or without TCLPA5-4 (n = 10/group, one-way ANOVA with post-hoc Tukey HSD test). D. The serum cTnT and CK-MB levels measured by ELISA assay in each group (n = 10/group, one-way ANOVA with post-hoc Tukey HSD test). E-K. Representative bands (E) and quantification (F-K) of pyroptosis pathway-associated proteins in each group (n = 10/group, one-way ANOVA with post-hoc Tukey HSD test). L-M. Representative HE staining (L) and quantification of inflammation infiltration (M) in each group (n = 10/group, one-way ANOVA with post-hoc Tukey HSD test). The left ventricle myocardium and right ventricle epicardium in each group are shown respectively. The immune cells in each group are marked with black arrows, and the disordered cardiomyocytes are marked with yellow arrows. N-O. Representative immunofluorescence images (O) and quantification (N) in each group, with staining NLRP3 in red, cTnT in green and nuclei in blue (n = 5/group, one-way ANOVA with post-hoc Tukey HSD test). Scale bar, 20μm. Data were presented as mean ± SD. AGPAT2, 1-acylglycerol-3-phosphate O-acyltransferase 2; AGPAT2^+/-^, heterozygous AGPAT2 knockout; αPD1, anti-PD1 (PD1 inhibitor); Caspase-1, Cysteine-aspartic proteases 1; CK-MB, creatine kinase-myocardial band; cleaved Gasdermin D, N-terminal gasdermin domain; cTnT, cardiac troponin T; GSDMD, Gasdermin D; HE staining, Hematoxylin and Eosin staining; ICIAM, immune checkpoint inhibitor-associated myocarditis; IgG, immunoglobulin G; IL-1β, Interleukin-1β; IL-18, Interleukin-18; LVEF, left ventricular ejection fraction; LVFS, left ventricular fractional shortening; NLRP3, NOD-like receptor pyrin domain-containing protein 3; TCLPA5-4, inhibitor of LPAR5.

In the AGPAT2^+/-^ mice, compared to the αPD1 group, TCLPA5-4 treatment significantly increased LVEF and LVFS (Figure 7B-C), decreased release of cTnT and CK-MB (Figure 7D), limited levels of pyroptosis-related cell-released substances (IL-1β, IL-18, HMGB1, LDH), restored ATP content and downregulated expression of key proteins in pyroptosis pathways (Supplemental Figure 6A-E, Figure 7E-K), and reduced HE-stained myocardial immunocyte infiltration area (Figure 7L, M) and NLRP3, cleaved-Caspase 1, Granzyme B and GSDMD fluorescence intensity (Figure 7N, O, Supplemental Figure 6F-K), with most of these alterations restoring close to the levels in the Control group.

In WT mice, compared to the αPD1 group and the αPD1+LPA group, TCLPA5-4 treatment (αPD1+LPA+TCLPA5-4 group) significantly increased LVEF and LVFS (Supplemental Figure 7B, C), decreased release of cTnT and CK-MB (Supplemental Figure 7D-E), limited levels of IL-1β, IL-18, HMGB1, LDH, prevented ATP decline (Supplemental Figure 7F-J), downregulated expression of key proteins in pyroptosis pathways (Supplemental Figure 7K-Q), reduced HE-stained myocardial immunocyte infiltration area (Supplemental Figure 7R, S) and fluorescence intensity of NLRP3, cleaved Caspase-1, Granzyme B and GSDMD (Supplemental Figure 7T-X).

In melanoma-bearing mice, TCLPA5-4 treatment did not affect the antitumor effects of PD1 inhibitor as assessed by the tumor growth curve and the final tumor volume (Supplemental Figure 8B, C, F), could effectively restored LVEF and LVFS (Supplemental Figure 8D, E) and maintained notable aggregation of CD4^+^ and CD8^+^ T cells along the stromal interface, indicative of normal tumor-killing effects in anti-PD1 therapy (Supplemental Figure 8G).

Overall, these results indicate that blocking LPAR5 with TCLPA5-4 can inhibit activation of the LPA-LPAR5-apoptosis axis, relieve myocardial damage of ICIAM, without interfering the antitumor effects of PD1 inhibitors.

### Association Between LPA Level and Risk of ICIAM

Between August 2021 and August 2024, a cohort of 168 cancer patients with suspicious cardiac involvement, was enrolled based on the protocol outlined in Figure 8A, there were 115 ICI users and among them, 61 patients were diagnosed as ICIAM. The patients were categorized into three groups based on the presence of ICIAM (diagnostic criteria(**24**) and patients’ characteristics were described in supplementary material): Group A (No ICI+No myocarditis, n=53), Group B (ICI+No myocarditis, n=54), and Group C (ICI+Myocarditis, n=61) (Supplemental Table 3, Supplemental Table 4). There were no significant differences in baseline clinical features across the three groups (P>0.05) (Supplemental Table 3). The serum LPA levels were higher in ICI+No Myocarditis group, especially in ICI+Myocariditis group as compared to non-ICI users (Figure 8B).

**Figure 8.**
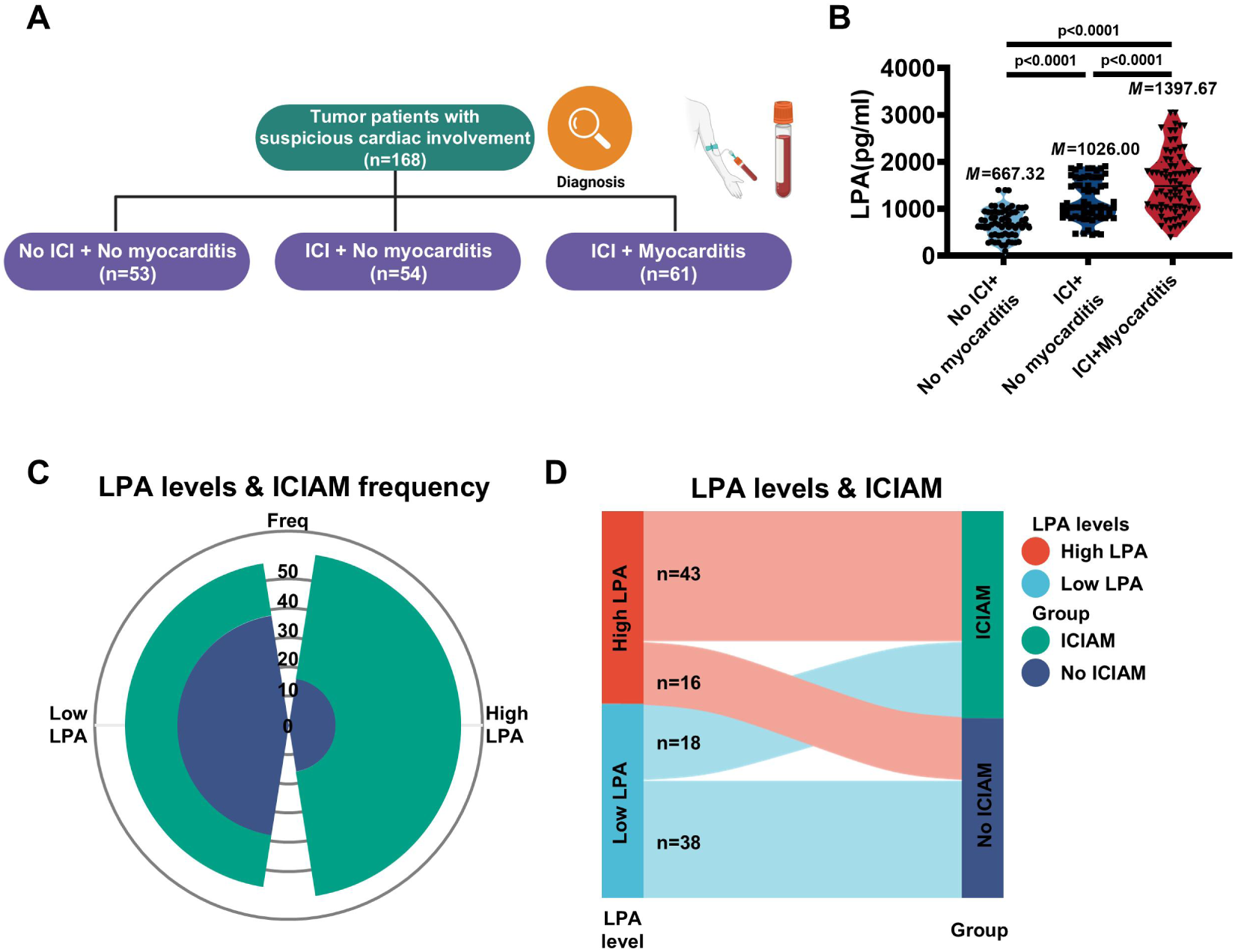
Association Between LPA Level and Risk of ICIAM. A. The study flowchart and participants. B. The serum LPA levels assessed by ELISA assay (n = 53 of No ICI+NO myocarditis group, n = 54 of ICI+NO myocarditis group, n = 61 of ICI+Myocarditis group, one-way ANOVA with post-hoc Sidak test). C-D. The rose diagram (C) and Sankey diagram (D) showing the relationship between serum LPA levels and ICIAM frequency. Continuous variables were presented as mean ± SD as distributed normally, as median (interquartile range, IQR) as skewed distribution. ICI, immune checkpoint inhibitor; ICI+Myocarditis, immune checkpoint inhibitor-associated myocarditis; ICI+NO myocarditis, patients with ICI but no myocarditis; LPA, lysophosphatidic acid; M, median; No ICI+NO myocarditis, patients without ICI and myocarditis.

To investigate the association between LPA levels and the risk of ICIAM, Groups B and C undergoing ICI therapy were analyzed. Using the median LPA value of 1153.98 pg/ml as the cut-point, high LPA (LPA>1153.98 pg/ml) and low LPA (≤1153.98 pg/ml) were grouped. The rose plot revealed a majority of patients in the high LPA group experienced ICIAM, whereas most in the low LPA group did not (Figure 8C). To be more specific, the Sankey diagram indicated that in the high LPA group, 43 cases (72.9%) had ICIAM, while 16 cases (27.1%) did not; in the low LPA group, 18 cases (32.1%) experienced ICIAM and 38 cases (67.9%) did not (Figure 8D) (Chi-square test, P<0.0001). Thus, elevated LPA levels may increase the likelihood of ICIAM.

## Discussion

Our study is the first to explore the roles of AGPAT2-IMRP in pathogenesis of ICIAM, and to assess novel diagnostic markers and potential therapeutic targets for ICIAM. The main findings are summarized as follows: 1) In αPD1-induced ICIAM mice, upstream immunity disorder initiated myocardial IMRP, leading to downstream AGPAT2 degradation via lysosome pathway accompanied by LPA accumulation. Accumulated LPA activated pyroptosis predominantly through LPAR5-NLRP3 pathways, causing pyroptotic myocardial damage and resultant ICIAM. LPAR5 knockdown by siRNA or inhibition with TCLPA5-4 effectively prevented myocardia from pyroptosis. 2) In ICI-treated patients, serum LPA elevation is linked with increased risk of ICIAM. These results supported that dysregulation of AGPAT2-LPA-LPAR5-pyroptosis axis involves in pathogenesis of ICIAM (Central Illustration) and several targets in this axis may be promising in diagnosis (LPA as a biomarker) and treatment option (lowering LPA or inhibiting LPAR5 as therapeutic targets) of ICIAM.

There are two basic theories in ICIAM pathogenesis: T cell cross-recognition and autoantigen generation(**11, 12, 14**). The activated T cell cross-recognition hypothesizes that molecular mimicry between tumor cells and heart tissue can cause T cell cross-reactivity, whereas the autoantigen generation assumes that autoantigen can induce autoimmune cardiotoxicity. The existing data support T cell-mediated immunity as a dominant mechanism, thereby attracting lots of further research enthusiasms, particularly in the following aspects: 1) to identify potential inciting antigens, 2) how to produce self-antigens and then to elicit an immune response, 3) additional roles of innate immunocytes and B cells. Undoubtedly, these key issues deserve to further elucidate so that precise treatment of ICIAM can be applied clinically. However, the ICIAM-associated immunity disorders, albeit critical, are only the upstream pathogenic component(**5, 12**). In fact, the downstream pathogenic effectors/regulators following initial immunity disorders are extremely complicated and remain little understood, especially in these aspects: 1) involvement of cardiac cell types, 2) cell death forms and pathways, 3) inflammation and its detriments to myocardium.

Recently, a very elegant translational study from Michel et al(**11**) sheds new light on the mechanisms of cardiotoxicity due to PD1 inhibition. The authors showed that LV dysfunction in tumor-bearing mice treated with anti-PD1 was accompanied by activation of CD8^+^ (dominants) and CD4^+^ T cells, together with dysregulation of proteome and lipidome metabolism, enhancing of oxidative stress and inflammation in myocardium, addressing the impacts of the downstream pathophysiology secondary to the upstream immunity disorders on cardiac dysfunction.

Likewise, concentrating on the downstream pathogenic components instead of the upstream immune dysfunction, our study is the first to demonstrate that dysregulation of AGPAT2-LPA-LPAR5-pyroptosis axis involves in the pathogenesis of ICIAM. As shown in our study, αPD1-induced myocardial infiltration of CD44^+^CD8^+^ and CD4^+^ T cells firstly initiated transcription and expression of proinflammatory cytokines and then led to AGPAT2 degradation along with LPA accumulation within lysosome via LAMP1 pathway. All of which are termed as “AGPAT2-IMRP”, similar to the “Warburg effect” proposed in oncology(**17, 26**) or the effect of inflammatory and/or metabolic reprogramming reported in the previous studies in immunology(**19, 27**). Accordingly, AGPAT2-IMRP becomes a core link of the upstream to the downstream components in this pathogenic axis.

The enzymatic function of AGPAT2 is vital in converting LPA into phosphatidic acid so that AGPAT2 downregulation/deficiency can cause LPA accumulation(**28, 29**). LPA, a multifunctional bioactive phospholipid intermediate in metabolism, acts as a signaling messenger by binding to LPARs(**30, 31**). Belonging to the G protein-coupled receptor (GPCR) family, LPARs consist of 6 subtypes, with major roles played by LPAR1, LPAR2, LPAR3 and LPAR5 in comprehensive pathophysiologic processes such as inflammation and pyroptotic cell death(**30, 32, 33**). As validated in this study, in cardiomyocyte membrane, LPAR5 was the dominant receptor of LPA and their combination, likely via intra- or extra-cellular domains of the receptor(**34**), could activate the G protein, exerting a critical effect on triggering subsequent cardiomyocyte injury. So, LPA-LPAR5 is another core link of the axis.

Despite there is little understanding of the terminal pathogenic components/ effectors of ICIAM as aforementioned especially in the involved cell types, cell death forms and pathways(**27, 35**). our study at least partially clarifies whether inflammation and cell death are critical or not and which of cell death programs is involved in ICIAM pathogenesis. Abundant data from our study illustrated that cardiomyocyte pyroptosis played a central role in myocardial injury, as evidenced by ruling out necroptosis or apoptosis with specific antagonists, by validating cardiomyocyte pyroptosis in cellular, animal and patient levels, and by stimulating LPAR5 with LPA only or combing with other stimulators or by blocking LPAR5 with TCLPA5-4 and their respective effects on the LPA-LPAR5-NLRP3-pyroptosis axis, all of which were also supported by theoretical effect of LPA-LPAR5-GPCR activation in which the G-protein acts as a crucial pyroptotic regulator influencing activation of NLRP3 inflammasome and caspases, thereby contributing to mediation of pyroptosis(**36, 37**).

Besides confirming the AGPAT2-LPA-LPAR5-NLRP3-pyroptosis axis involved in pathogenesis in cellular and animal levels, to further examine whether there are potential candidates in this axis can be translated into optimization of clinical practice, we finalized with a clinical study by including of a cohort of 116 cancer patients according to the prespecified study protocol and found that when the cancer patients were treated with anti-PD1, the elevation of serum LPA levels was associated with the increased risk of ICIAM.

Overall, this study confirms that the AGPAT2-LPA-LPAR5-NLRP3-pyroptosis axis involves in ICIAM pathogenesis, of which AGPAT2-IMRP is the most essential component linking the upstream of immunity disorders to the downstream of myocardial injury pathways. Moreover, our study identifies that at least two targets in this axis may become promising candidates in surveillance, diagnosis and treatment of ICIAM, thus being of clinical translational values detailed as follows:

Firstly, based on the findings that the myocardial LPA accumulation was noted in animal study and the serum LPA elevation observed in the clinical cohort of ICIAM patients, the serum LPA levels can be adopted as an indicative to surveil early stage of ICIAM since elevation of myocardial injury biomarkers of cTnT and CK-MB, reduction of echocardiographic left ventricular function of LVEF and LVFS are all not very sensitive to subclinical type of ICIAM patients(**24, 38, 39**). Additionally, controlling of the serum LPA levels may be expected to mitigate ICI-induced myocardial damage theoretically irrespective of no available data in this study.

Secondarily, the conventional medications for ICIAM such as glucocorticoids and neutralizing antibodies may interfere the effectiveness of cancer immunotherapy with ICIs, increase the risks of cardiovascular events, and reduce patient survival rates(**4, 7, 40–43**). Thus, avoiding direct intervention of immune cells or factors will become a desirable approach in ICIAM management. Several preclinical studies in oncology and cardiology suggested that inhibiting of the LPAR5 signal conferred beneficial effects on restoring the tumor-killing ability of CD8^+^ T cells, controlling tumor proliferation, invasion and immune escape, and reducing various cardiovascular events(**20, 36, 44**). Interestingly, this study also determined that LPAR5 acted as the primary receptor mediating myocardial pyroptotic damage and further confirmed that specific inhibition of LPA-LPAR5 function with TCLPA5-4 effectively alleviated ICIAM without promoting tumor progression in tumor-bearing mice, thus supporting that intervention with TCLPA5-4 may be a novel promising target.

### Limitations

This study has elucidated the involvement of the AGPAT2-IMRP-LPA axis in the pathophysiological processes of ICIAM across various levels, including in vitro cardiomyocytes, in vivo animal models, and clinical cases. However, there remain several limitations. First, the investigation of the ICIAM mechanism relies mostly on experiments in cellular and animal models, posing constraints on its extrapolation to clinical applications. Second, in the pilot study, female mice experienced high pre-modeling mortality rate (15% higher as compared to male mice), though no sex-based differences in ICIAM performance could be observed between male and female mice with successful modeling. In order to ensure higher successful modeling rate for subsequent TCLPA5-4 treatment experiments, only male mice were used for ICIAM model in this study. Future research is needed to include modeling mice from both sexes to enhance the study’s relevance. Third, due to the ethical issues, blocking of LPA-LPAR5 with TCLPA5-4 is performed only on animal model rather than clinical cases, rendering the extrapolation of the efficacy of TCLPA5-4 uncertain clinically. Finally, because of low incidence of ICIAM, we do not include a large number of patients with long-term follow-up in our cohort, thereby preventing us from drawing definite conclusion about roles of LPA in surveillance and intervention of ICIAM patients.

## Conclusions

Our study demonstrates that a novel mechanism of the AGPAT2-LPA-LPAR5 -NLRP3-pyroptosis axis is involved in ICIAM pathogenesis, of which anti-PD1-induced AGPAT2-IMRP is the most essential component linking the upstream of immunity disorders to the downstream of myocardial injury pathways (Central Illustration). Additionally, our study also identifies that at least two targets in this axis may serve as promising candidates in surveillance, diagnosis and treatment of ICIAM, thus being of clinical translational values However, large-scale clinical studies are still warranted for further validation of our findings since the current data are mainly from cellular or animal experiments.

## Declarations

### Author Contributions

**YC** designed animal experiments, conducted animal modeling and evaluating, collected data and write the first draft, revised the article and took public responsibility for the content. **YS, HZ, XW, YX, JZ, XH and MH** assisted in conducting animal experiments, animal parameter evaluation and statistics, reviewing the drafts for major intellectual content. **XW, JZ, XH and MH** helped in data processing, statistics, secondary inspection of intellectual content and article polishing. **JG and LC** designed the technological roadmap of the article together, guided the experiment and participated in the revision of the manuscript. All the authors above have given final approval of the version to be published and agreed to be fully accountable for all aspects of the work, ensuring thorough investigation and resolution of any inquiries regarding its accuracy or integrity.

## Data Availability Statement

All data points generated or analyzed during this study are included in this article or supplementary materials and there are no further underlying data necessary to reproduce the results.

## Ethical approval

The ARRIVE guidelines (Animal Research: Reporting of In Vivo Experiments guidelines, version 2.0) for the care and use of laboratory animals were followed and approved by Animal Ethics Committee of Zhongshan Hospital, Fudan University (Ethics approval No. 2019-09). For clinical studies, the protocol was approved by the Ethics Committee of Zhongshan Hospital, Fudan University (Ethics approval No. B2021-275).

## Funding

This study was supported by the National Natural Science Foundation of China (No.81201095), the Young Scientists Fund of the National Natural Science Foundation of China (No.82500442), Shanghai Shenkang Hospital Development Center municipal hospital diagnosis and treatment technology project (SHDC22023207) and Non-profit Central Research Institute Fund of Chinese Academy of Medical Sciences (2024-JKCS-28).

## Conflict- of- Interest

All authors declare no disclosure of interest and no relationships with industry.

## Acknowledgments

All authors would like to thank Aijun Sun of Shanghai Institute of Cardiovascular Diseases for helpful discussions related to this work, and Jianguo Jia of National Clinical Research Center for Interventional Medicine for pathological section processing.

## List of abbreviations

αPD1: anti-PD1
AGPAT2: 1-acylglycerol-3-phosphate O-acyltransferases
irAE: immune-related adverse event
ICI: Immune checkpoint inhibitor
ICIAM: ICI-associated myocarditis
IMRP: inflammatory metabolic reprogramming
LPA: lysophosphatidic acid
LPAR: LPA receptor
LPAR5: LPA receptor subtype 5
PD1: programmed cell death protein-1

## Central Illustration

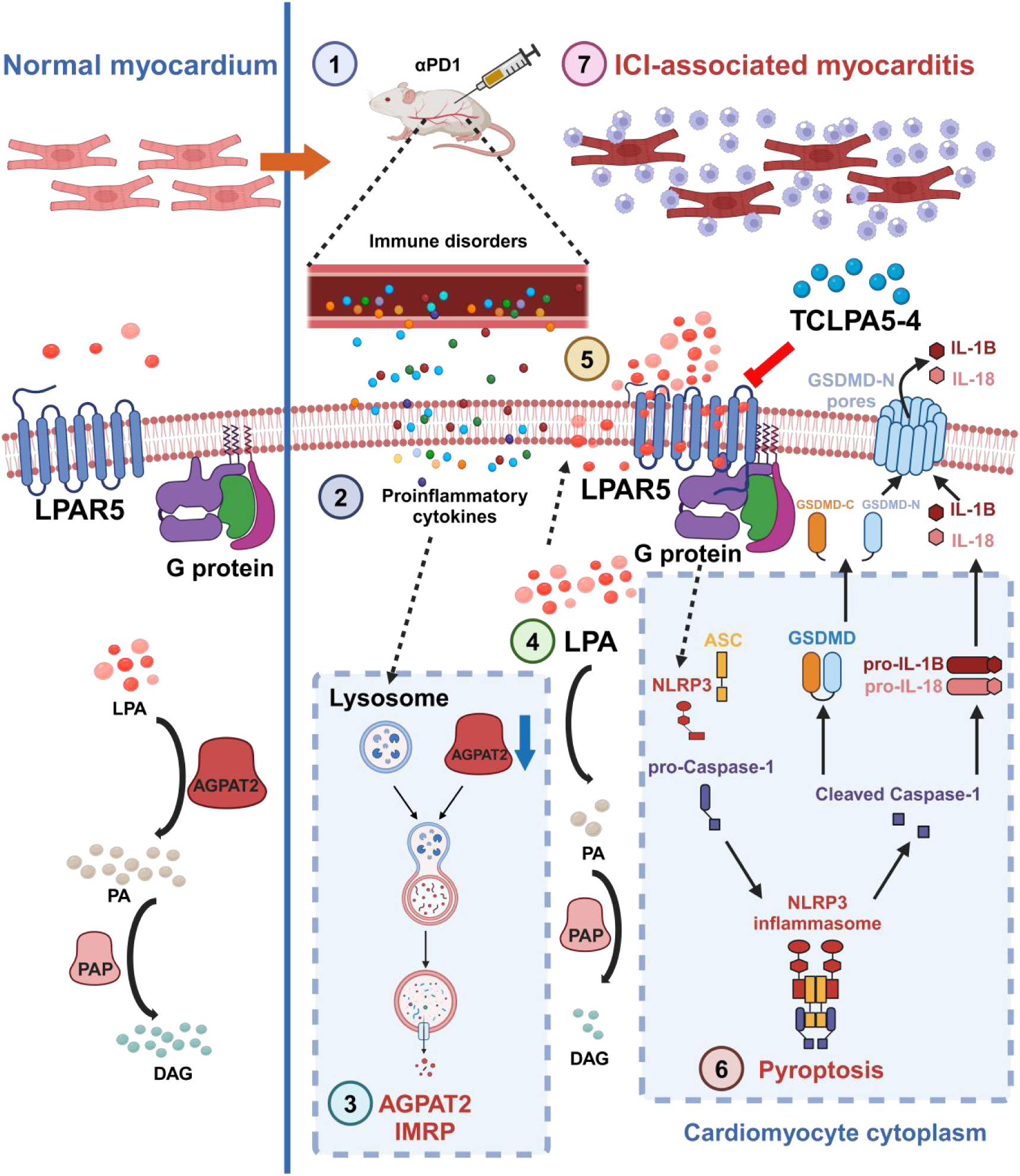

### AGPAT2-LPA-LPAR5-Pyroptosis Axis Plays a Key Role in ICIAM Pathogenesis

PD1 inhibitor-induced immunity disorder can cause myocardial AGPAT2-IMRP via lysosome, resulting in AGPAT2 degradation and LPA accumulation. The accumulated LPA binds to LPAR5 in cardiomyocyte and then activates LPAR5-NLRP3-pyroptosis pathway, thus leading to myocardial injury and resultant ICIAM. αPD1, PD1 inhibitor; IMRP, inflammatory metabolic reprogramming; AGPAT2, 1-acylglycerol-3-phosphate O-acyltransferase 2; LPA, lysophosphatidic acid; PA, phosphatidic acid; PAP, phosphatidic acid; DAG, diacylglycerol; LPAR5, lysophosphatidic acid receptor 5; GSDMD-C, C-terminal gasdermin domain; GSDMD-N, N-terminal gasdermin domain; ICIAM, immune checkpoint inhibitor-associated myocarditis.

## Supplemental Tables

**Supplemental Table 1.** Antibodies.

| Target antigen | Vendor Source | Catalog # | Working concentration | Persistent ID /URL |
|---|---|---|---|---|
| NLRP3 | Abcam | ab283819 | WB 1:1000<br>IF 1:50 | https://www.abcam.cn/products/primary-antibodies/nlrp3-antibody-rm1021-ab283819.html#lb |
| pro Caspase-1+p10+p12 | Abcam | ab179515 | WB 1:1000 | https://www.abcam.cn/products/primary-antibodies/pro-caspase-1-p10-p12-antibody-epr16883-ab179515.html |
| Gasdermin D | Cell Signaling Technology | 39754 | WB 1:1000 | https://www.cellsignal.cn/products/primary-antibodies/gasdermin-d-e9s1x-rabbit-mab/39754 |
| cleaved Gasdermin D | Cell Signaling Technology | 10137 | WB 1:1000 | https://www.cellsignal.cn/products/primary-antibodies/cleaved-gasdermin-d-asp276-e3e3p-rabbit-mab/10137 |
| IL-1β | Cell Signaling Technology | 31202 | WB 1:1000 | https://www.cellsignal.cn/products/primary-antibodies/il-1b-d6d6t-rabbit-mab/31202 |
| IL-18 | Cell Signaling Technology | 57058 | WB 1:1000 | https://www.cellsignal.cn/products/primary-antibodies/il-18-e8p5o-rabbit-mab/57058 |
| AGPAT2 | Novus | NBP2-92662 | WB 1:1000<br>IF 1:500<br>IP 1:200 | https://www.novusbio.com/products/agpat2-antibody_nbp2-92662 |
| LAMP1 | DSHB | 1D4B | WB 1:1000<br>IF 1:500<br>IP 1:200 | https://dshb.biology.uiowa.edu/1D4B |
| β-tubulin | Cell Signaling Technology | 86298 | WB 1:5000 | https://www.cellsignal.cn/products/primary-antibodies/b-tubulin-d3u1w-mouse-mab/86298 |
| isotype control immunoglobulin G | Cell Signaling Technology | 3900 | IP 1:200 | https://www.cellsignal.cn/products/primary-antibodies/rabbit-da1e-mab-igg-xp-isotype-control/3900 |
| cardiac Troponin T | Abcam | ab8295 | IF 1:500 | https://www.abcam.cn/products/primary-antibodies/cardiac-troponin-t-antibody-1c11-ab8295.html |
| CD4 | Cell Signaling Technology | 82116 | IF 1:500 | https://www.cellsignal.cn/products/antibody-conjugates/cd4-rm4-5-rat-mab-apc-conjugate/82116 |
| CD8 | Cell Signaling Technology | 98941 | IF 1:500 | https://www.cellsignal.cn/products/primary-antibodies/cd8a-d4w2z-xp-rabbit-mab/98941 |
| CD44 | Cell Signaling Technology | 95235 | IF 1:500 | https://www.cellsignal.cn/products/antibody-conjugates/cd44-im7-rat-mab-alexa-fluor-555-conjugate/95235 |
| Granzyme B | Proteintech | 13588-1-AP | IF 1:100 | https://www.ptgcn.com/products/GZMB-Antibody-13588-1-AP.htm#product-information |
| GSDMD | Proteintech | 66387-1-Ig | IF 1:50 | https://www.ptgcn.com/products/GSDMD-Antibody-66387-1-Ig.htm#product-information |
| Caspase-1 | Abclonal | A0964 | IF 1:100 | https://abclonal.com.cn/catalog/A0964 |
| goat anti-rabbit IgG (H+L) | Cell Signaling Technology | 14708 | WB 1:10,000 | https://www.cellsignal.cn/products/secondary-antibodies/anti-rabbit-igg-h-l-biotinylated-antibody/14708 |
| goat anti-mouse IgG (H+L) | Cell Signaling Technology | 14709 | WB 1:10,000 | https://www.cellsignal.cn/products/secondary-antibodies/anti-mouse-igg-h-l-biotinylated-antibody/14709 |
| goat anti-rat IgG (H+L) | Cell Signaling Technology | 7077 | WB 1:10,000 | https://www.cellsignal.cn/products/secondary-antibodies/anti-rat-igg-hrp-linked-antibody/7077 |
| goat anti-rabbit IgG (H+L) (Alexa Fluor® 488 Conjugate) | Cell Signaling Technology | 4412 | WB 1:1000 | https://www.cellsignal.cn/products/secondary-antibodies/anti-rabbit-igg-h-l-f-ab-2-fragment-alexa-fluor-488-conjugate/4412 |
| goat anti-mouse IgG (H+L) (Alexa Fluor® 594 Conjugate) | Cell Signaling Technology | 8890 | WB 1:1000 | https://www.cellsignal.cn/products/secondary-antibodies/anti-mouse-igg-h-l-f-ab-2-fragment-alexa-fluor-594-conjugate/8890 |
| goat anti-rat IgG (H+L) (Alexa Fluor® 647 Conjugate) | Cell Signaling Technology | 4418 | WB 1:1000 | https://www.cellsignal.cn/products/secondary-antibodies/anti-rat-igg-h-l-alexa-fluor-647-conjugate/4418 |

IF: Immunofluorescence; IP: Immunoprecipitation; WB: Western immunoblot.

**Supplemental Table 2.**
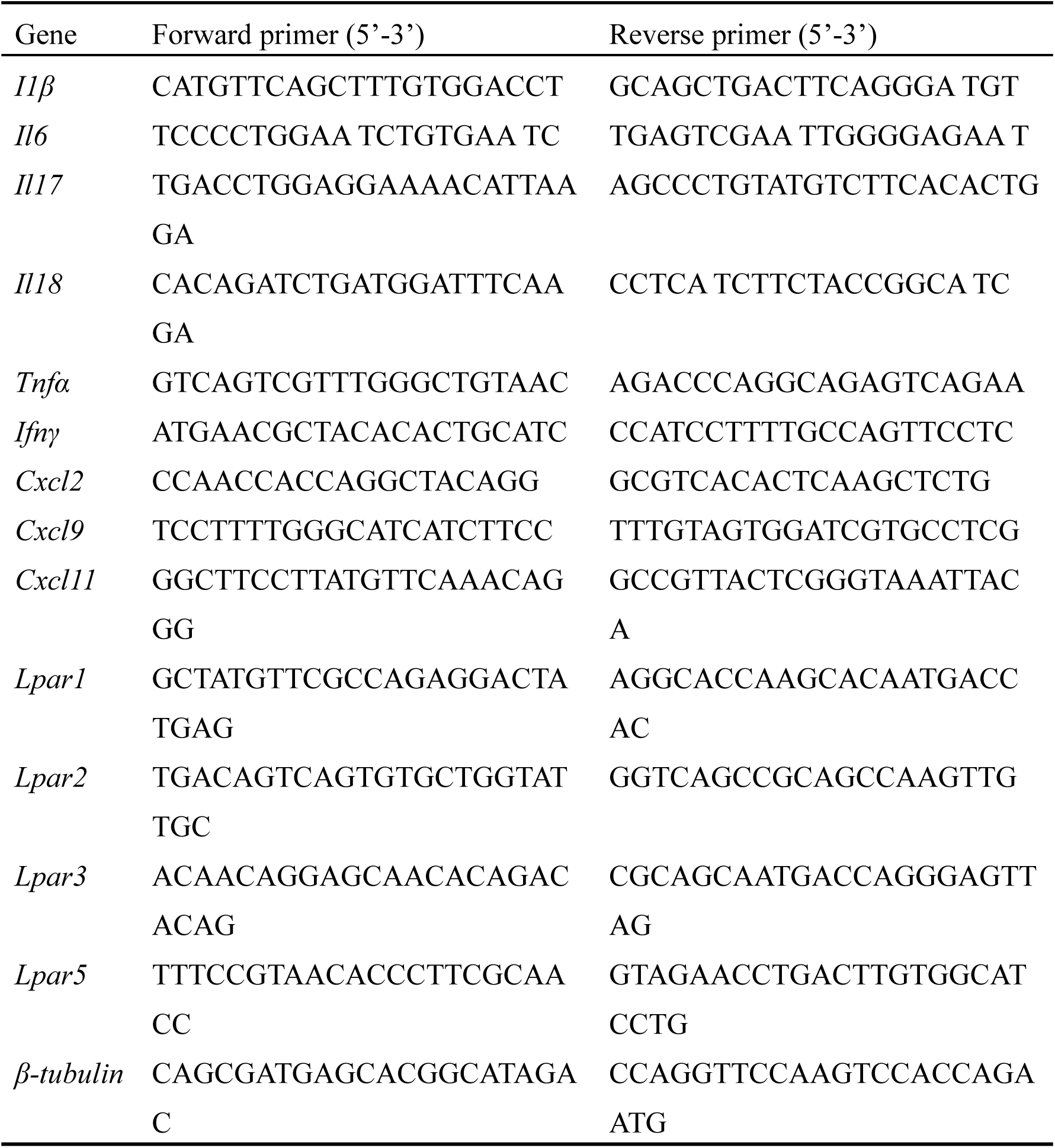
RT-qPCR Primers.

**Supplemental Table 3.** Baseline characteristics of tumor patients.

|  | Gruop A<br>(n=53) | Gruop B<br>(n=54) | Gruop C<br>(n=61) | Across groups<br>p value |
|---|---|---|---|---|
| Age at presentation (years) | 66 (57,70) | 61 (54,68) | 66 (60,72) | 0.20 |
| Females | 16 (30) | 16 (30) | 16 (26) | 0.87 |
| Past smoker | 3 (6) | 3 (6) | 2 (3) | 0.81 |
| Active smoker | 3 (6) | 2 (4) | 5 (8) | 0.61 |
| Hypertension | 32 (60) | 30 (55) | 36 (59) | 0.87 |
| Diabetes mellitus | 16 (30) | 13 (24) | 17 (28) | 0.77 |
| Dyslipidemia | 8 (15) | 10 (19) | 12 (20) | 0.80 |
| CVA/TIA | 1 (2) | 0 (0) | 2 (3) | 0.42 |
| CAD | 8 (15) | 7 (13) | 11 (18) | 0.75 |
| Prior history of LV systolic dysfunction | 2 (4) | 2 (4) | 6 (10) | 0.27 |
| Hypothyroidism | 0 (0) | 0 (0) | 2 (3) | 0.17 |

Data are presented as median (lower quartile, upper quartile) or quantity (percentage), as appropriate. Gruop A: No ICI+No myocarditis group; Gruop B: ICI+No myocarditis group; Gruop C, ICI+Myocarditis (ICIAM) group; CAD, coronary artery disease; CVA, cerebrovascular accident; ICI, immune-checkpoint inhibitors; TIA, transient ischemic attack. P values are calculated using one-way ANOVA with post-hoc Tukey HSD test for measurement data and Fisher's exact test for count data.

**Supplemental Table 4.** Clinical synopses, diagnostic criteria, therapies and outcomes of enrolled ICIAM patients.

| Patient ID | Gender | Tumor Type | Major characters | cTnT (ng/ml) | LVEF(%) | Diagnostic criteria | Steroid Therapy | Outcome<sup>a</sup> |
|---|---|---|---|---|---|---|---|---|
| 1 | M | lung squamous cell carcinoma | chest tightness, new decreased LVEF | 0.749 | 48 | cTnT elevation+2 minor criteria | √ | survival |
| 2 | M | hepatocellular carcinoma | fatigue, new prolonged QRS duration | 0.058 | 66 | cTnT elevation+2 minor criteria | √ | survival |
| 3 | M | lung small cell lung carcinoma | chest pain, new IVB | 0.047 | 62 | cTnT elevation+2 minor criteria | √ | survival |
| 4 | M | lung squamous cell carcinoma | fatigue, new ventricular arrhythmia | 0.257 | 66 | cTnT elevation+2 minor criteria | √ | survival |
| 5 | M | lung adenocarcinoma | shortness of breath, new decreased LVEF | 0.471 | 46 | cTnT elevation+2 minor criteria | √ | survival |
| 6 | M | gastric adenocarcinoma | fatigue, new IVB | 0.141 | 65 | cTnT elevation+2 minor criteria | √ | survival |
| 7 | M | lung adenocarcinoma | myositis, CMR：acute myocarditis | 0.975 | 53 | cTnT elevation+1 major criteria | √ | survival |
| 8 | M | lung adenocarcinoma | chest pain, new IVB | 0.056 | 66 | cTnT elevation+2 minor criteria | √ | survival |
| 9 | F | gastric adenocarcinoma | chest pain, new ventricular arrhythmia | 0.089 | 65 | cTnT elevation+2 minor criteria | √ | survival |
| 10 | M | liver adenocarcinoma | chest pain, new decreased LVEF | 0.143 | 49 | cTnT elevation+2 minor criteria | √ | survival |
| 11 | M | gastric adenocarcinoma | shortness of breath, new decreased LVEF | 0.290 | 48 | cTnT elevation+2 minor criteria | √ | survival |
| 12 | M | gastric adenocarcinoma | dizziness，ventricular arrhythmia | 0.316 | 67 | cTnT elevation+2 minor criteria | √ | survival |
| 13 | M | intrahepatic cholangiocarcin-oma | diplopia, new prolonged QRS duration | 0.100 | 64 | cTnT elevation+2 minor criteria | √ | survival |
| 14 | M | liver adenocarcinoma | palpitations, ventricular arrhythmia | 0.075 | 67 | cTnT elevation+2 minor criteria | √ | survival |
| 15 | F | intrahepatic cholangiocarcinoma | shortness of breath, new AVB | 0.187 | 65 | cTnT elevation+2 minor criteria | √ | non-survival: died of tumor after 10 months |
| 16 | F | esophagus adenocarcinoma | myalgias, prolonged QRS duration | 0.307 | 57 | cTnT elevation+2 minor criteria | √ | survival |
| 17 | M | gastric adenocarcinoma | lower-extremity oedema, new decreased LVEF | 0.300 | 45 | cTnT elevation+2 minor criteria | √ | survival |
| 18 | M | lung squamous cell carcinoma | fatigue, new AVB | 0.108 | 62 | cTnT elevation+2 minor criteria | √ | survival |
| 19 | F | intrahepatic cholangiocarcinoma | palpitations, suggestive CMR: myocarditis | 0.342 | 55 | cTnT elevation+2 minor criteria | √ | survival |
| 20 | M | colon adenocarcinoma | palpitations, new decreased LVEF, myasthenia gravis | 0.398 | 48 | cTnT elevation+3 minor criteria | √ | survival |
| 21 | M | liver adenocarcinoma | dizziness, new prolonged QRS duration | 0.101 | 58 | cTnT elevation+2 minor criteria | √ | survival |
| 22 | M | hepatocellular carcinoma | chest pain, ventricular arrhythmia | 0.230 | 68 | cTnT elevation+2 minor criteria | √ | non-survival: died of tumor after 18 months |
| 23 | M | hepatocellular carcinoma | shortness of breath, new prolonged QRS duration | 0.047 | 66 | cTnT elevation+2 minor criteria | √ | survival |
| 24 | F | intrahepatic cholangiocarcinoma | fatigue, new IVB | 0.281 | 65 | cTnT elevation+2 minor criteria | √ | survival |
| 25 | M | lung adenocarcinoma | shortness of breath, ventricular arrhythmia | 0.213 | 65 | cTnT elevation+2 minor criteria | √ | survival |
| 26 | F | thymic squamous cell carcinoma | diplopia, multiple myositis | 0.922 | 63 | cTnT elevation+2 minor criteria | √ | non-survival: died of irAE after 3 months |
| 27 | F | intrahepatic cholangiocarcinoma | shortness of breath, new decreased LVEF, CMR: acute myocarditis | 0.883 | 45 | cTnT elevation+1 major+2 minor criteria | √ | survival |
| 28 | F | rectal adenocarcinoma | shortness of breath, CMR： acute myocarditis | 1.190 | 69 | cTnT elevation+1 major criteria | √ | non-survival: died of irAE after 1 month |
| 29 | F | cervical squamous cell carcinoma | dizziness, new AVB | 0.110 | 58 | cTnT elevation+2 minor criteria | √ | survival |
| 30 | F | liver cancer (undifferentiated) | fatigue, new IVB | 0.124 | 69 | cTnT elevation+2 minor criteria | √ | survival |
| 31 | M | intrahepatic cholangiocarcinoma | palpitations, new prolonged QRS duration | 0.389 | 67 | cTnT elevation+2 minor criteria | √ | non-survival: died of tumor after 15 months |
| 32 | M | intrahepatic cholangiocarcinoma | chest pain, new ventricular arrhythmia | 0.058 | 63 | cTnT elevation+2 minor criteria | √ | non-survival: died of tumor after 12 months |
| 33 | M | nasopharyngeal squamous cell carcinoma | shortness of breath, new prolonged QRS duration | 0.121 | 65 | cTnT elevation+2 minor criteria | √ | survival |
| 34 | M | lung squamous cell carcinoma | fatigue, new AVB | 0.062 | 66 | cTnT elevation+2 minor criteria | √ | survival |
| 35 | F | lung squamous cell carcinoma | lower-extremity oedema, new decreased LVEF, | 0.548 | 44 | cTnT elevation+2 minor criteria | √ | survival |
| 36 | M | gastric adenocarcinoma | shortness of breath, new AVB | 0.075 | 64 | cTnT elevation+2 minor criteria | √ | survival |
| 37 | M | lung adenocarcinoma | muscle weakness, new IVB | 0.266 | 58 | cTnT elevation+2 minor criteria | √ | survival |
| 38 | M | gastric adenocarcinoma | fatigue, new prolonged QRS duration | 1.930 | 62 | cTnT elevation+2 minor criteria | √ | survival |
| 39 | M | lung adenocarcinoma | dizziness, new AVB | 0.086 | 66 | cTnT elevation+2 minor criteria | √ | survival |
| 40 | M | esophagus adenocarcinoma | shortness of breath, myalgias | 0.398 | 67 | cTnT elevation+2 minor criteria | √ | survival |
| 41 | M | gastric adenocarcinoma | fatigue, new AVB | 0.024 | 65 | cTnT elevation+2 minor criteria | √ | survival |
| 42 | M | hepatocellular carcinoma | fatigue, new decreased LVEF, CMR： acute myocarditis | 0.039 | 39 | cTnT elevation+1 major criteria+2 minor criteria | √ | survival |
| 43 | M | intrahepatic cholangiocarcinoma | fatigue, new AVB | 1.410 | 65 | cTnT elevation+2 minor criteria | √ | non-survival: died of tumor after 20 months |
| 44 | M | intrahepatic cholangiocarcinoma | chest pain, new AVB | 0.033 | 67 | cTnT elevation+2 minor criteria | √ | survival |
| 45 | M | hepatocellular carcinoma | shortness of breath, new ventricular arrhythmia | 0.105 | 65 | cTnT elevation+2 minor criteria | √ | non-survival: died of tumor after 18 months |
| 46 | M | hepatocellular carcinoma | muscle weakness, suggestive CMR：myocarditis | 2.870 | 60 | cTnT elevation+2 minor criteria | √ | survival |
| 47 | F | gastric signet ring cell carcinoma | shortness of breath, new IVB | 0.770 | 72 | cTnT elevation+2 minor criteria | √ | non-survival: died of tumor after 8 months |
| 48 | M | breast cancer | shortness of breath, suggestive CMR：myocarditis | 0.030 | 61 | cTnT elevation+2 minor criteria | √ | survival |
| 49 | M | rectal adenocarcinoma | shortness of breath, new IVB | 0.164 | 67 | cTnT elevation+2 minor criteria | √ | survival |
| 50 | F | nasopharyngeal squamous cell carcinoma | fatigue, new prolonged QRS duration | 0.726 | 69 | cTnT elevation+2 minor criteria | √ | survival |
| 51 | M | lung small cell lung carcinoma | fatigue, new ventricular arrhythmia | 0.079 | 65 | cTnT elevation+2 minor criteria | √ | survival |
| 52 | M | lung adenocarcinoma | dizziness, new prolonged QRS duration | 0.082 | 66 | cTnT elevation+2 minor criteria | √ | survival |
| 53 | M | bladder urothelial carcinoma | shortness of breath, new decreased LVEF | 0.157 | 47 | cTnT elevation+2 minor criteria | √ | survival |
| 54 | M | lung adenocarcinoma | palpitations, new decreased LVEF | 0.619 | 49 | cTnT elevation+2 minor criteria | √ | survival |
| 55 | M | esophagus adenocarcinoma | fatigue, new IVB | 0.139 | 64 | cTnT elevation+2 minor criteria | √ | survival |
| 56 | F | esophagus adenocarcinoma | palpitations, new ventricular arrhythmia | 0.422 | 62 | cTnT elevation+2 minor criteria | √ | survival |
| 57 | M | bladder urothelial carcinoma | shortness of breath, new IVB | 0.699 | 62 | cTnT elevation+2 minor criteria | √ | survival |
| 58 | M | lung adenocarcinoma | dizziness, new AVB | 0.404 | 65 | cTnT elevation+2 minor criteria | √ | survival |
| 59 | M | lung adenocarcinoma | fatigue, new ventricular arrhythmia | 0.125 | 66 | cTnT elevation+2 minor criteria | √ | survival |
| 60 | F | breast cancer | palpitations, new ventricular arrhythmia | 0.147 | 69 | cTnT elevation+2 minor criteria | √ | survival |
| 61 | F | breast cancer | lower-extremity oedema, new decreased LVEF | 0.344 | 48 | cTnT elevation+2 minor criteria | √ | survival |

AVB, atrioventricular block; CMR, cardiac magnetic resonance imaging; cTnT, cardiac troponin T; F, female; ICIAM, ICI-associated myocarditis; irAE, immune-related adverse event; IVB, intraventricular conduction delay; LVEF, left ventricular ejection fraction; M, male. <sup>a</sup> Outcome was defined as survival or death during the observation period (between August 2021 and August 2024).

## Supplemental Figures

**Supplemental Figure 1.**
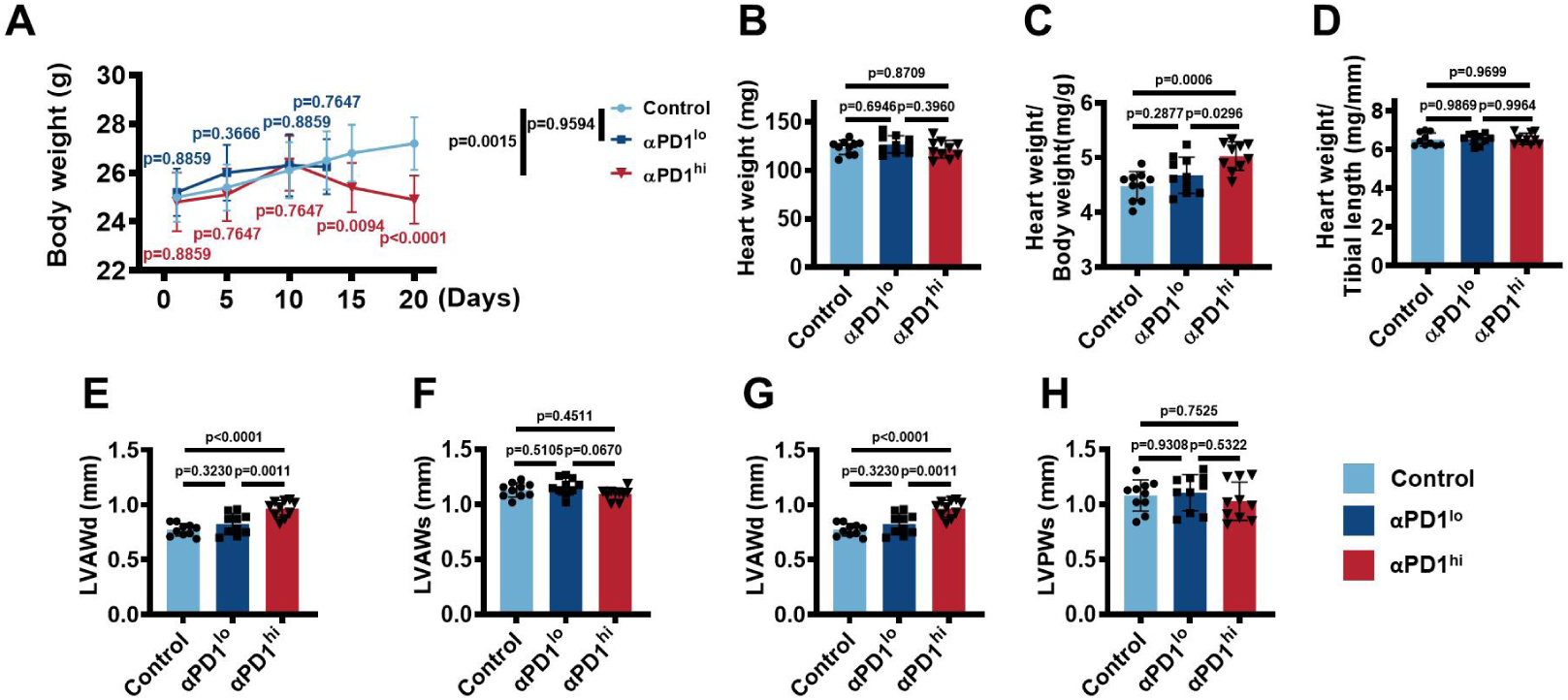
A Novel ICIAM Mouse Model Recapitulating ICIAM Patients’ Characteristics. A. Changes of mice body weight throughout the experimental period in control, αPD1^lo^ and αPD1^hi^ group (n = 10/group, two-way ANOVA with post-hoc Tukey HSD test). B-D. Heart weight (B), heart weight to body weight ratio (C) and heart weight to tibial length ratio (D) in each group (n = 10/group, one-way ANOVA with post-hoc Tukey HSD test). E-H. Quantification of LVAWd (E), LVAWs (F), LVPWd (G) and LVPWs (H) in each group (n = 10/group, one-way ANOVA with post-hoc Tukey HSD test). Data were presented as mean ± standard deviation (SD) when there were at least 3 independent experimental data available (The same requirement for data expression unless specifically indicated in the following figures). αPD1^hi^, 8 times of αPD1; αPD1^lo^, 5 times of αPD1; LVAWd, diastolic left ventricular anterior wall thickness; LVAWs, systolic left ventricular anterior wall thickness; LVPWd, diastolic left ventricular posterior wall thickness; LVPWs, systolic left ventricular posterior wall thickness.

**Supplemental Figure 2.**
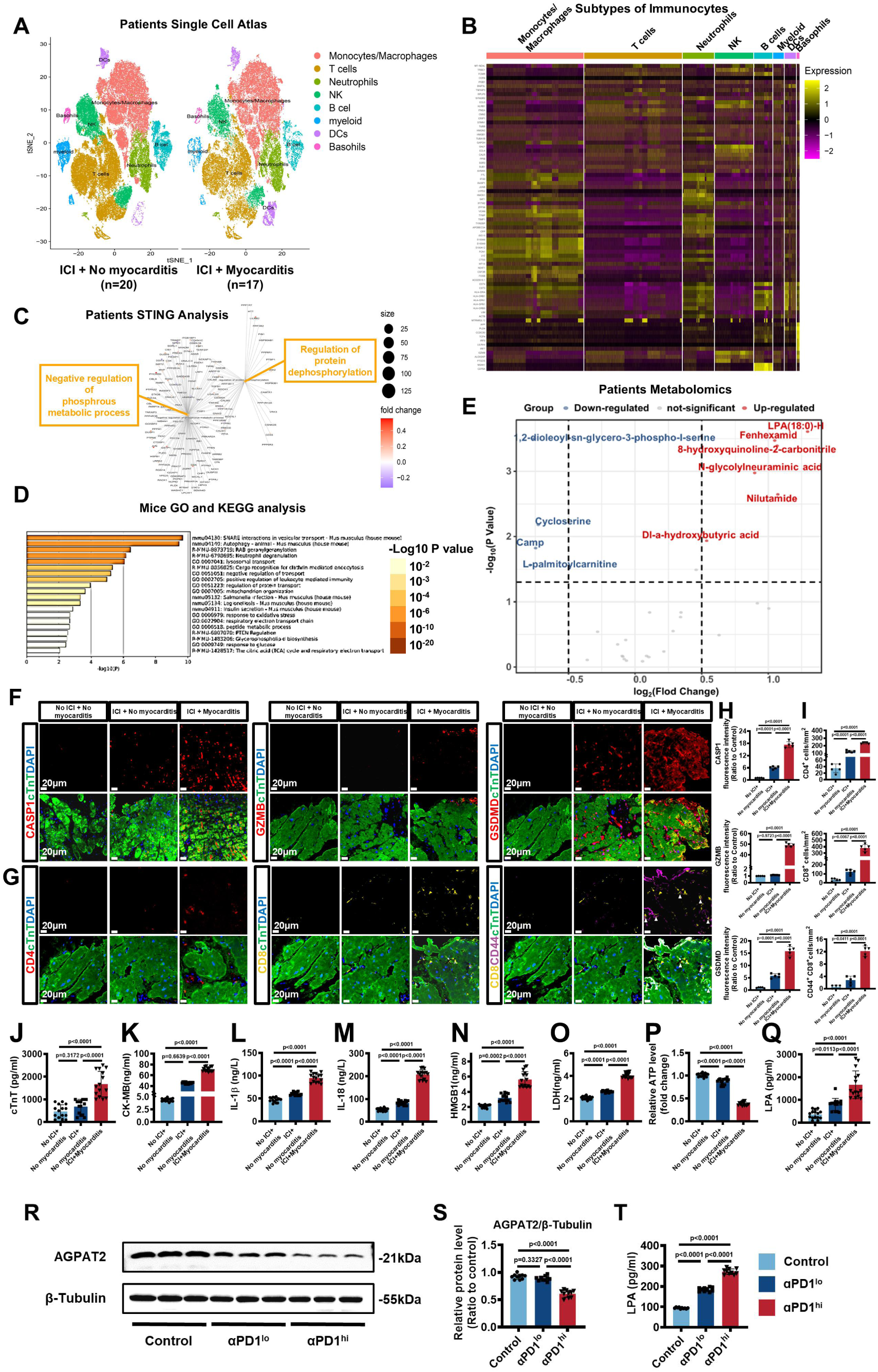
Omics and Confirmation of AGPAT2-IMRP in ICIAM Patients and Mice. A. The single-cell atlas of myocardial and peripheral blood mononuclear cells (PBMCs) composed of 8 different subtypes in ICI+No myocarditis (n=20) and ICI+Myocarditis (n=17) groups was constructed by cluster analysis. B. Machine annotation combined with cell molecular markers for annotating of different immune cell subgroups. C. The gene interaction network suggested that the differential genes were mainly related to phosphorus metabolic process. D. GO and KEGG analysis showed the most significant differential signaling pathways. E. Volcano plot of differential lipid metabolites in tumor patients with ICI+No myocarditis or ICI+Myocarditis (n = 5/group). F, H. Representative immunofluorescence staining (F) and quantification (H) of Caspase 1, Granzyme B and GSDMD in myocardial biopsy sections from patients, with staining Caspase 1, Granzyme B and GSDMD in red, respectively; cTnT in green and nuclei in blue in each group (n = 5/group, one-way ANOVA with post-hoc Tukey HSD test). Scale bar, 20μm. G, I. Representative immunofluorescence staining (G) and quantification (I) of CD4^+^ cells, CD8^+^ cells and CD44^+^CD8^+^ double-positive cells (marked with white arrows) in myocardial biopsy sections from patients, with staining CD4 in red, CD8 in yellow, CD44 in magenta, cTnT in green and nuclei in blue in each group (n = 5/group, one-way ANOVA with post-hoc Tukey HSD test). Scale bar, 20μm. J-Q. The serum cTnT, CK-MB, IL-1β, IL-18, HMGB1, LDH and LPA levels of patients measured by ELISA assays, and the ATP level measured by ATP assay (n = 15/group, one-way ANOVA with post-hoc Tukey HSD test). R-S. Representative bands (R) and quantification (S) of AGPAT2 protein in mice hearts from Control, αPD1^lo^ and αPD1^hi^ groups (n = 10/group, one-way ANOVA with post-hoc Tukey HSD test). T. The serum LPA level of mice measured by ELISA assay (n = 10/group, one-way ANOVA with post-hoc Tukey HSD test). Data were presented as mean ± standard deviation (SD) when there were at least 3 independent experimental data available. αPD1, anti-PD1 (PD1 inhibitor); αPD1^hi^, 8 times of αPD1; αPD1^lo^, 5 times of αPD1; AGPAT2, 1-acylglycerol-3-phosphate O-acyltransferase 2; ATP, adenosine triphosphate; CASP1, Caspase 1, Cysteine-aspartic proteases 1cysteine-aspartic proteases 1; CK-MB, creatine kinase-myocardial band; cTnT, cardiac troponin T; DC, dendritic cell; GO, Gene Ontology; GSDMD, GasderminD; GZMB, Granzyme B; HMGB1, high-mobility group box 1; IL-1β, interleukin-1 beta; IL-18, Interleukin-18; KEGG, Kyoto Encyclopedia of Genes and Genomes; LDH, lactate dehydrogenase; LPA, lysophosphatidic acid; NK, natural killer cell; NLRP3, NOD-like receptor pyrin domain-containing protein 3.

**Supplemental Figure 3.**
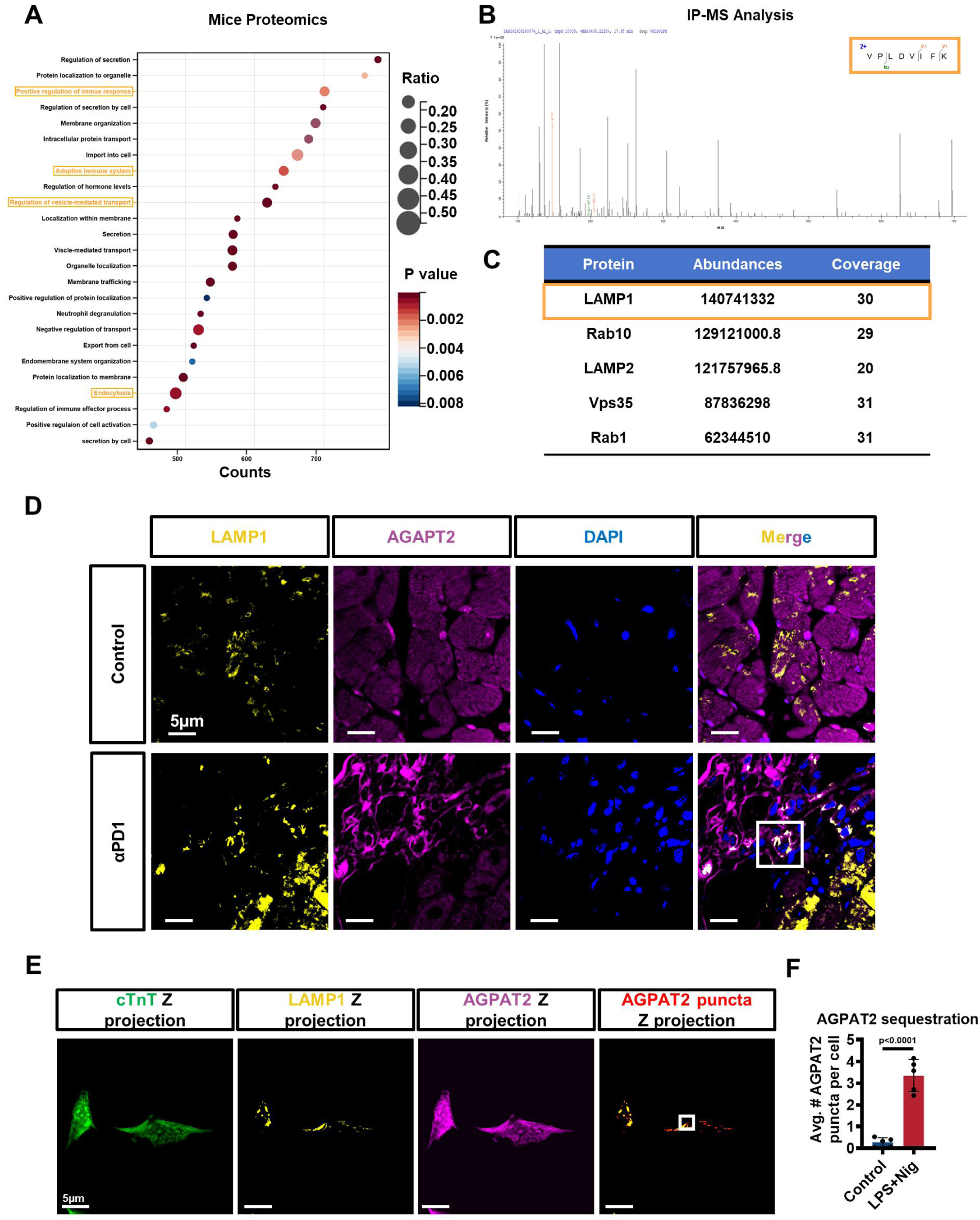
AGPAT2 is Mainly Decomposed by Lysosomes in Inflammatory Metabolic Reprogramming. A. The heatmap showing the most significant enrichment proteins in mice. B. Immunoprecipitation mass spectrometry (IP-MS) detected the most characteristic peptide fragment bound to AGPAT2. C. IP-MS detected the top 5 proteins in peptide abundance and coverage from Co-IP samples bound to AGPAT2. D. Representative 2D immunofluorescence colocalization images (marked with square) of mice hearts subjected to αPD1 treatment with staining LAMP1 in yellow, AGPAT2 in magenta, and nuclei in blue (n = 4/group). Scale bar, 5μm. E-F. Representative immunofluorescence images (E) and quantification (F) of HL-1 cardiomyocytes highlighted AGPAT2 puncta localization (marked with square) outside the cTnT signal and colocalized within LAMP1-positive compartments, with staining cTnT in green, LAMP1 in yellow, AGPAT2 in magenta (n = 5/group, unpaired two-tailed Student’s t-test). Scale bar, 5μm. Data were presented as mean ± standard deviation (SD). AGPAT2, 1-acylglycerol-3-phosphate O-acyltransferase 2; αPD1, anti-PD1 (PD1 inhibitor); Avg.#AGPAT2 puncta per cell, average AGPAT2 positive puncta per cell; cTnT, cardiac troponin T; LAMP1, lysosomal-associated membrane protein 1; LAMP2, lysosomal-associated membrane protein 2; Rab1, ras-related protein Rab 1; Rab10, ras-related protein Rab 10; Vps35, vacuolar protein sorting 35.

**Supplemental Figure 4.**
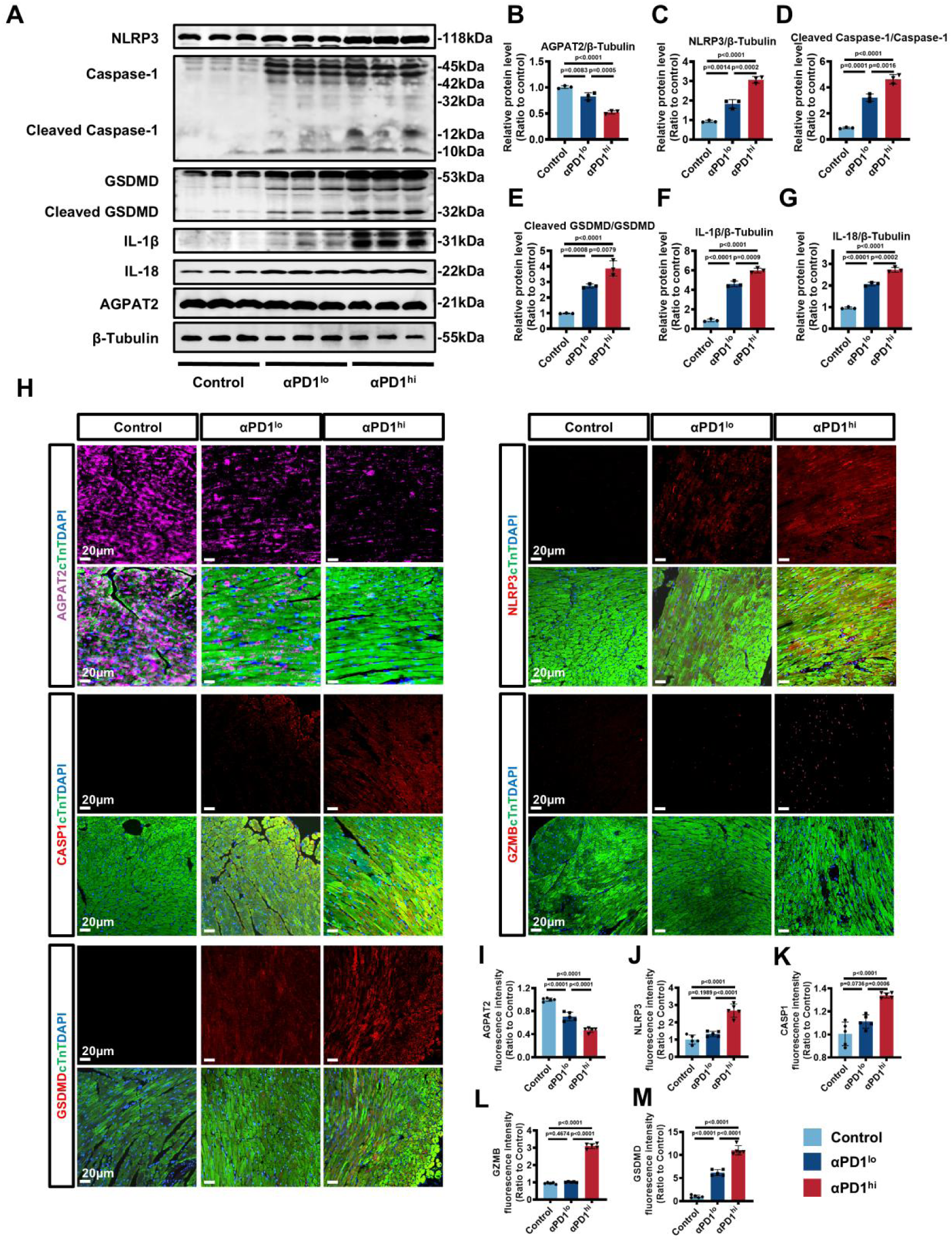
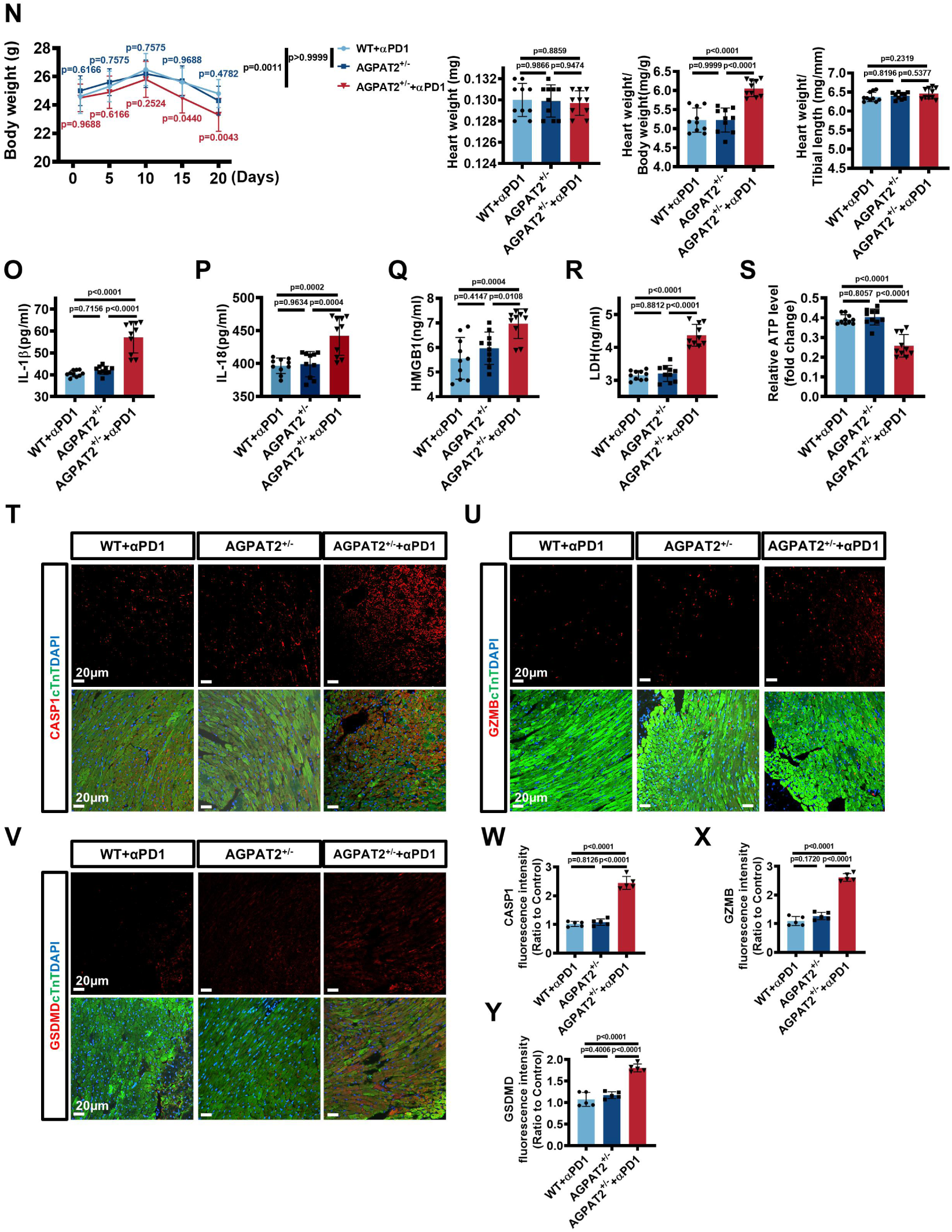
Downregulation of AGPAT2 Expression Leads to Myocardial Pyroptosis. A-G. Representative bands (A) and quantification (B-G) of AGPAT2 and pyroptosis pathway-associated proteins in control, αPD1^lo^ and αPD1^hi^ groups (n = 10/group, one-way ANOVA with post-hoc Tukey HSD test). H-M. Representative immunofluorescence images (H) and quantification of AGPAT2 (I), NLRP3 (J), Caspase 1 (K), Granzyme B (L) and GSDMD (M) in each group, with staining AGPAT2 in magenta, NLRP3, Caspase 1, Granzyme B and GSDMD in red, respectively, cTnT in green and nuclei in blue in each group (n = 10/group, one-way ANOVA with post-hoc Tukey HSD test). Scale bar, 20μm. K. Body weight changes during the αPD1 treatment (n = 10/group, two-way ANOVA with post-hoc Tukey HSD test) and heart weight, heart weight to body weight ratio, heart weight to tibial length ratio (n = 10/group, one-way ANOVA with post-hoc Tukey HSD test) in each group. L-P. The serum IL-1β, IL-18, HMGB1, LDH levels were measured by ELISA assays, and the ATP level in the heart was measured by ATP assay in each group (n = 10/group, one-way ANOVA with post-hoc Tukey HSD test). Q-V. Representative immunofluorescence images (Q-S) and quantification of Caspase 1 (T), Granzyme B (U) and GSDMD (V) in each group, with staining Caspase 1, Granzyme B and GSDMD in red, respectively, cTnT in green and nuclei in blue of three groups (n = 10/group, one-way ANOVA with post-hoc Tukey HSD test). Scale bar, 20μm. Data were presented as mean ± standard deviation (SD). αPD1, anti-PD1 (PD1 inhibitor); αPD1^hi^, high dose of anti-PD1; αPD1^lo^, low dose of anti-PD1; AGPAT2, 1-acylglycerol-3-phosphate O-acyltransferase 2; ATP, adenosine triphosphate; CASP1, Caspase-1, Cysteine-aspartic proteases 1; cleaved Gasdermin D, N-terminal gasdermin domain; cTnI, cardiac troponin I; GSDMD, Gasdermin D; GZMB, Granzyme B; HMGB1, high-mobility group box 1; IgG, immunoglobulin G; IL-1β, Interleukin-1β; IL-18, Interleukin-18; LDH, lactate dehydrogenase; NLRP3, NOD-like receptor pyrin domain-containing protein 3.

**Supplemental Figure 5.**
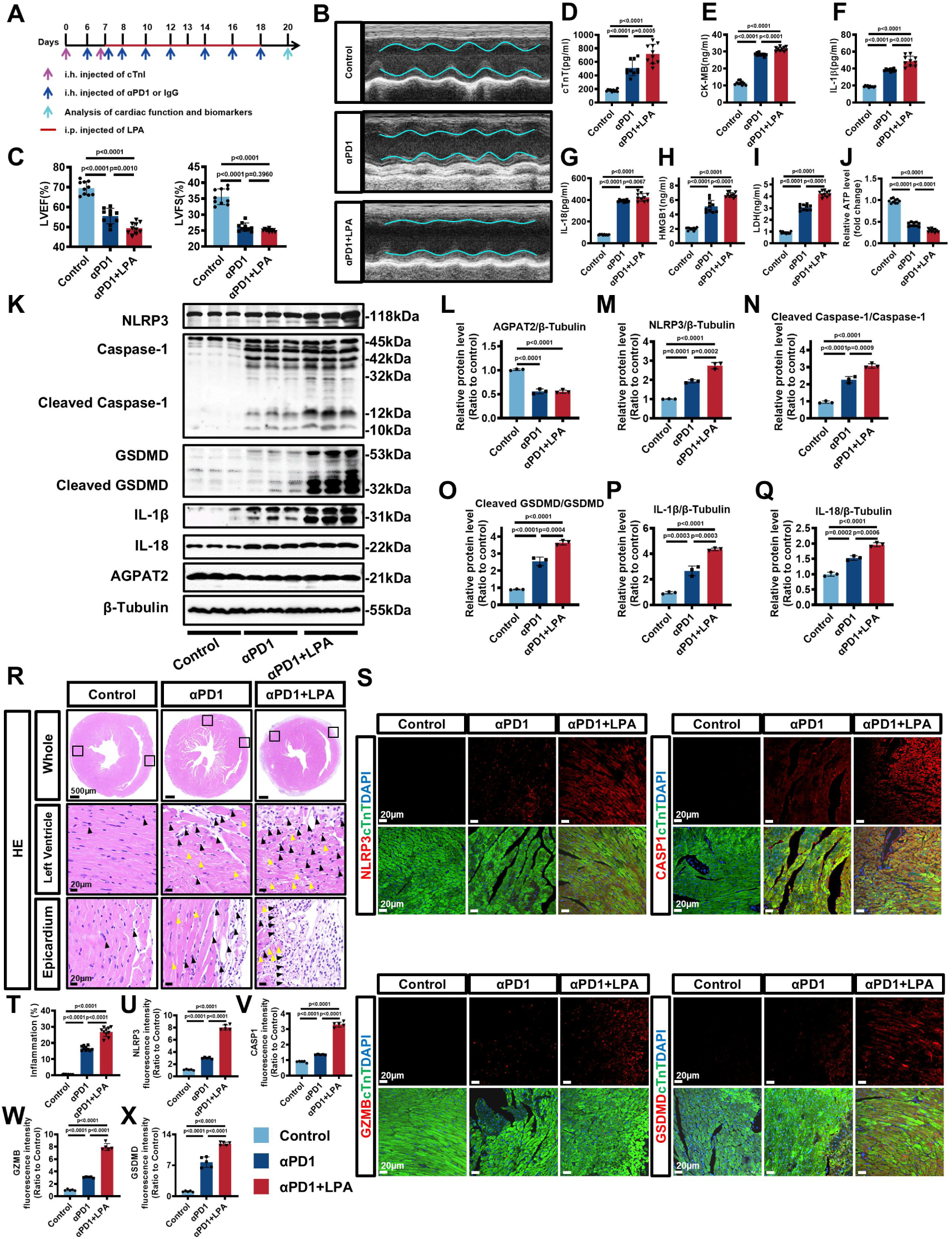
The Accumulation of LPA Exacerbates Myocardial Pyroptosis. A. Experimental scheme depicted strategies in Control, αPD1 and αPD1+LPA groups, the latter with adding LPA to imitate LPA accumulation. B-C. Representative M-mode echocardiographic images (B) and quantification of LVEF, LVFS (C) in each group (n = 10/group, one-way ANOVA with post-hoc Tukey HSD test). D-J. The serum cTnT, CK-MB, IL-1β, IL-18, HMGB1, LDH levels were measured by ELISA assays, and the ATP level in the heart was measured by ATP assay in each group (n = 10/group, one-way ANOVA with post-hoc Tukey HSD test). K-Q. Representative bands (K) and quantification (L-Q) of AGPAT2 and pyroptosis pathway-associated proteins in each group (n = 10/group, one-way ANOVA with post-hoc Tukey HSD test). R, T. Representative HE staining (R) and quantification of inflammation infiltration area (T) in each group (n = 10/group, one-way ANOVA with post-hoc Tukey HSD test). The left ventricle myocardium and right ventricle epicardium in each group are shown respectively. The immune cells in each group are marked with black arrows, and the disordered cardiomyocytes are marked with yellow arrows. S, U-X. Representative immunofluorescence images (S) and quantification of NLRP3(U), Caspase 1 (V), Granzyme B (W) and GSDMD (X) in myocardial biopsy sections from mice, with staining NLRP3, Caspase 1, Granzyme B and GSDMD in red, respectively; cTnT in green and nuclei in blue in each group (n = 10/group, one-way ANOVA with post-hoc Tukey HSD test). Scale bar, 20μm. Data were presented as mean ± standard deviation (SD). αPD1, anti-PD1 (PD1 inhibitor); AGPAT2, 1-acylglycerol-3-phosphate O-acyltransferase 2; ATP, adenosine triphosphate; CASP1, Caspase 1, Cysteine-aspartic proteases 1; CK-MB, creatine kinase-myocardial band; cleaved Gasdermin D, N-terminal gasdermin domain; cTnI, cardiac troponin I; cTnT, cardiac troponin T; GSDMD, Gasdermin D; GZMB, Granzyme B; HE staining, Hematoxylin and Eosin staining; HMGB1, High-mobility group box 1; IgG, immunoglobulin G; IL-1β, Interleukin-1β; IL-18, Interleukin-18; LDH, lactate dehydrogenase; LPA, lysophosphatidic acid; LVEF, left ventricular ejection fraction; LVFS, left ventricular fractional shortening; NLRP3, NOD-like receptor pyrin domain-containing protein 3.

**Supplemental Figure 6.**
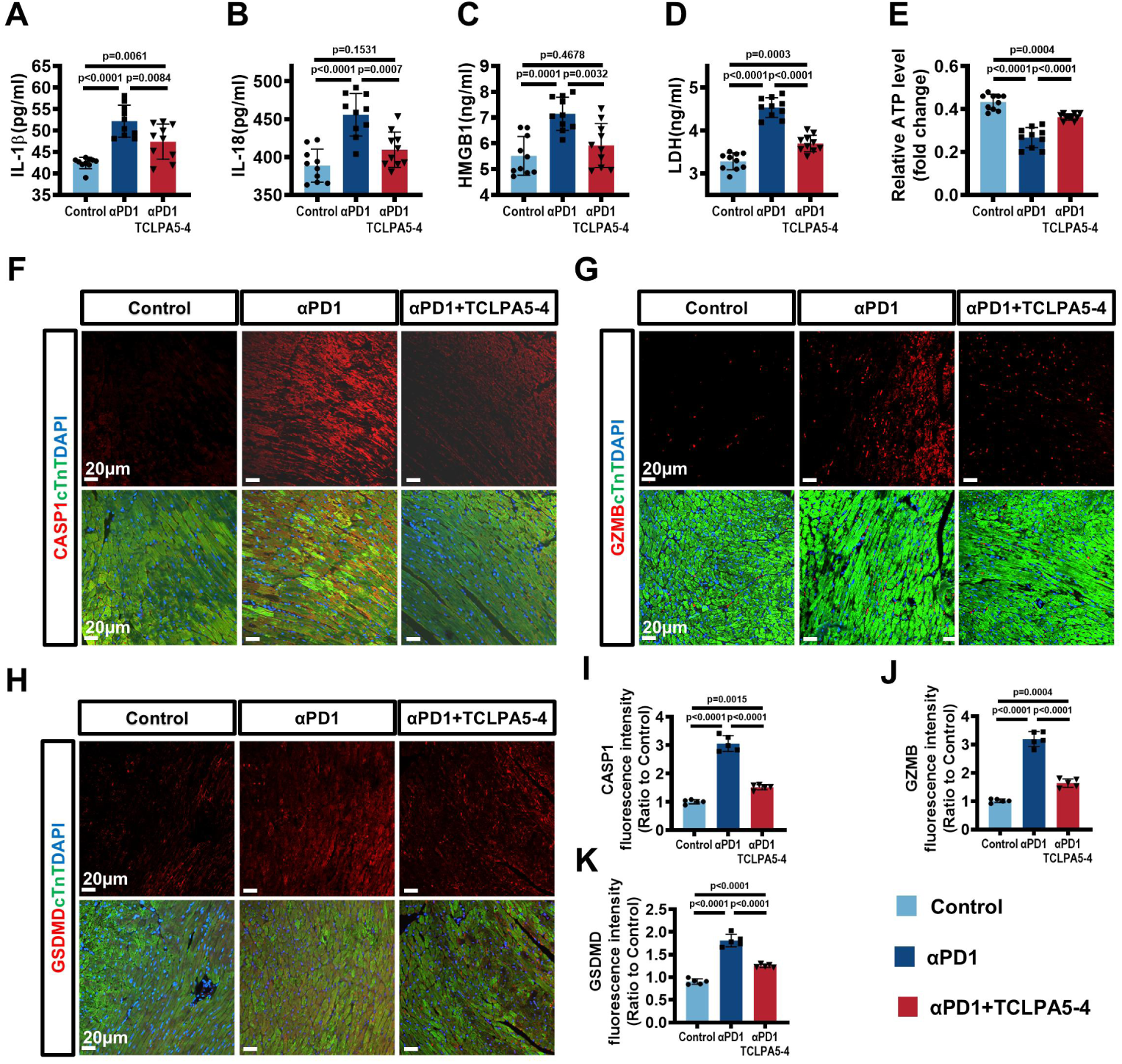
TCLPA5-4 Relieves Pyroptotic Myocardial Injury in AGPAT2 Deficiency Mice. A-E. The serum IL-1β, IL-18, HMGB1, LDH levels were measured by ELISA assays, and the ATP level in the heart was measured by ATP assay in AGPAT2^+/-^ ICIAM mice treated with or without TCLPA5-4 (n = 10/group, one-way ANOVA with post-hoc Tukey HSD test). F-I. Representative immunofluorescence images (F) and quantification of Caspase 1 (G), Granzyme B (H) and GSDMD (I) in myocardial biopsy sections from mice, with staining Caspase 1, Granzyme B and GSDMD in red, respectively; cTnT in green and nuclei in blue in each group (n = 10/group, one-way ANOVA with post-hoc Tukey HSD test). Scale bar, 20μm. Data were presented as mean ± standard deviation (SD). ATP, adenosine triphosphate; CASP1, Caspase 1, Cysteine-aspartic proteases 1; GSDMD, Gasdermin D; GZMB, Granzyme B; HMGB1, high-mobility group box 1; IL-1β, Interleukin-1β; IL-18, Interleukin-18; LDH, lactate dehydrogenase.

**Supplemental Figure 7.**
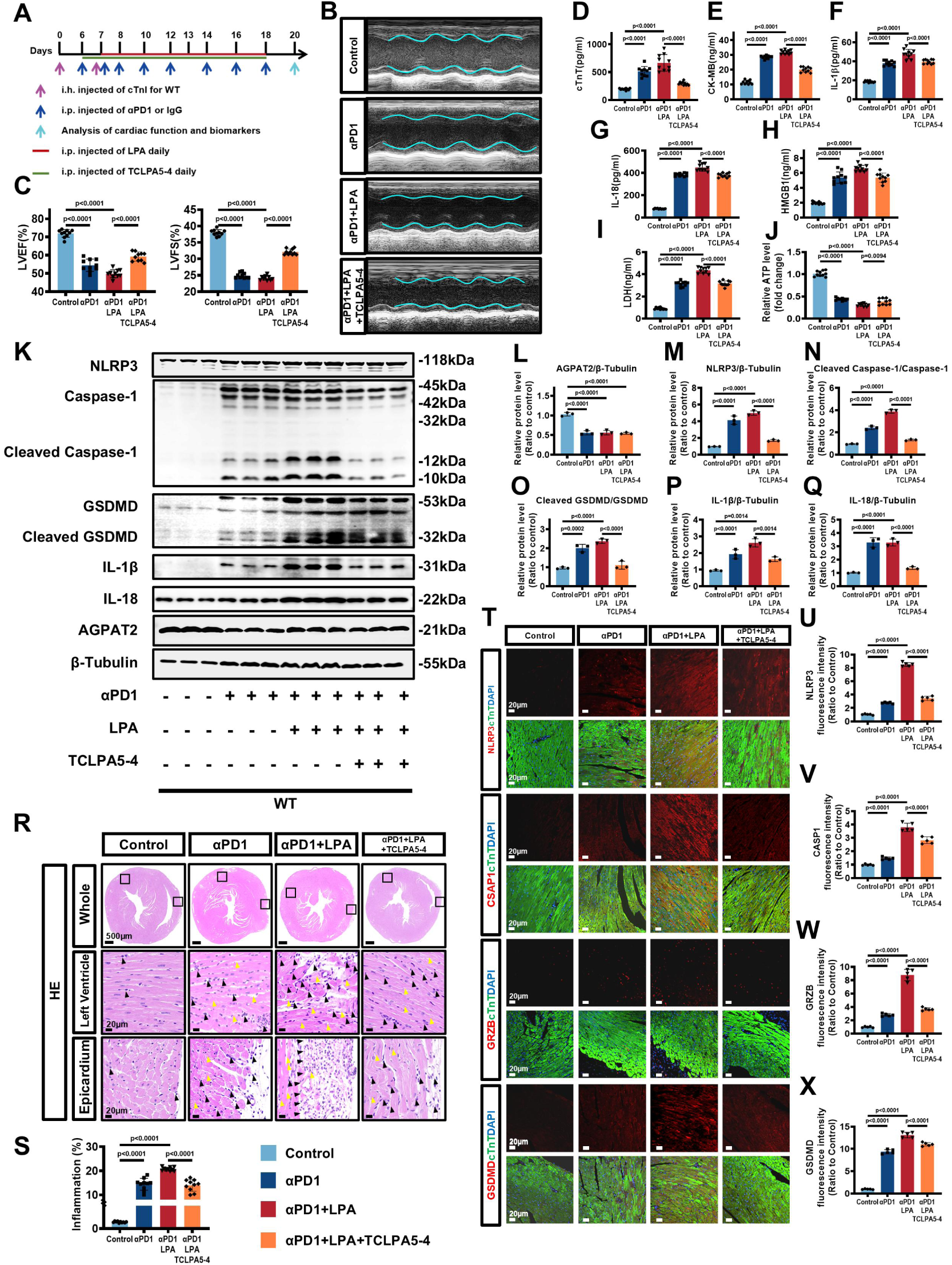
TCLPA5-4 Improves ICIAM in Wild Type Mice via LPA-LPAR5 Blockade. A. Experimental scheme depicted the treatment effect of TCLPA5-4 in WT ICIAM mice. B-C. Representative M-mode echocardiographic images (B) and quantification of LVEF, LVFS (C) in each group (n = 10/group, one-way ANOVA with post-hoc Tukey HSD test). D-J. The serum cTnT, CK-MB, IL-1β, IL-18, HMGB1, LDH levels were measured by ELISA assays, and the ATP level in the heart was measured by ATP assay in each group (n = 10/group, one-way ANOVA with post-hoc Tukey HSD test). K-Q. Representative bands (K) and quantification (L-Q) of AGPAT2 and pyroptosis pathway-associated proteins in each group (n = 10/group, one-way ANOVA with post-hoc Tukey HSD test). R, S. Representative HE staining (R) and quantification of inflammation infiltration (S) in each group (n = 10/group, one-way ANOVA with post-hoc Tukey HSD test). The left ventricle myocardium and right ventricle epicardium in each group are shown respectively. The immune cells in each group are marked with black arrows, and the disordered cardiomyocytes are marked with yellow arrows. T-X. Representative immunofluorescence images (T) and quantification of NLRP3 (U), Caspase 1 (V), Granzyme B (W) and GSDMD (X) in myocardial biopsy sections from mice, with staining NLRP3, Caspase 1, Granzyme B and GSDMD in red, respectively; cTnT in green and nuclei in blue in each group (n = 10/group, one-way ANOVA with post-hoc Tukey HSD test). Scale bar, 20μm. Data were presented as mean ± standard deviation (SD). αPD1, anti-PD1 (PD1 inhibitor); AGPAT2, 1-acylglycerol-3-phosphate O-acyltransferase 2; ATP, adenosine triphosphate; CASP1, Caspase-1, Cysteine-aspartic proteases 1; CK-MB, creatine kinase-myocardial band; cleaved Gasdermin D, N-terminal gasdermin domain; cTnT, cardiac troponin T; GSDMD, Gasdermin D; GZMB, Granzyme B; HE staining, Hematoxylin and Eosin staining; HMGB1, High-mobility group box 1; IgG, immunoglobulin G; IL-1β, Interleukin-1β; IL-18, Interleukin-18; LDH, lactate dehydrogenase; LVEF, left ventricular ejection fraction; LVFS, left ventricular fractional shortening; NLRP3, NOD-like receptor pyrin domain-containing protein 3; TCLPA5-4, inhibitor of LPAR5; WT, wild type.

**Supplemental Figure 8.**
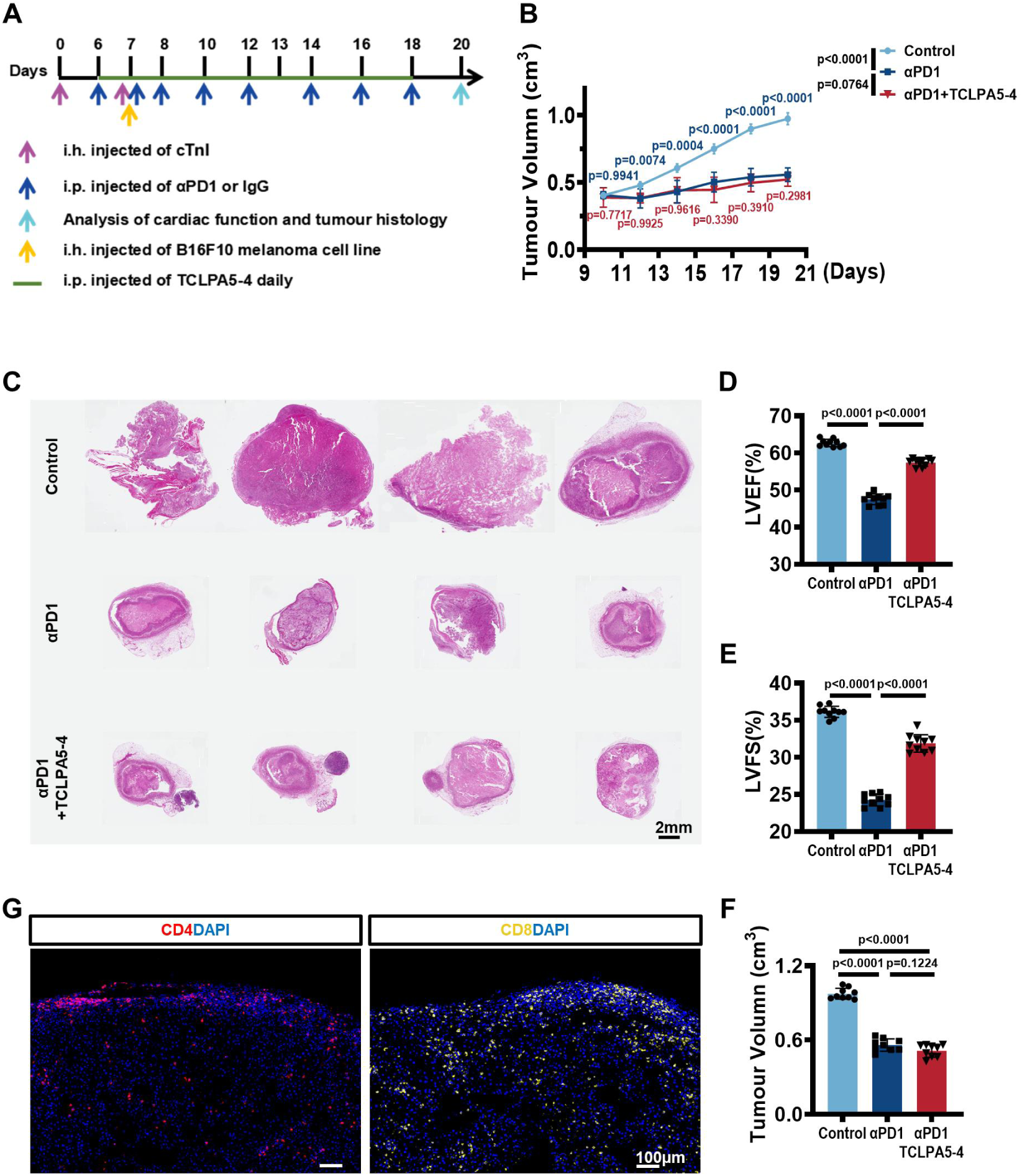
TCLPA5-4 Has no Interference on Anti-Tumor Effects of PD1 Inhibitor. A. Experimental scheme depicted the treatment effect of αPD1 and TCLPA5-4 in melanoma AGPAT2+/- ICIAM mice. B. Tumor growth curve. (n = 10/group, two-way ANOVA with post-hoc Tukey HSD test). Scale bar, 2mm. C. Tumor morphology visualized by HE staining. (Representative data from n = 4/group, total sample size, n = 10/group). D, E. M-mode echocardiographic quantification of LVEF, LVFS in each group (n = 10/group, one-way ANOVA with post-hoc Tukey HSD test). F. Tumor volume at the end of the treatment period in each group (n = 10/group, one-way ANOVA with post-hoc Tukey HSD test). G. Representative immunofluorescence images of intratumoral CD4+ and CD8+T cell recruitment and CD8+T cell accumulation at the tumor-stroma interface after 12 days of indicated therapy as a sign for an anti-tumor immune reaction, with staining CD4 in red, CD8 in yellow and nuclei in blue in each group (n = 10/group, one-way ANOVA with post-hoc Tukey HSD test). Scale bar, 100μm. Data were presented as mean ± standard deviation (SD). AGPAT2, 1-acylglycerol-3-phosphate O-acyltransferase 2; AGPAT2^+/-^, heterozygous AGPAT2 knockout; αPD1, anti-PD1 (PD1 inhibitor); cTnI, cardiac troponin T; HE staining, Hematoxylin and Eosin staining; ICIAM, immune checkpoint inhibitor-associated myocarditis; IgG, immunoglobulin G; LVEF, left ventricular ejection fraction; LVFS, left ventricular fractional shortening; TCLPA5-4, inhibitor of LPAR5.

**Supplemental Figure 9.**
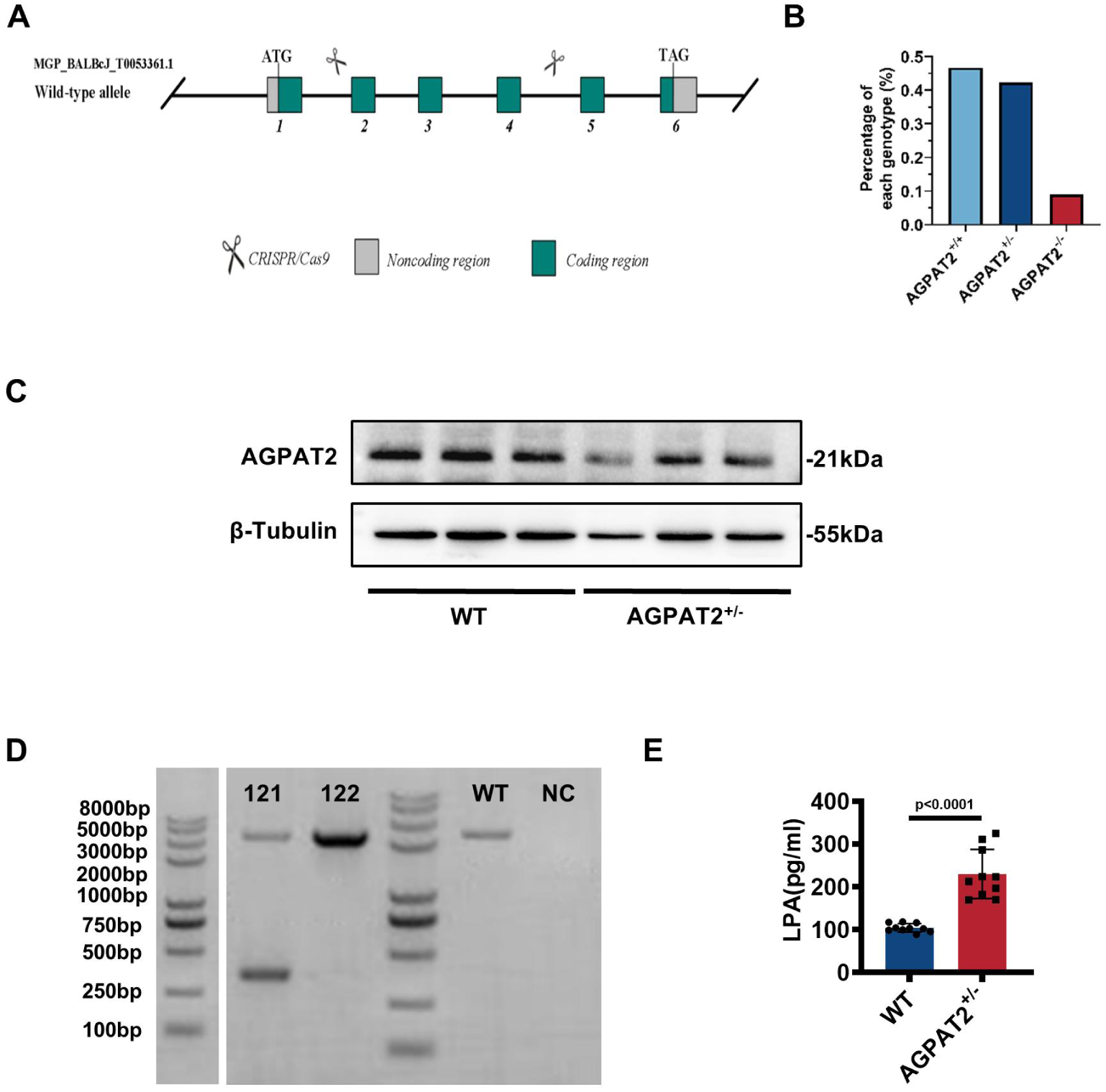
Construction and Characteristics of Heterozygous AGPAT2 Knockout Mice. A. Experimental scheme depicted the construction of AGPAT2 knockout mice using CRISPR-Cas9 system. B. Survival ratio of mice from the WT (AGPAT2^+/+^), AGPAT2 heterozygous knockout (AGPAT2^+/-^) and AGPAT2 homozygous knockout (AGPAT2^-/-^) genotypes. C. Representative bands of AGPAT2 from WT and AGPAT2^+/-^ mice hearts (Representative data from n = 3/group, total sample size, n= 10/group). D. AGPAT2 gDNA agarose gel electrophoresis results: 1.5% agarose gel; KO band: ∼367bp; WT band: 2186bp; 121: AGPAT2^+/-^ mice; 122: AGPAT2^+/+^ mice; WT: wild type BALB/cJGpt mice; NC: blank control. E.Theserum LPA levels measured by ELISA assay (n= 10/group, unpaired two-tailedStudent’s t-test). Data were presented as mean ± standard deviation (SD). AGPAT2, 1-acylglycerol-3-phosphate O-acyltransferase 2; CRISPR-Cas9, Clustered regularly interspaced short palindromic repeats-CRISPR associated protein 9; gDNA, genomic DNA; KO, knockout; LPA, Lysophosphatidic acid; WT, wild type.

## Supplemental Methods

### Generation of the ICIs-related myocarditis model and melanoma mouse model

Wild type (WT) BALB/cJGpt mice were purchased from Vital River Laboratories (Shanghai, China), AGPAT2 heterozygous knockout BALB/cJGpt mice (AGPAT2^+*/-*^, Stock No. T012747) from Gempharmatech (Nanjing, China). All mice were maintained in a temperature-controlled facility with a 12-hour light/dark cycle at 22±2°C and fed with free access to standard mouse chow and tap water.

Due to high homozygous lethality(**45**) (Figure S7B), AGPAT2 heterozygous knockout mice were generated by CRISPR/Cas9 system. Briefly, 2 single guide RNAs (sgRNAs)-targeting the introns on both sides of the targeted region were respectively constructed and transcribed in *vitro*. Then Cas9 mRNA and sgRNA will be co-injected into zygotes. Thereafter, the zygotes were transferred into the oviduct of pseudopregnant ICR females. And F0 mice was birthed after 19-21 days of transplantation, all the offspring of ICR females (F0 mice) were identified by PCR and sequencing of tail DNA (Figure S7D). Finally, crossing F0 mice with BALB/cJGpt mouse to build up heterozygous mice. Exon 2 and 4 of *Agpat2* were used as knockout region. This kind of mice had more accumulation of LPA due to AGPAT2 deficiency (Figure S7E).

To induce ICIs-associated myocarditis (ICIAM) in the tumor-free mice, 6-week-old male mice were randomly assigned to 3 groups. The control group were treated with 250ug rat immunoglobulin G 2a (IgG 2a, clone: 2A3, InVivoMab, USA) via intraperitoneal injection at day 6, 7 and 8, and day 1 followed by every second day for 8 times. The low-αPD1 (αPD1^lo^) group were first immunized subcutaneously on day 0 and day 7, respectively, with a solution of 250ug murine cardiac troponin I peptide (cTnI, Sangon Biotech, China) diluted in complete Freund’s adjuvant (Sigma, USA). Then, treated with 250ug anti-mouse PD1 antibodies (αPD1, clone: RMP1-14, InVivoMab, USA) intraperitoneally at day 6, 7, 8, 10, 12 for 5 times. The high-αPD1 (αPD1^hi^) group received the same cTnI and complete Freund’s adjuvant immunization as the αPD1^lo^ group, followed by 250ug αPD1 for 8 times.

To induce ICIAM in the tumor-bearing mice, 6-week-old AGPAT2^+/-^ male mice were subcutaneously transplanted with 5×10^5^/mouse melanoma cells B16F10 (CH3 BioSystems, USA) between passages 3-4 on day 7. Likewise, cTnI and complete Freund’s adjuvant, αPD1 or IgG 2a were injected with the same as used in the control group or the αPD1^hi^ group in the tumor-free mice.

Body weights were evaluated every second day. On day 13 in the αPD1^lo^ group and day 20 in the αPD1^hi^ group for tumor-free mice, day 20 for tumor-bearing mice, echocardiography was performed, tibial length was measured and heart tissues were obtained from mice euthanized using deep isoflurane (5%) anesthesia, perfused and rinsed in ice-cold phosphate buffer saline (PBS, Servicebio Biotech, China), weighted and snap-frozen in liquid nitrogen. The experimental design paradigms are shown in the corresponding figures.

No mice were excluded from the study except for failure of the subcutaneous transplantation of melanoma cells. All animal experiments were conducted in accordance with “the Animal Research: Reporting of In Vivo Experiments guidelines (ARRIVE guidelines, version 2.0)” and were approved by the Animal Care and Use Committee of Zhongshan Hospital, Fudan University (2021–09). In animal studies, the investigators were blind to treatment/genotype groups during the experiment and quantification. Mice were randomly allocated to each group using a random number generator. Sample size was determined by power calculations based on data from previous studies with an α level of 0.05 and power of 0.80. The number of animals for different experiments is shown in the footnotes of each figure.

### In vivo treatments

To examine the role of lysophosphatidic acid (LPA), mice were injected intraperitoneally with LPA (Cat No. L7260, Sigma) with a dose of 1mg/kg (dissolved in DMSO) once daily from day 7 for tumor-free mice for 12 consecutive days.

To assess the blocking effect of lysophosphatidic acid receptor (LPAR), mice were given TCLPA5-4 (Cat No. HY-107615, MEDCHEMEXPRESS) with a dose of 10mg/kg (dissolved in DMSO) once daily by intraperitoneal injection from day 6 for tumor-free and tumor-bearing mice for 13 consecutive days.

### Isolation of neonatal rat ventricular cardiomyocytes (NRVCs)

Neonatal rat ventricular cardiomyocytes (NRVCs) were isolated and cultured as previously described^45^. Briefly, hearts of 1- to 3-day-old pups were dissected and atria were removed. Ventricles were minced and digested repeatedly with 0.125% trypsin EDTA (Gibco, USA) at 37℃ for 30 second to 1 minute per cycle. When the cell supernatant became cloudy, the supernatant was neutralized by Dulbecco’s modified Eagle’s medium (DMEM, Gibco, USA) High Glucose, with 10% Fetal Bovine Serum (FBS, Gibco, USA). After digesting for 20-30 cycles, supernatant containing isolated cells were collected. The obtained cells were centrifuged at 25℃, 1000 rpm for 5 minutes to separate non-myocardial cells and cultured at 37℃ in a humidified 5% CO2 incubator. One hour later, the supernatant containing suspended cardiomyocytes was transferred into other plates to reduce fibroblast contamination. Next, cell supernatant was passed through a 100 μm strainer before NRVCs being cultured in DMEM High Glucose supplemented with 10% FBS, and 1% penicillin/streptomycin (P/S) (Gibco, USA) for 2-3 days. Cardiomyocytes were used in experiments after the cells formed a confluent monolayer and contracted in synchrony.

### Isolation of adult mouse cardiomyocytes

Chest cavity of anaesthetised mouse was opened to below diaphragm to fully expose heart. Descending aorta and inferior vena cava were cut, then 7 ml EDTA buffer was injected into apex of right ventricle. Clamp the aorta and removed heart by cutting behind clamp.

Clamped heart is submerged in 60 mm dish of EDTA buffer. 10 ml EDTA buffer isinjected into apex of left ventricle (LV). Heart is transferred to dish of perfusion buffer. 3 ml perfusion buffer is injected into apex of LV. Then heart is transferred to dish of collagenase buffer. 30-40 ml collagenase buffer is injected into apex of LV, until digestion is apparent.

Clamp is removed, heart tissue is then pulled gently into ∼1mm^3^ pieces using forceps, and dissociated by gentle pipetting. After that, stop buffer is added, cell suspension is passed through 100 μm strainer and cardiomyocytes are gravity settle for 20 min. Cardiomyocytes then undergo two further rounds of gravity settling to achieve a pure population.

### Cell Treatment and virus infection

HL-1 mouse cardiomyocytes were purchased from Type Culture Collection of the Chinese Academy of Sciences (Shanghai, China), used between passages 3-4 (same passage cells in each experiment) and cultured in DMEM High Glucose with 10% FBS and 1% P/S (Hyclone, USA) at 37℃ in a humidified 5% CO2 incubator. Cells were seeded on a 6-or 96-well plate or a confocal dish 24 hours before experiments.

To explore occurrence of AGPAT2-inflammatory metabolic reprogramming (AGPAT2-IMRP), HL-1s were pre-treated for 1 hour with DMSO, 1nM Bafilomycin A1 (BafA1, lysosomal inhibitor, Cat No. D1515, Sigma) or 1nM MG132 (ubiquitination inhibitor, Cat No. M8699, Sigma), respectively. Then followed by 6-hour stimulation with lipopolysaccharide (LPS, Cat No. HY-D1056, MedChenExpress) at 1μg/ml with Nigericin (Nig, Cat No. HY-100381, MedChenExpress) at 15μM.

The mouse AGPAT2 plasmid (3Flag-AGPAT2 plasmid) was purchased from (Biotech, China). To demonstrate the interaction between AGPAT2 and LAMP1, HL-1s were transfected with 2.5μg 3-Flag-AGPAT2 plasmid for 24 hours using Lipofectamine 3000 (Cat No. L3000008, Thermo Fisher) according to the manufacturer’s instructions. Then incubated 6 hours with LPS+Nig (1μg/ml+15μM).

To determine the Ic50 of lysophosphatidic acid (LPA), cells were stimulated with LPA (Cat No. L7260, Sigma) at (0, 0.01, 0.1, 0.5, 1, 2.5, 5, 10, 15, 20, 25μM) for 6 hours.

To explore the main pathway of LPA-induced cell death, cells were pre-incubated for 1 hour with PBS, 22µM SPD304 (TNF-α/necrosis inhibitor, Cat No. HY-111255, MedChenExpress), 18µM Z-DEVD-FMK (Caspase-3/apoptosis inhibitor, Cat No. HY-12466, MedChenExpress), 10µM INF39 (NLRP3/pyroptosis inhibitor, Cat No. HY-101868, MedChenExpress) separately, followed by LPA stimulating at 20μM for 6 hours.

To explore the effect of knockdown/overexpression of AGPAT2 on LPA-induced pyroptotic injury of cardiomyocytes, HL-1s were firstly transfected with AGPAT2 siRNA (siAGPAT2), scrambled siRNA (NC-AGPAT2), AGPAT2 overexpression plasmid (OE-AGPAT2) or vehicle plasmid (Veh-AGPAT2) (Biotech, China), respectively for 24 hours. And then treated with DMSO or LPS+Nig+LPA (1μg/ml+15μM+20μM) for 6 hours. Furthermore, NRVCs were infected by lentivirus (MOI=150, for 72 hours) as follows: knockdown virus (LV-shAGPAT2), scrambled virus (LV-shNC), overexpression virus (LV-OE-AGPAT2). And then the infected NRVCs were further treated with 6-hour LPS+Nig+LPA (1μg/ml+15μM+20μM).

To explore which of the LPA receptors (LPARs) plays a key protective role in myocardial pyroptosis, experiments with specific inhibitors of various LPARs were carried out first. HL-1s were pre-incubated with DMSO, 10µM Ki16425 (LPAR1-3 inhibitor, Cat No. HY-13285, MedChemExpress) or 5µM TCLPA5-4 (LPAR5 inhibitor, Cat No. HY-107615, MedChemExpress) for 1 hours, separately, before stimulating with LPS+Nig+LPA as described above. To inhibit the function of each LPAR separately, HL-1s were transfected with LPAR1 siRNA (siLPAR1), LPAR2 siRNA (siLPAR2), LPAR3 siRNA (siLPAR3), LPAR5 siRNA (siLPAR5) or scrambled siRNA (NC-LPAR) (Biotech, China) for 24 hours independently, followed by LPS+Nig+LPA treatments and assessment of myocardial pyroptotic injury.

### Echocardiography

Transthoracic echocardiography was performed in mice anesthetized with 2% isoflurane using a 30-MHz high-frequency scan probe (Vevo 2100, VisualSonics, Canada) on day 13 for αPD1^lo^ group and day 20 for other groups. Left ventricular ejection fraction (LVEF), Left ventricular fractional shortening (LVFS), left ventricular anterior wall (LVAW) and left ventricular posterior wall (LVPW) at end-diastole and end-systole were measured as previously described.

### Western Immunoblot

Total proteins isolated from the left ventricle of heart tissues and HL-1 cells were homogenized in an ice-cold radioimmunoprecipitation assay (RIPA) lysis buffer for 30 min with protease and phosphatase inhibitors (Thermo Fisher Scientific, USA) and then centrifuged at 12,000 rpm for 15 minutes at 4°C. The supernatant protein concentration was measured by the bicinchoninic acid method using BCA protein assay kit (Pierce™ BCA Protein Assay Kit, Thermo Fisher Scientific, USA) following the manufacturer’s instructions. Proteins were denatured in 5×SDS-PAGE (sodium dodecyl sulfate-polyacrylamide gel electrophoresis) loading buffer at 100°C for 10 min. Then, 30μg/lane protein was size-fractionated by SDS-PAGE and transferred onto 0.22μm immobilon polyvinylidene difluoride (PVDF) membranes (Merck Millipore, USA). Membranes were cut into different parts according to the molecular weight of each protein and blocked with 5% nonfat milk (BD Biosciences, USA) for 2 hours. Each membrane was incubated with primary antibody at 4°C overnight: anti-NLRP3 (1:1000, Abcam, UK), anti-pro Caspase-1+p10+p12 (p10 and p12 are two kinds of cleaved Caspase-1) (1:1000, Abcam, UK), anti-Gasdermin D (1:1000, CST, USA), anti-cleaved Gasdermin D (1:1000, CST, USA), anti-IL-1β (1:1000, CST, USA), anti-IL-18 (1:1000, CST, USA), anti-AGPAT2 (1:1000, Novus, USA), anti-LAMP1 (1:1000, DSHB, USA), anti-β-tubulin (1:5000, CST, USA). The membranes were thoroughly washed for the second day by TBST (Sangon Biotech, China) and incubated for 1 hour in the dark with secondary antibodies, DylightTM800 4XPEG-conjugated goat anti-rabbit IgG (H+L), goat anti-mouse IgG (H+L) and goat anti-rat IgG (H+L) (1:10,000, CST, USA). The proteins were visualized using an ECL Western blotting detection system #WBKLS0500 (MilliporeSigma, USA). The density of each band was quantified by densitometric analysis with Image J software (Image Processing and Analysis in Java, National Institutes of Health, USA). A list of antibodies sources and applications is available in the associated table (Supplementary Table 1).

### Real-time quantitative PCR

Trizol Reagent (TRIzol™, Cat No.15596018CN, Thermo Fisher Scientific, USA) was used to extract total RNA and 1μg RNA was reverse transcribed by the PrimeScript™ RT Master Mix (Perfect Real Time) (Cat No. RR036, TaKaRa, China) for the synthesis of cDNA. Real-time quantitative PCR (RT-qPCR) was performed by SYBR Green PCR Master Mix (Cat No. 4368577, Thermo Fisher Scientific, USA) in the ABI 7300 Detection System (Thermo Fisher Scientific, USA). β-tubulin was employed as the endogenious reference gene. Relative gene expression was calculated using 2^-ddCt^ method. The primers are listed in Supplementary Table 2 (Sangon Biotech, China).

### Enzyme-Linked Immunosorbent Assay (ELISA)

Mouse and Human plasma samples were detected by ELISA according to the manufacturer’s instructions. The Cardiac troponin T (cTnT) Assay Kit (Cat No. H149-4, Nanjing Jiancheng bioengineering Institute, China), the Creatine kinase MB (CK-MB) isoenzyme Assay Kit (Cat No. H197-1-1, Nanjing Jiancheng bioengineering Institute, China), the IL-1β Assay Kit (Cat No. MLB00C for mouse and Cat No. DLB50 for human, R&D Systems, USA), the IL-18 Assay Kit (Cat No. 7625 for mouse and DL180 for human, R&D Systems, USA), the LPA Assay Kit (Cat No. EIA07496m for mouse and Cat No. EIA07496h for human, Wuhan Xinqidi Biotechnology, China), the HMGB1 Assay Kit (Cat No. EIA06324m for mouse and Cat No. EIA06324h for human, Wuhan Xinqidi Biotechnology, China), the enhanced ATP assay kit (Cat No. S0027, Beyotime, China), the LDH Assay Kit (Cat No. P0393S, Beyotime, China).

### Cell Viability

HL-1 cells were seeded at a concentration of 5×10^3^ cells/well in a 96-well plate. Different treatments were implemented as described above. The cell viabilities of each group were detected using the Cell Counting Kit-8 (CCK-8, Beyotime Biotechnology, China) for measuring the absorbance values at a wavelength of 450 nm, 1 hour after adding the working solution.

### Proteomics, metabolomics and lipidomics

The snap-frozen mice hearts were manually ground in liquid nitrogen and subjected to the biphasic simultaneous protein extraction (SIMPLEX) workflow(**46**). Following phase separation and protein precipitation, the molecular layer was subjected to analysis for TMT Quantitative proteomics by Q Exactive plus Mass Spectrometer.

For label-free metabolomics and targeted lipidomics of human serum, a liquid chromatography coupled to high resolution mass spectrometry setup was employed while the serum lipidome was investigated by shotgun lipidomics. The proteomics, metabolomics and lipidomics analysis were performed and analyzed by APTBIOtech (Shanghai, China).

### Functionally enrichment and diagram design

The raw data of mass spectrometry were analyzed for inventory identification and quantitative analysis using software XCMS 3.7(**47**), Mascot 2.7(**48**) and Proteome Discoverer 3.1 (**49**).

CAMERA (**50**) was used for annotation of isotopesand adducts. Proteins wi th a 1.2-fold change (≥1.2 or≤0.83) and corresponding adjusted P<0.05 were id entified as target molecules and entered into Metascape database (**51**) for annot ation, visualization and integrated discovery and then subjected to Gene Ontolo gy (**52**) and Kyoto Encyclopedia of Genes and Genomes (**53**) pathway database.

Sangerbox (**54**) platform was used to analyze and make transcript heat maps. Experiment diagrams were created by BioRender (**55**).

### Single-cell transcriptome

The study initiated by obtaining dataset GSE180045 and dataset GSE120567 from the GEO database. The former includes single-cell transcriptome sequencing data from peripheral blood of eight ICIAM patients, while the latter comprises transcriptome sequencing data from cardiac biopsies of nine ICIAM patients. These datasets were collectively analyzed using the Seurat package to reduce dimensionality, cluster the sequencing data, and create a single-cell map. Subsequently, the singleR package was employed to annotate distinct cell clusters by integrating the specific markers associated with each cluster.

### Morphological and histological analysis

The heart and tumor tissues were fixed with 4% paraformaldehyde and embedded in paraffin. The paraffin-embedded hearts or tumors were then sectioned at a thickness of 5μm and stained with hematoxylin-eosin (HE) and Masson trichrome staining reagent according to the manufacturer’s instructions (Servicebio Biotech, China). For measurements, 5 random high-power fields from 5 sections of each heart or tumor were chosen and quantified in a blinded manner. The area of the inflammatory infiltration and fibrosis were evaluated through morphometric analysis of the HE and Masson staining sections respectively. The images were observed and captured with a light microscope (LM, BX51, Olympus, Japan, objective lens magnification of ×2 or ×40; eyepiece magnification of ×10) and processed with Image-Pro Plus software (Version 5.0, Media Cybernetics, USA).

### Immunofluorescent Staining and confocal fluorescence microscopy

Slides of heart sections were deparaffinized, rehydrated, and boiled in Tris-EDTA solution (Beyotime Biotechnology, China) at 95°C for 20 min for antigen retrieval. In addition, the HL-1 or NRVC cells were plated on confocal dish, given different treatments as described previously, and fixed at room temperature (RT) with 4% paraformaldehyde for 20 min. Then the HL-1 cells were treated with 0.1% Triton X-100 for 5 min, and washed three times with PBS. Then, all slides were blocked with 5% goat serum and incubated with primary antibody at 4°C overnight: anti-NLRP3 (1:50, Abcam, UK), anti-cardiac Troponin T (cTnT, 1:500, Abcam, UK), anti-AGPAT2 (1:500, Novus, USA), anti-LAMP1 (1:500, DSHB, USA), anti-CD4 (1:500, CST, USA), anti-CD8 (1:500, CST, USA); anti-CD44 (1:500, CST, USA); anti-Granzyme B (1:100, Proteintech, China), anti-GSDMD (1:50, Proteintech, China) and anti-Caspases-1 (1:100, Abclonal, China). The slides were rewarmed at 37°C for 1 hour, washed by PBS thoroughly and incubated with anti-rabbit IgG (1:1000, Alexa Fluro®488 Conjugate, CST), anti-mouse IgG (1:1000, Alexa Fluro®594 Conjugate, CST) and anti-rat IgG (1:1000, Alexa Fluro®647 Conjugate, CST) at 37°C for 1 hour. Nuclei were counterstained with 4′,6-diamidino-2-phenylindole (DAPI) (62248, Invitrogen, USA). A list of primary antibodies sources and applications is available in the associated table (Supplementary Table 1).

Finally, the stained sections and confocal dish were observed with Olympus FV3000 spinning disk confocal microscope (Olympus, Japan, objective lens magnification of ×40; eyepiece magnification of ×10). 3-dimensional reconstruction of confocal micrographs was performed using Z-axis serious plugin and the fluorescence intensity was processed with Image J software. Operator bias was minimized during image acquisition through selection of fields of view and focal planes based on counterstains that were unrelated to the experimental question.

### Identify AGPAT2 and LAMP1 interaction proteins by mass spectrometry

Five 10 cm dishes HL-1 cells were transfected with 3Flag-AGPAT2 plasmids with Lipofectamine 3000 and continued cultured for 48 hours, followed by 6 hours’ incubation with LPS+Nig (1 μg/ml+15 μM). Then cells were lysed in NP-40 lysis buffer supplemented with protease inhibitor cocktail (HY-K0010, MedChemExpress, China), and were immunoprecipitated with the anti-Flag magnetic beads (82103, CST, USA), Precipitates were separated by SDS-PAGE, and were then stained with Coomassie blue. Subsequently, protein bands were cut into small pieces and were subjected in-gel digestion. Finally, the extracted peptides from the gel pieces were prepared for proteomic data analysis using liquid chromatography-tandem mass spectrometry (LC-MS/MS).

### Co-immunoprecipitation (Co-IP)

Total protein was extracted from HL-1 cells using NP-40 lysis buffer supplemented with protease inhibitor cocktail (HY-K0010, MCE, China). 20uL total protein was extracted as input. Then, cell lysates were incubated and rocked with primary antibody of AGPAT2 (Novus, USA), or LAMP1 (DSHB, USA) or isotype control immunoglobulin G (CST, USA) and 20 mL protein A/G agarose beads (16-266, Millipore, China) at 4°C overnight with rotation. Centrifuged the cell lysates (12,000 rpm for 15 min at 4°C) to collect protein A/G agarose beads, and washed the beads three to five times with NP-40 lysis buffer. Proteins were denatured in 1×SDS-PAGE loading buffer at 100°C for 10 min. Then proteins were subjected to SDS-PAGE and analyzed by Western Blot.

### Clinical data

Tumor patients with suspicious cardiac involvement were enrolled and divided into 3 groups: Group A (No ICI+No myocarditis, n=53), Group B (ICI+No myocarditis, n=54), and Group C (ICI+Myocarditis, n=61). ICIAM was diagnosed according to *the 2022 ESC guidelines for oncocardiology*^24^. Patients with pre-existing cardiovascular diseases were excluded. Myocardial samples were obtained in patients with confirmed ICIAM by myocardial biopsy according to *the 2022 ESC guidelines for oncocardiology***^24^**.

Patients receiving ICIs in our center underwent routine blood sample collection at day 1 before ICIs administration, then on days 7, 14, 21, 35 and 49 after ICIs-treatment. Based on the diagnostic criteria described in *the 2022 ESC guidelines for oncocardiology***^24^**, the presence of ICIAM in patients was determined. In cases of the occurrence of ICIAM, blood samples were additionally collected on the same day for supplemental analysis.

All work involving human subjects was conducted in accordance with the World Medical Association Declaration of Helsinki, obtained informed consent from all participants, and was approved by the Ethics Committee of Zhongshan Hospital, Fudan University (Ethics approval No. B2021-275).

### Tumor Studies

B16F10 cells were subcutaneously implanted before the mice been given various treatments as indicated. Tumor volume (in cm^3^) was measured with calipers and calculated as (width^2^×length)/2 per week for 2 weeks (every two days for 14 days). At the end of the experiment, the tumors were dissected and weighted for analysis.

### Statistical Analysis

Normally distributed measurement data were expressed as mean ± standard deviation (SD); Non normally distributed measurement data are expressed as median (lower quartile, upper quartile); Count data are expressed as number (percentage).

The differences between two groups were compared with unpaired two-tailed Student’s t test, and the differences among multiple groups were compared with one-way or two-way ANOVA with post-hoc Tukey HSD test. Pearson or Fisher chi square test was used to compare the categorical data between the two groups. P<0.05 was considered statistically significant.

Data were analyzed by using SPSS software (version 22.0, SPSS Inc., USA), Prism software (version 8.4.3, GraphPad, USA) and R (version 4.1.0, R Foundation for Statistical Computing, Austria).

### Data Availability Statement

The authors confirm that the data supporting the findings of this study are available within the article or its supplementary materials.

## Supplemental Western Immunoblot Original Images

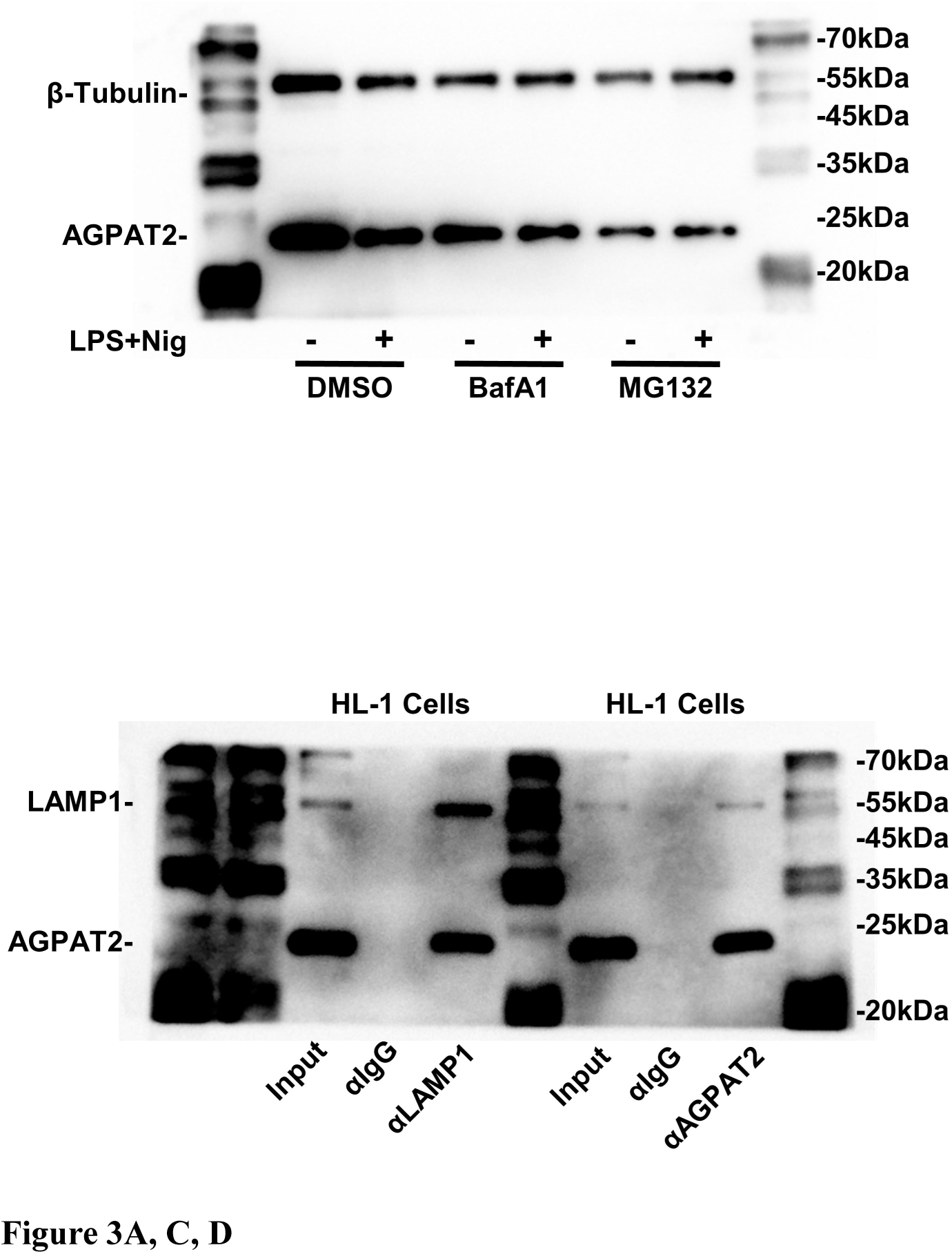

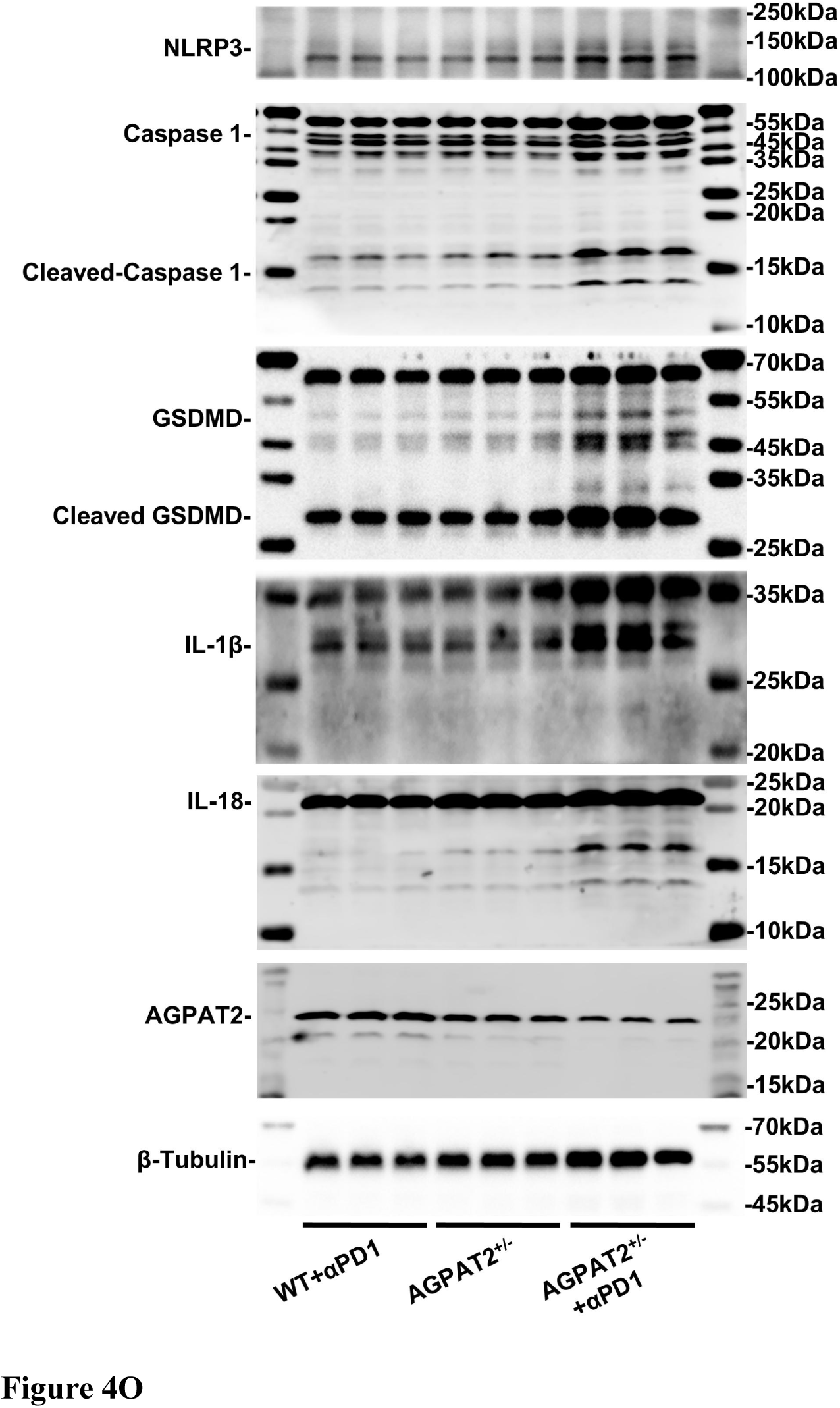

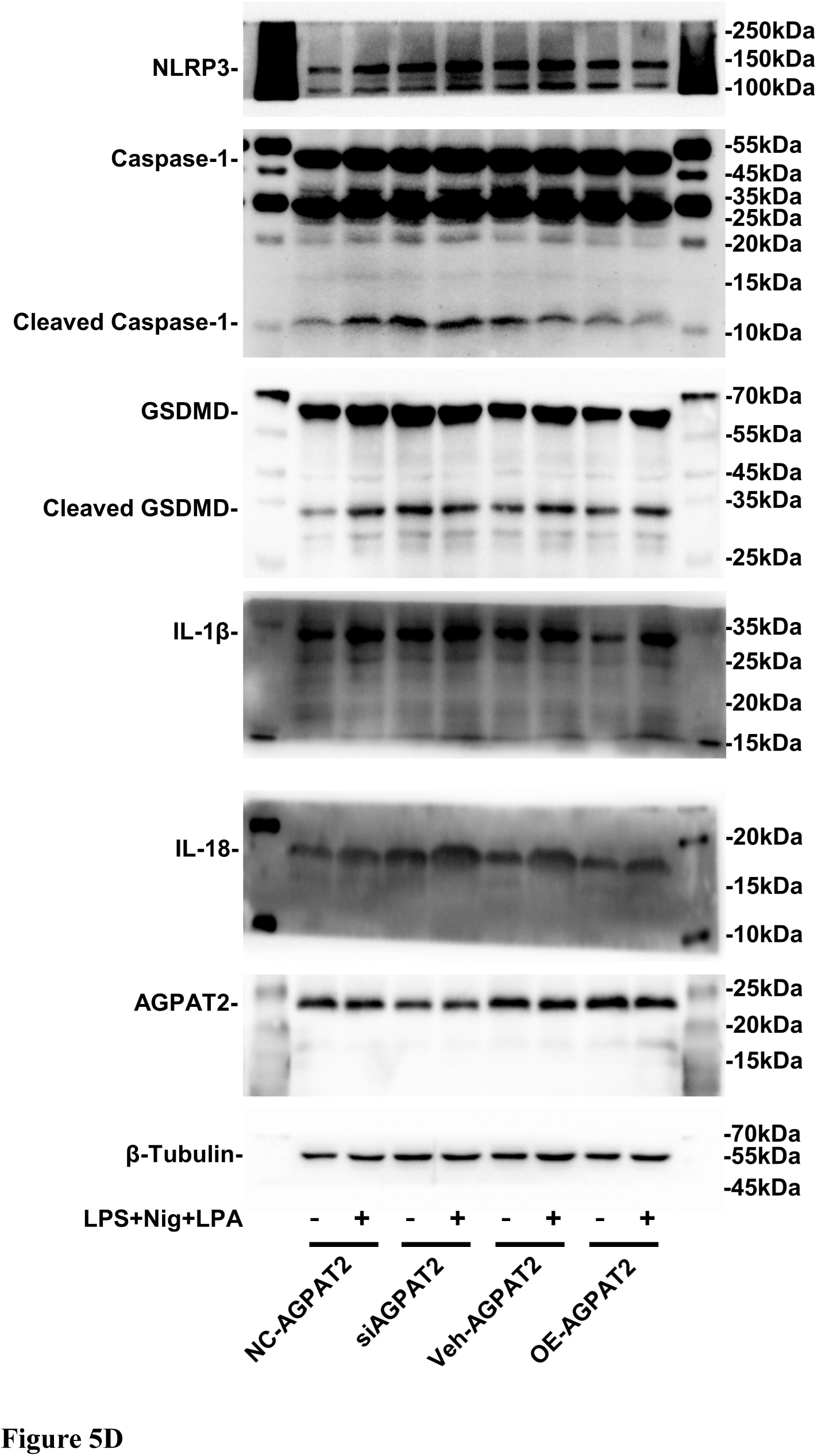

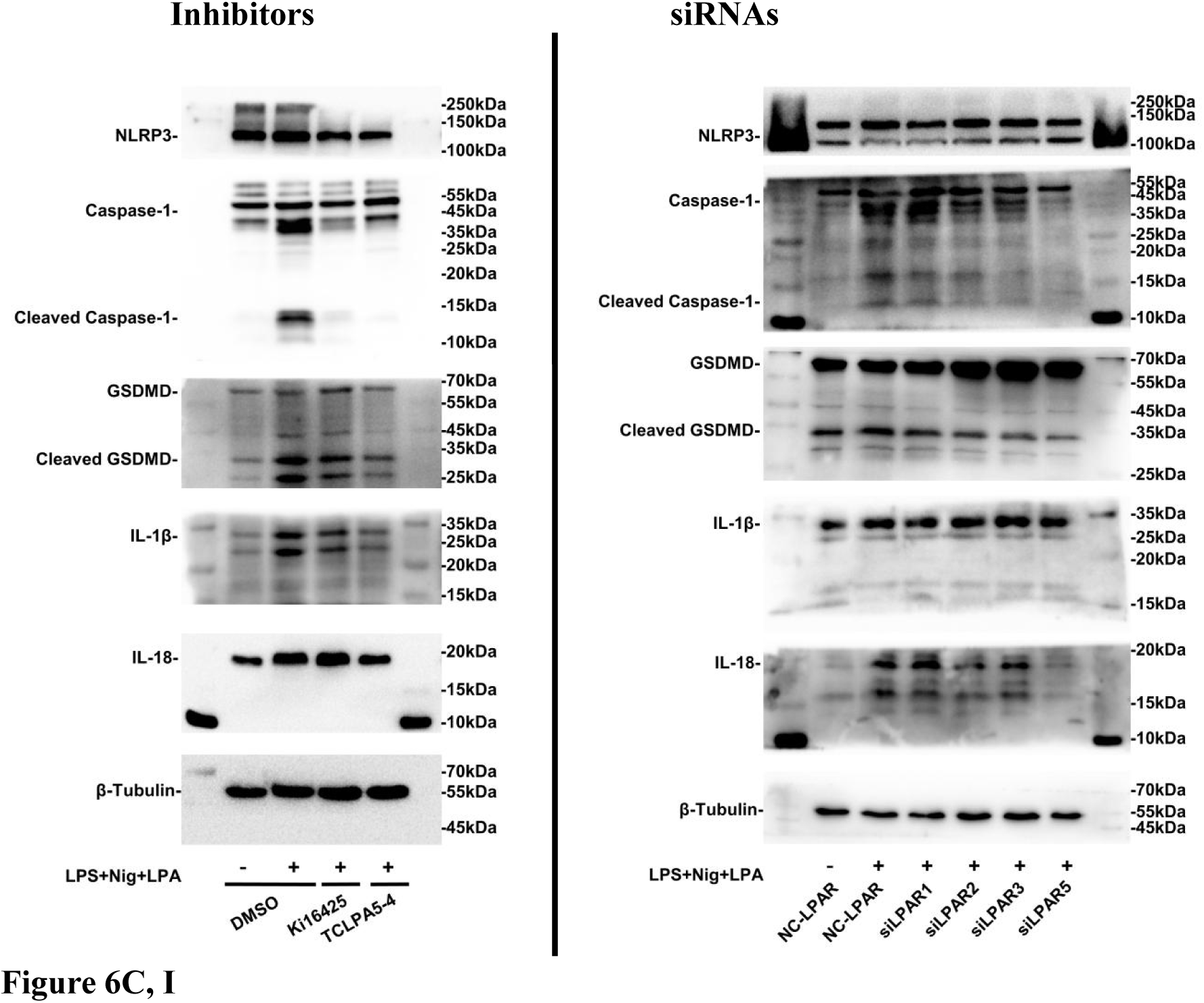

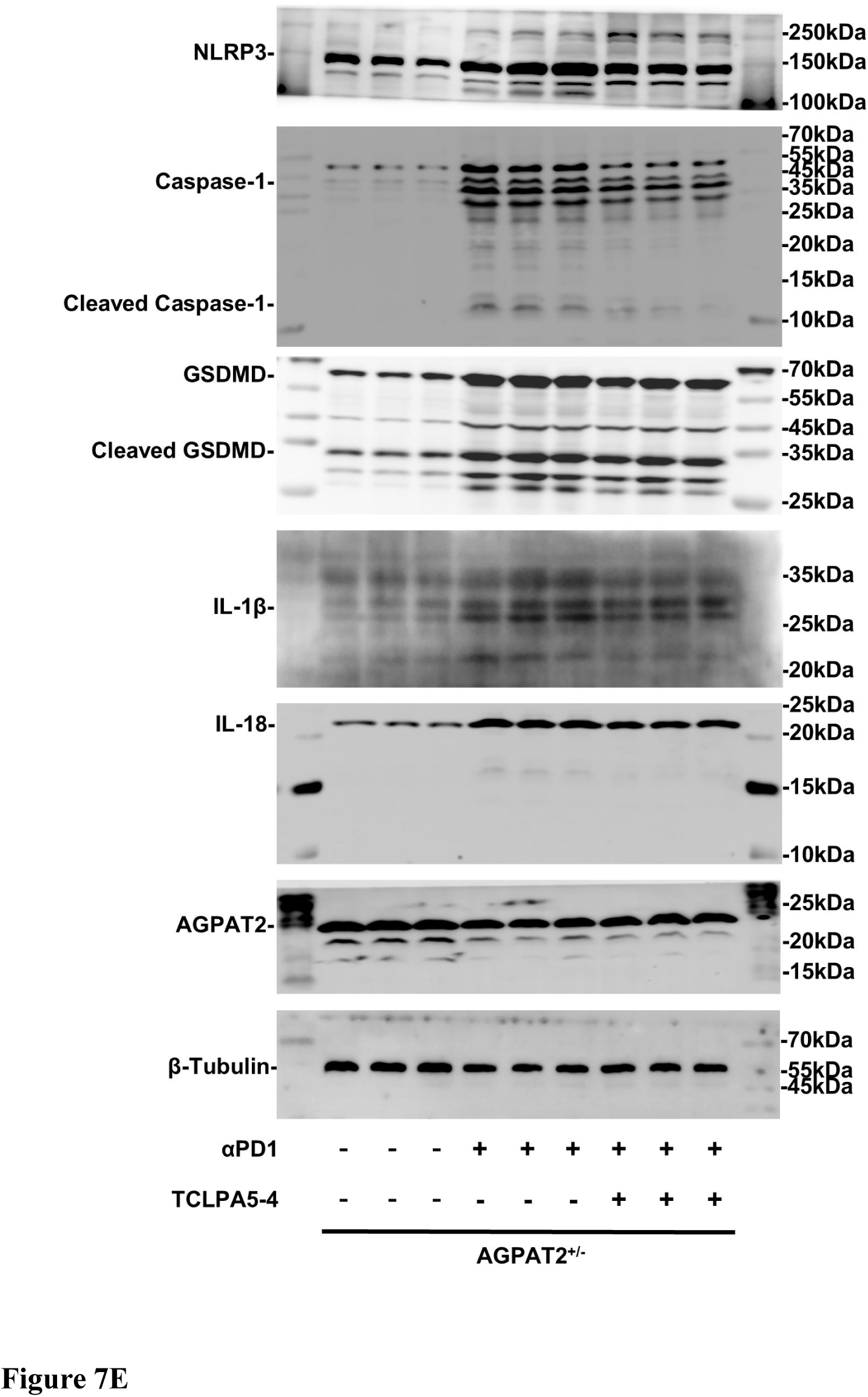

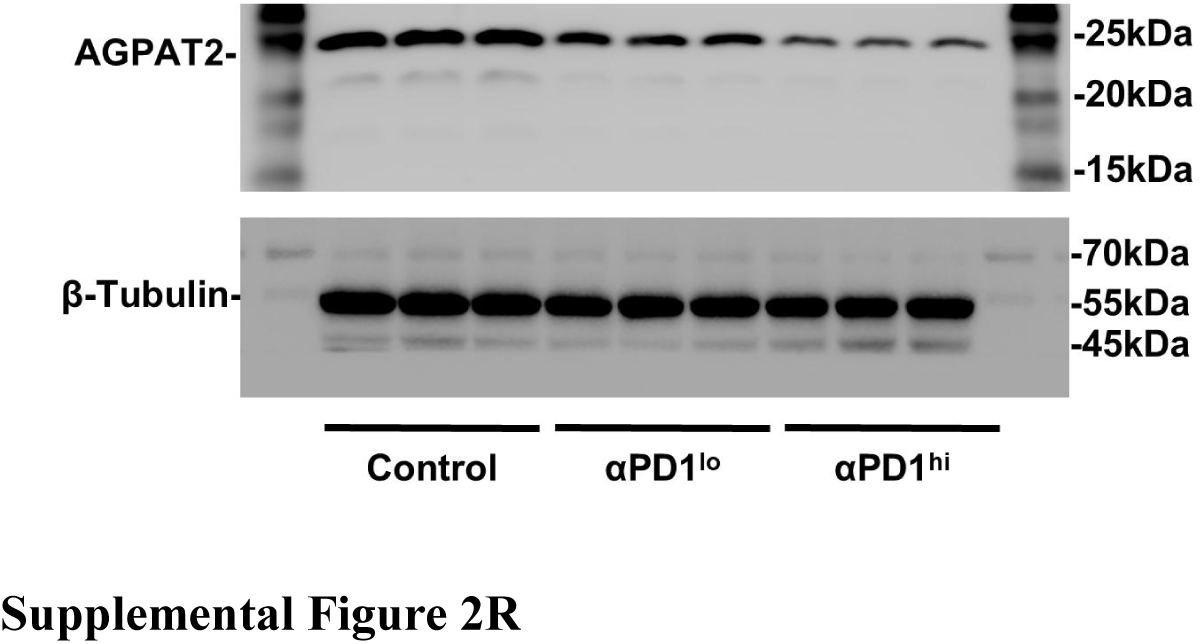

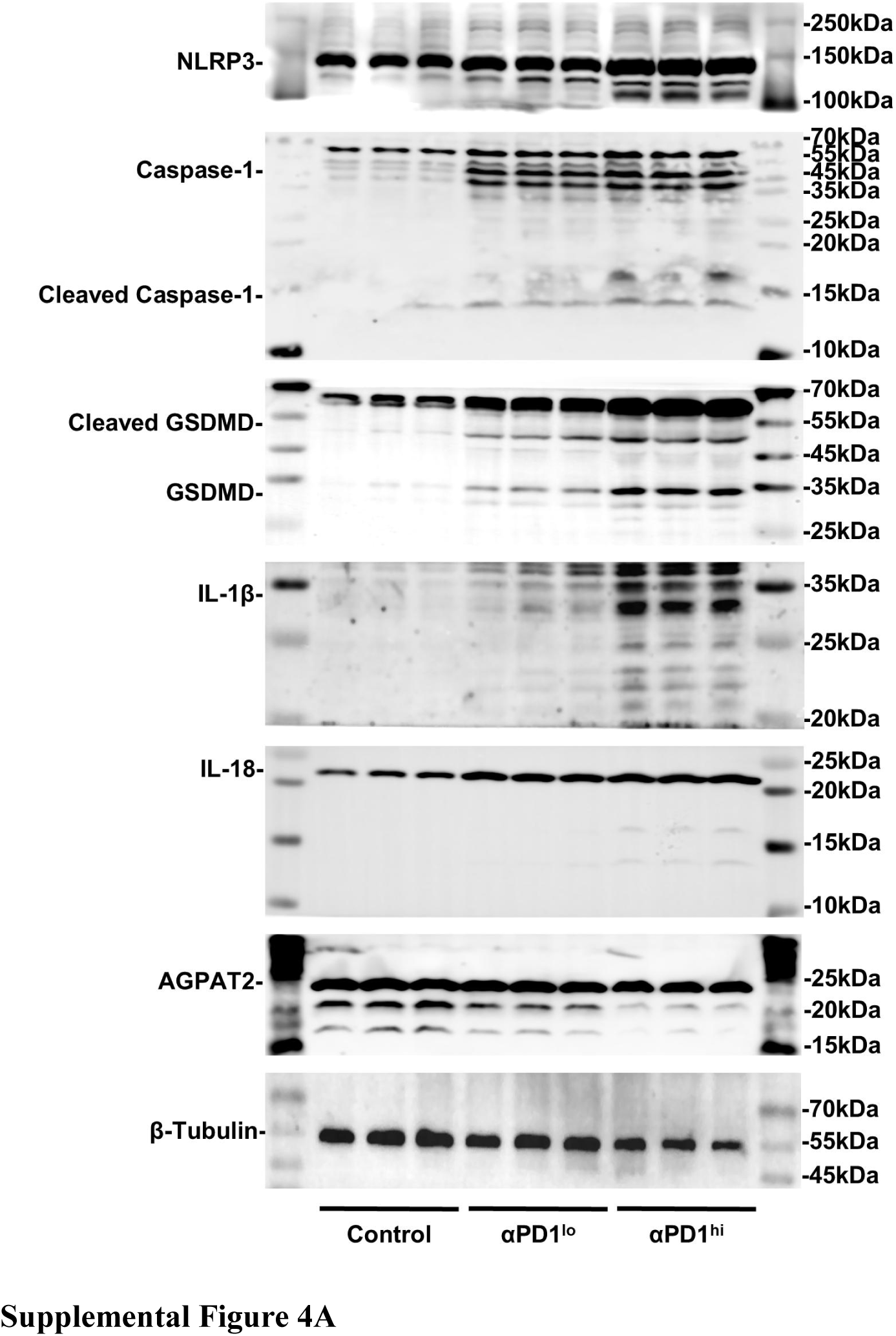

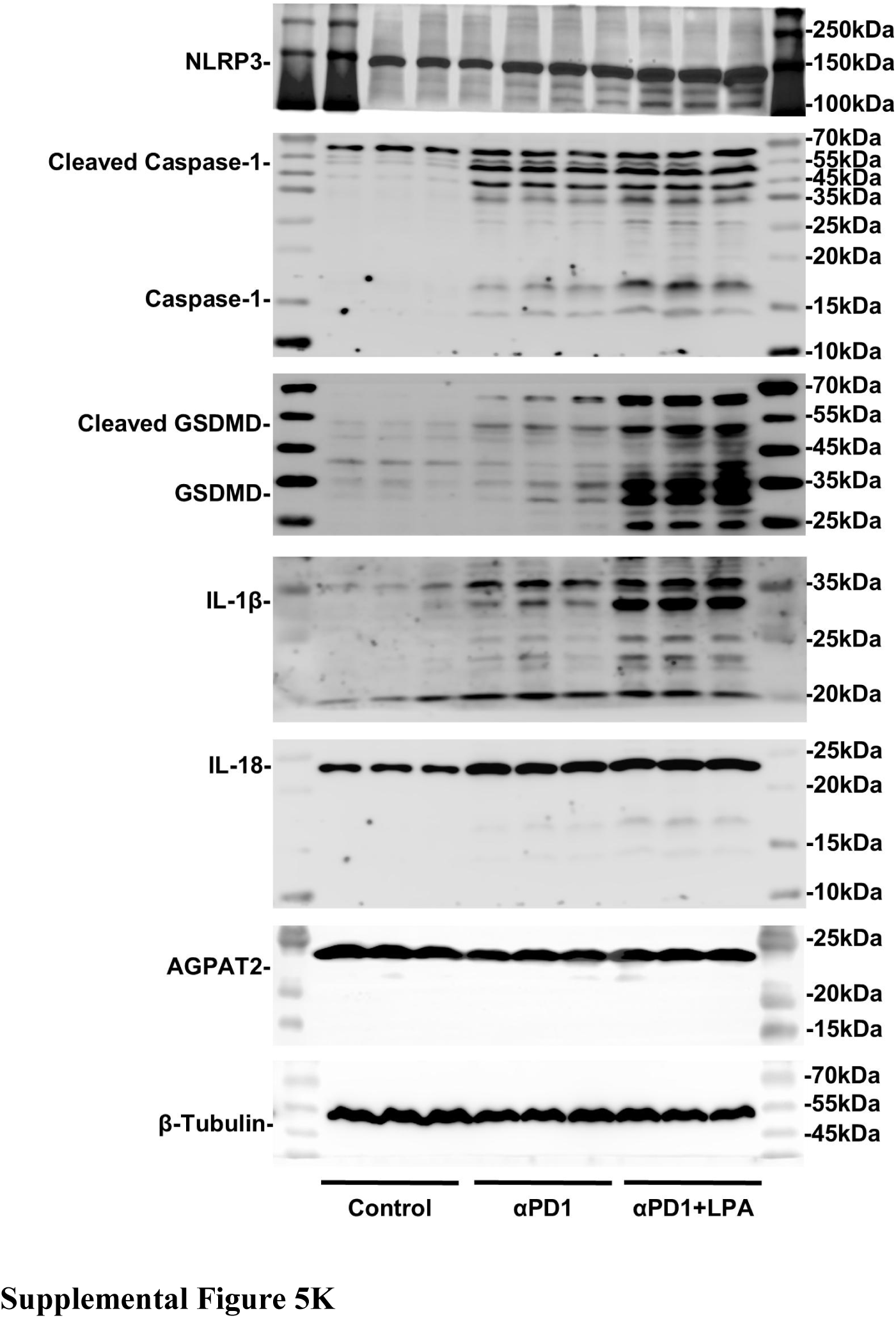

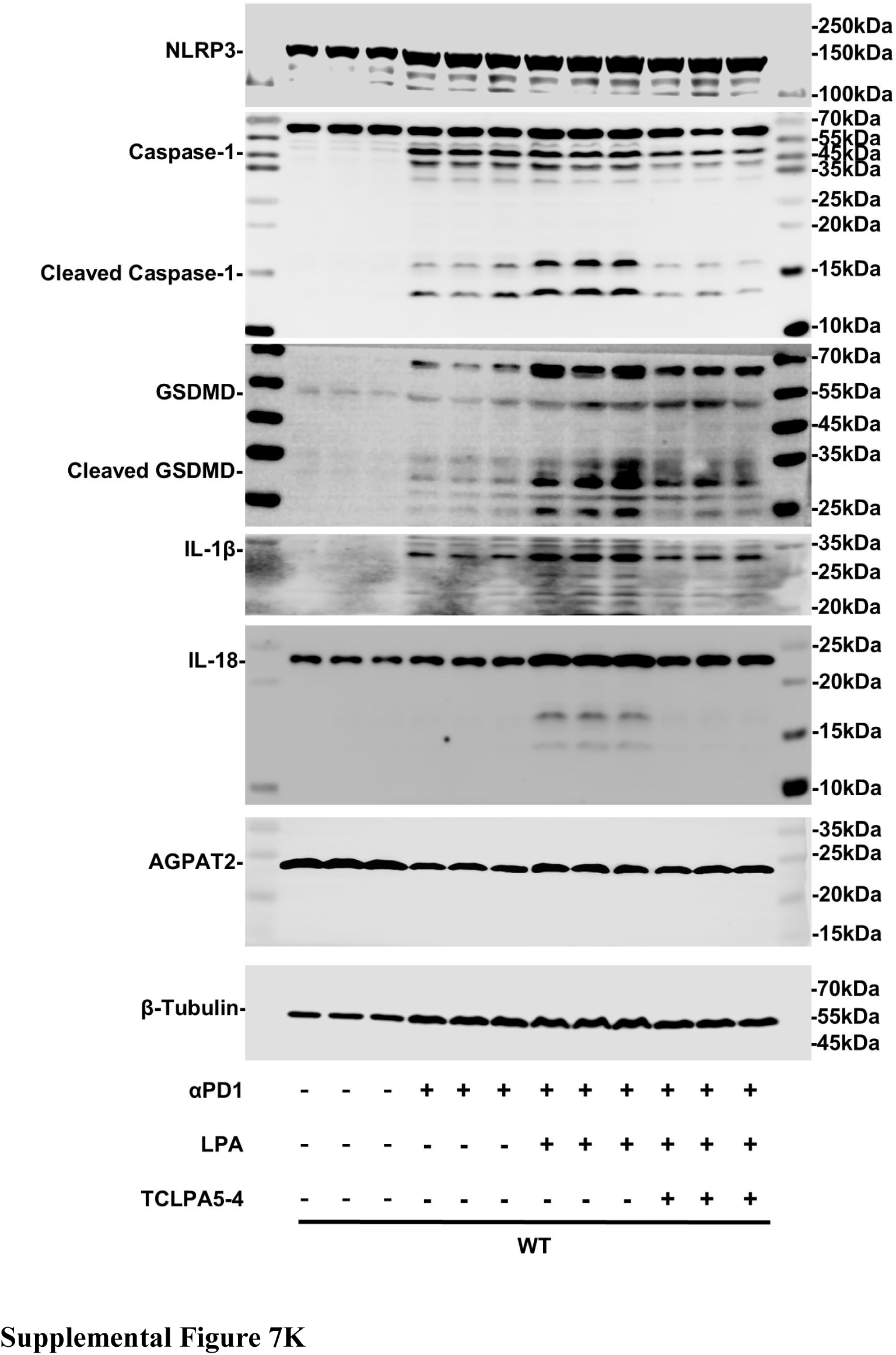

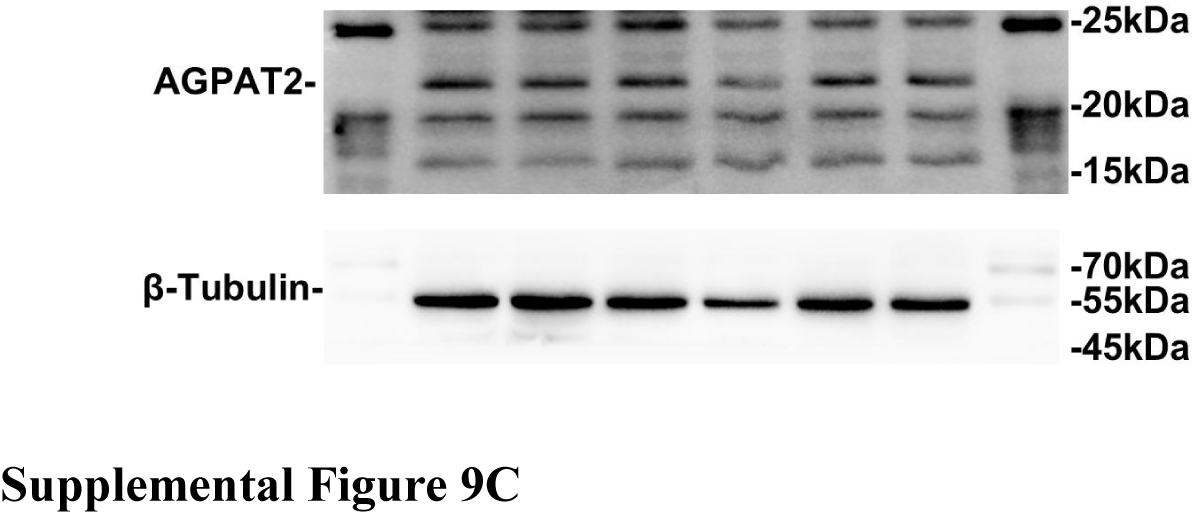

## Notes

### Competing Interest Statement

The authors have declared no competing interest.

